# The results of biodiversity-ecosystem functioning experiments are realistic

**DOI:** 10.1101/725812

**Authors:** Malte Jochum, Markus Fischer, Forest Isbell, Christiane Roscher, Fons van der Plas, Steffen Boch, Gerhard Boenisch, Nina Buchmann, Jane A. Catford, Jeannine Cavender-Bares, Anne Ebeling, Nico Eisenhauer, Gerd Gleixner, Norbert Hölzel, Jens Kattge, Valentin H. Klaus, Till Kleinebecker, Markus Lange, Gaëtane Le Provost, Sebastian T. Meyer, Rafael Molina-Venegas, Liesje Mommer, Yvonne Oelmann, Caterina Penone, Daniel Prati, Peter B. Reich, Abiel Rindisbacher, Deborah Schäfer, Stefan Scheu, Bernhard Schmid, David Tilman, Teja Tscharntke, Anja Vogel, Cameron Wagg, Alexandra Weigelt, Wolfgang W. Weisser, Wolfgang Wilcke, Peter Manning

**Author notes:** Materials and Correspondence: Please address requests to.

## Abstract

A large body of research shows that biodiversity loss can reduce ecosystem functioning, thus providing support for the conservation of biological diversity^1–4^. Much of the evidence for this relationship is drawn from biodiversity-ecosystem functioning experiments (hereafter: biodiversity experiments), in which biodiversity loss is simulated by randomly assembling communities of varying species diversity, and ecosystem functions are measured^5–9^. This random assembly has led some ecologists to question the relevance of biodiversity experiments to real-world ecosystems, where community assembly may often be non-random and influenced by external drivers, such as climate or land-use intensification^10–18^. Despite these repeated criticisms, there has been no comprehensive, quantitative assessment of how experimental and real-world plant communities really differ, and whether these differences invalidate the experimental results. Here, we compare data from two of the largest and longest-running grassland biodiversity experiments globally (Jena Experiment, Germany; BioDIV, USA) to related real-world grassland plant communities in terms of their taxonomic, functional, and phylogenetic diversity and functional-trait composition. We found that plant communities of biodiversity experiments have greater variance in these compositional features than their real-world counterparts, covering almost all of the variation of the real-world communities (82-96%) while also containing community types that are not currently observed in the real world. We then re-analysed a subset of experimental data that included only ecologically-realistic communities, i.e. those comparable to real-world communities. For ten out of twelve biodiversity-ecosystem functioning relationships, biodiversity effects did not differ significantly between the full dataset of biodiversity experiments and the ecologically-realistic subset of experimental communities. This demonstrates that the results of biodiversity experiments are largely insensitive to the inclusion/exclusion of unrealistic communities. By bridging the gap between experimental and real-world studies, these results demonstrate the validity of inferences from biodiversity experiments, a key step in translating their results into specific recommendations for real-world biodiversity management.

## Main Text

Concerns over the consequences of biodiversity loss for human well-being triggered the growth of biodiversity-ecosystem functioning (hereafter: biodiversity-functioning) research, an important field of ecology over the past 25 years^1, 3, 19–21^. Some of the most influential studies in this field are based on biodiversity-ecosystem functioning experiments (hereafter: biodiversity experiments), in which communities of varying diversity are randomly assembled and the responses of ecosystem processes are measured^6, 22^. These experiments, often conducted using grassland communities^8^, aim to isolate the effects of species richness from other factors known to affect ecosystem processes, such as climate, nutrient availability, and the presence of particular plant functional types. By doing so, they have provided strong evidence that biodiversity can affect the functioning of ecosystems – most commonly with a positive log-linear relationship between diversity and plant productivity^1, 2, 5, 7, 21, 23, 24^. However, the relevance of biodiversity experiments to real-world ecosystems (i.e. those where community assembly is influenced by external drivers, such as climate or land-use) has been repeatedly questioned^10–14, 18^. Criticisms highlight several common features of experimental designs, namely random assembly, as opposed to the non-random assembly/disassembly of real-world ecosystems^13^, initial sowing of even species abundances (but see ^25–28)^, and the repeated removal of non-target species (but see ^29, 30^). These factors may alter community assembly processes, leading to unrealistic communities that possess functional properties that are rare or absent in the real world. Although numerous researchers have argued for the relevance of biodiversity experiments^15, 17, 31, 32^ and provided evidence to counter these criticisms^26, 33, 34^, we do not know how closely plant communities in biodiversity experiments resemble those of related real-world ecosystems (but see ^35^ for a local-scale comparison), or if the presence of unrealistic communities affects the conclusions drawn from these experiments. Here we perform a comprehensive, quantitative assessment of the differences and similarities between plant communities from biodiversity experiments and related real-world ecosystems and test the applicability of experimental results to real-world ecosystems.

We quantitatively compared the plant communities of the World’s largest and longest-running grassland biodiversity experiments to those of nearby real-world communities where diversity gradients are created by natural environmental variation and global change drivers. These experiments are the Jena Experiment, established 2002 in Germany (hereafter: Jena Experiment)^6, 30^ and the BioDIV experiment, established 1994 at the Cedar Creek Ecosystem Science Reserve, Minnesota, USA (hereafter: BioDIV)^5, 36–38^ (Fig. 1). We compared experimental communities from the Jena Experiment with those of agricultural grasslands in three regions of Germany, spanning a broad range of site conditions and land-use intensities – the Biodiversity Exploratories^39, 40^ – and semi-natural grasslands close to the Jena Experiment (hereafter: “Jena real world”). BioDIV’s experimental communities were compared to nearby, naturally-assembled prairie-grassland communities at Cedar Creek, including fertilized grasslands^33, 41, 42^ and those undergoing successional change^43^ (see Methods and Supporting Information, Table S1). We combined species-specific cover data from annual vegetation surveys (3,330 and 9,954 plot-year combinations in the German and the US datasets, respectively) with phylogenetic information and plant functional-trait data to characterize and quantitatively compare communities based on a range of properties known to influence ecosystem functioning^44, 45^, including measures of taxonomic diversity and evenness, phylogenetic diversity, functional diversity and community abundance-weighted means (CWM) of selected functional traits of vascular plants, hereafter referred to as “community properties” in a Principal Component Analysis (PCA) (see Methods, Fig. 1).

**Fig. 1.**
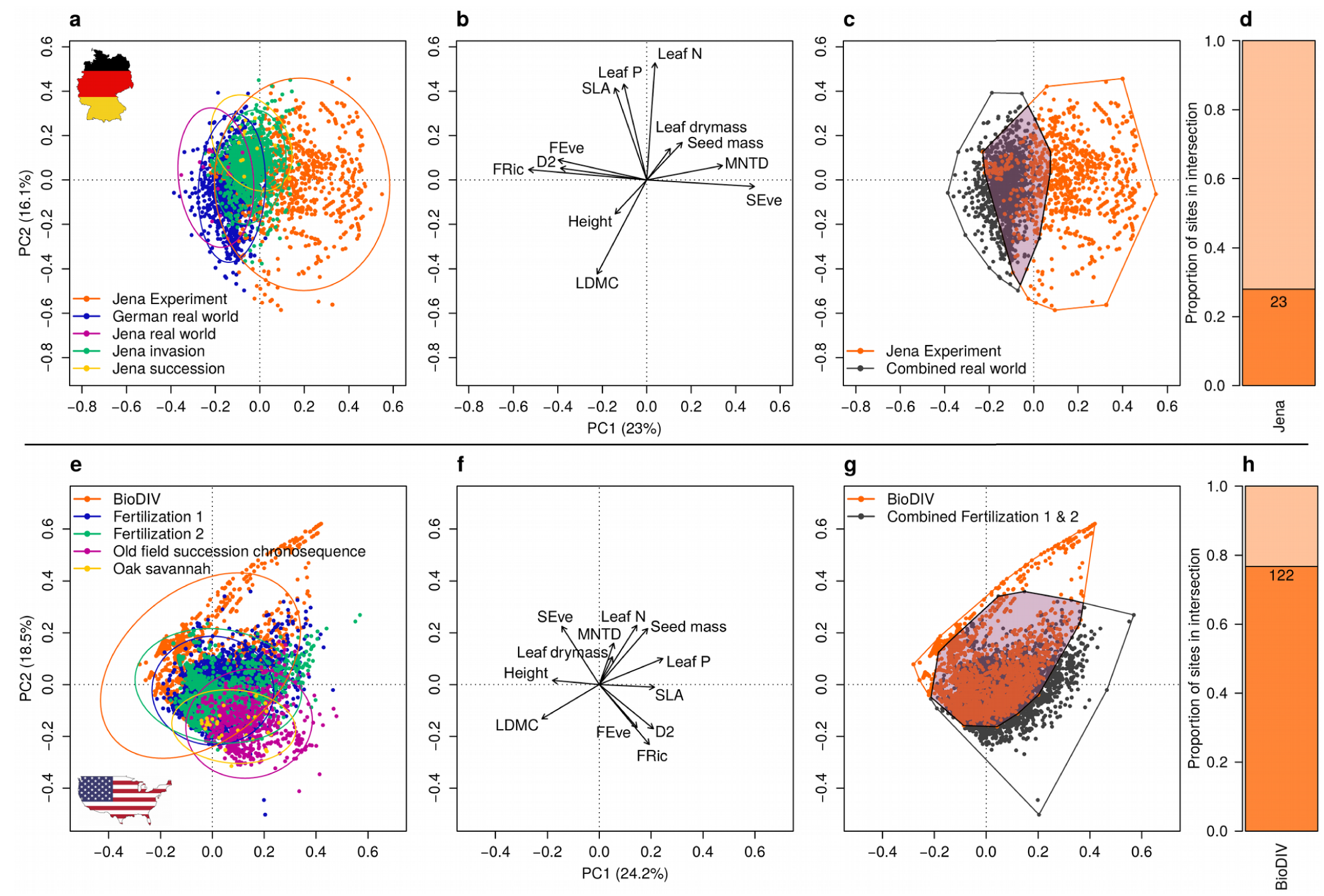
Experimental versus real-world communities. Upper row: German comparison (n=3330 plot-year combinations). Lower row: US comparison (n=9954 plot-year combinations). a-c and e-g: First two axes of a PCA on 12 plant-community properties (see panels b and f, variance-inflation factor-selected community weighted mean traits, functional diversity metrics, phylogenetic diversity and taxonomic diversity metrics), where each dot represents a single plot in a single year. a and e: Distribution of the experimental (orange) and various real-world plots with 95% confidence ellipses (variables scaled for PCA) for each subset. b and f: PCA factor loadings for community properties (arrows proportionally increased to improve visibility; see Supporting Information, Table S13 and S14 for PCA factor loadings and the full dataset, respectively). c and g: Two-dimensional representation of three-dimensional convex hull volumes for experimental (orange) and combined real-world communities (German real world and Jena real-world plots for the German, Fertilization 1 and 2 plots for the US comparison, gray) and their intersection (shaded area). d and h: Number and proportion (strong versus light color) of biodiversity experiment plots in the intersection, where each plot with at least one annual community in the intersection is defined as included.

The results of our comparison showed that experimental plant communities occupy a larger area of multivariate community-property space than real-world communities, despite the latter covering a wide range of climatic, edaphic and management conditions, particularly in the German dataset^39, 46^ (Fig. 1a,e). Furthermore, 82-96% of real-world communities were nested within the space occupied by experimental communities, and additional data collected at Jena showed that experimental communities migrated towards the space occupied by real-world communities when not weeded (Supporting Information Fig. S1). Across both the German and the US datasets, the properties that differed most strongly between experimental and real-world plant communities were mean nearest taxon distance (MNTD), Simpson’s evenness (SEve), and CWM seed mass, which were typically higher in experimental than in real-world communities (Supporting Information, Fig. S2 and S3 and Table S2 and S3). These findings were robust to the inclusion or exclusion of particular community properties (Supporting Information, Fig. S4 and S5b,d and Table S4 and S5).

Overall, three conclusions can be drawn from this comparative analysis: first, biodiversity experiments successfully create plant communities that vary greatly in functionally-important community properties. Second, real-world communities are confined to narrower regions of multivariate community-property space compared with experiments. Third, while the properties of many experimental communities are not observed in related real-world communities, our findings show that a subset of randomly-assembled experimental communities are comparable to real-world communities, (Fig. 1), even though their taxonomic community composition might differ.

In the second step of our analysis, we identified “unrealistic” (i.e., unobserved in the real world) plant communities in biodiversity experiments and tested the sensitivity of biodiversity-ecosystem functioning relationships to the exclusion of these communities. To do this, we identified plots from biodiversity experiments whose communities fell outside the multidimensional community-property space occupied by real-world plant communities (hereafter: “unrealistic communities”). This was achieved by calculating the intersection of three-dimensional convex hull volumes defined by experimental and real-world communities (Fig 1; see Methods for alternative analyses). When using the full set of community properties, 28% and 77% of experimental plots were deemed realistic in Jena and BioDIV, respectively. These realistic biodiversity-experiment communities had significantly higher sown diversity (Jena: av = 21.7 realistic vs. 3.5 unrealistic, BioDIV: 7.8 vs. 1.7; Fig 2) and more sown functional groups (Jena: 2.8 vs. 1.9, BioDIV: 3.5 vs. 1.5), but lower Simpson’s evenness (Jena: 0.5 vs. 0.7, BioDIV: 0.6 vs. 0.9; Fig. 1) than the unrealistic communities. However, realistic and unrealistic experimental communities did not differ in most functional trait CWMs (both Jena Experiment and BioDIV; see Fig. 1, Supporting Information, Fig. S6 and S7, Table S6 and S7).

**Fig. 2.**
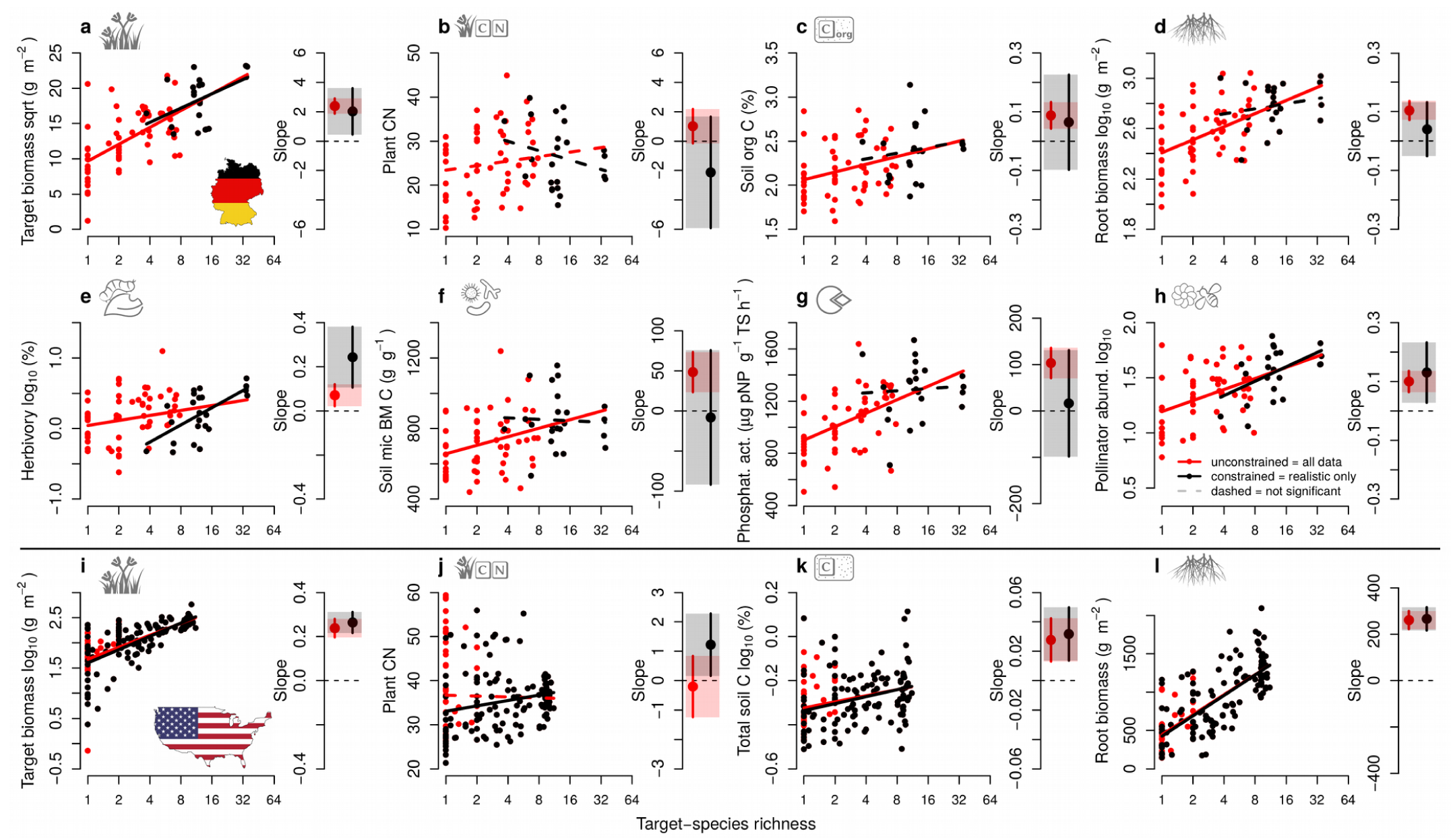
Biodiversity-ecosystem functioning relationships. Relationship between realized target plant species richness (averaged per plot between 2006 and 2015, axis on log_2_-scale) and various ecosystem functions in German (panels a-h, Jena Experiment) and US (panels i-l, BioDIV) biodiversity experiments containing all plots (all dots and red lines) and only realistic plots (black dots and lines). Constrained (realistic plots only) and unconstrained slopes are shown for each panel together with 95% confidence intervals (error bars and shaded areas). For model parameters such as sample sizes, slope estimates, confidence intervals, p-values and adjusted R^2^ values, see Supporting Information, Table S8. Dashed regression lines show non-significant relationships (p>0.05). Note that panels a-d and i-l show the same ecosystem functions for both experiments (organic versus total C in c and k). BM denotes biomass and CN denotes C:N ratios. Where indicated in the y-axis label, data were transformed to meet model assumptions. Response variables were averaged over all available years. Function symbols modified from originals by Hamish, Saeful Muslim, Alice Noir, Lluis Pareras, Creative Stall, Atif Arshad, Made and amantaka from the Noun Project.

Finally, we fitted simple linear models to test how plant species richness affected eight selected ecosystem functions from both the above- and belowground subsystems. These were: plant aboveground and belowground biomass, plant aboveground C:N ratio, soil carbon (C) content, invertebrate leaf herbivory, soil microbial biomass C, phosphatase activity in the soil and pollinator abundance (Fig. 2). This was done for both the full datasets and the subset of realistic plots. In both experiments, and across the different ecosystem functions, the slopes of experimental biodiversity-functioning relationships were remarkably insensitive to the removal of unrealistic communities. A paired t-test of unconstrained versus constrained slopes showed no significant change in slope estimates (t=1.40, df=11, p=0.19) and confidence intervals for slope estimates overlapped each other’s mean for all but two model pairs. The two exceptions to this were both initially weak biodiversity-functioning relationships: Jena-Experiment herbivory, where the positive slope increased when constrained to realistic plots, and BioDIV plant C:N, where a non-significant, slightly negative slope turned into a positive significant one (see Supporting Information Table S8). The finding that the slope of the biodiversity-functioning relationship was largely unaffected by the exclusion of unrealistic communities was robust to changing the set of community properties and the method used to identify realistic communities (Supporting Information, Fig. S8). The goodness of fit (adjusted R² values) was also not strongly affected by constraining the dataset (mean: 0.24 versus 0.15 for unconstrained and constrained models, respectively; Supporting Information, Table S8), and the average percentage change in maximum functioning was ±10.3% (SE: 4%; Supporting Information, Table S9). Together, these, results show that the form, strength, and magnitude of the relationship between biodiversity and functioning that has been identified in biodiversity experiments is generally robust to the removal unrealistic communities.

To check whether the results of the analyses of constrained versus unconstrained communities might have been influenced by the smaller sample size of the former, we assessed the sensitivity of the results to reduced replication. In four out of twelve cases, constraining data led to a change from a significant to a non-significant relationship (Jena soil organic C content, root biomass, soil microbial biomass C and phosphatase activity; Fig. 2). By performing a sensitivity analysis in which we randomly reduced the unconstrained dataset to the same size as the realistically constrained dataset (Supporting Information Fig. S9), we found that the slope of the biodiversity-functioning relationship in the realistic subset for these four relationships was shallower than most random-selection slopes. This suggests for certain ecosystem functions, particularly soil processes, that the slope of the real-world biodiversity-functioning relationship might differ from that observed in biodiversity experiments.

Changes in biodiversity-functioning relationships caused by constraining were partly caused by a reduction in the underlying species richness gradient in Jena. Here, minimum species richness changed from 1 to 3.7 between unconstrained and constrained plots. For BioDIV, which covers a relatively narrower range of species richness, the gradient was not reduced (Fig. 2 and Supporting Information, Table S10). The shorter species richness gradient was associated with a reduction in the range of functioning covered across the reduced biodiversity gradient. Overall, reductions were 31% in Jena and 7% in BioDIV (Supporting Information, Table S10). As such, the low diversity levels in the Jena Experiment, although needed for experimental design that can identify diversity effects and their underlying mechanisms^47^, are generally found to be unrealistic when compared to current German real-world communities.

Our results show that the biodiversity-functioning relationships observed in biodiversity experiments are not an experimental artefact caused by the presence of unrealistic communities, and that the mechanisms observed in these experiments are also likely to operate in real-world communities^5, 9, 30, 32, 47^. The question remains, however, how important biodiversity-functioning relationships are as drivers of ecosystem functioning in the real world relative to factors such as land use or climate^7, 14, 48^. Although strong and positive biodiversity-functioning relationships have been reported in real-world studies^4, 22, 34, 49–51^, other studies describe weak or negative relationships^4, 52, 53^. This inconsistency, and the discrepancy between experimental and real-world patterns, is commonly attributed to the presence of covarying environmental or biological factors that also drive ecosystem functioning^54^, and which obscure, confound or negate the effects of biodiversity (e.g., nutrient availability, climate, and the dominant functional traits of the community;^50, 55–57)^. These factors are likely to be closely coupled in real-world ecosystems, but decoupled in experiments. Indeed, across our datasets, the average correlation strength of the eight measures of dominant functional traits (CWM’s) with Simpson’s evenness, functional, and phylogenetic diversity properties was higher in real-world than in experimental data subsets; mean absolute correlation coefficients were 0.18 and 0.22 in German and American real-world sites, compared to 0.08 and 0.16 in their respective experiments (Supporting Information, Table S11 and S12).

While the biodiversity experiments used in our analysis cover a wide range of plant-community properties, only a fraction of this multidimensional space is occupied by related real-world communities. The remainder of space covered by the experimental communities is currently not observed in the real-world communities that we considered; however, this “unrealized plant community property space” may be useful in predicting ecosystem functioning in the future, when novel combinations of species and environmental conditions may emerge^31, 58^.

In conclusion, we show that although biodiversity experiments deliberately include plant communities that may not currently occur under real-world conditions, excluding those communities does not result in significantly-altered biodiversity-functioning relationships in most cases. Our results complement previous reports of significant biodiversity-functioning relationships in the real world^4, 34, 40, 49, 51, 55^ by showing that constraining experimental datasets to contain only realistic plant communities does not change the core conclusions of biodiversity-functioning research. To advance this field, we must acknowledge both the strengths and limitations of biodiversity experiments. Specifically, our improved understanding should be used to develop a new generation of experiments, e.g. that focus on realistic patterns of community change. At the same time, we must maintain and further examine the valuable resource of long-term biodiversity experiments, e.g. by re-analyzing existing experimental data to simulate a range of possible biodiversity-change scenarios. By moving beyond critiques of experimental design and placing experimental biodiversity-functioning research in the context of natural communities, we advance the current debate from verbal arguments to a quantitative investigation, thus increasing the robustness and applicability of biodiversity-functioning research.

## Material and Methods

### 1. Overview and data origin

We chose the largest (Jena) and longest-running (BioDIV) grassland biodiversity experiments in the world for our comparison. The Jena Experiment^6, 30^ was chosen as a Central-European example of a long-term, intensively studied biodiversity experiment^30, 59^. In the Jena “main” experiment, combinations of 1, 2, 4, 8, 16 and 60 species from a pool of 60 Arrhenatherion grassland species^60^ were sown in 82 originally 20 m × 20 m plots on a former agricultural field in 2002. This species richness gradient was crossed with a gradient of functional group richness (1 to 4 functional groups; small herbs, tall herbs, grasses, legumes), where species were randomly chosen from the respective functional groups^6^. Jena Experiment plots are maintained by weeding (two or three times a year). All plots are mown twice per year and mown biomass is removed, a common management of meadows in the region, and do not receive any fertilizers. The Jena Experiment includes two invasion sub-experiments; one set of Jena “invasion” plots was not weeded after initial sowing and studied regularly until 2009, another set was weeded initially, but weeding was stopped in 2010; here, we use the former for 2003–2009 and the latter for 2010–2015. Jena mown “succession” plots were not initially sown and are excluded from all management except for the mowing.

As a real-world counterpart to the Jena Experiment, we chose the grassland plots of the Biodiversity Exploratories project (hereafter: “German real world”). This large-scale, long-term research project was established in 2006 to assess the effects of land-use intensity on biodiversity and ecosystem functioning in three regions of Germany^39^. The 150 grassland plots measure 50 m × 50 m and were selected to cover a wide and representative range of land-use intensities, here composed of varying levels of mowing frequency, grazing intensity and fertilization^61^. Species richness in Exploratories grasslands ranged from nine to 70 species, within a 4 m × 4 m subplot, across all years used in our study. Exploratories data were augmented by the inclusion of data from 14 semi-natural grassland plots in the Saale river valley near the Jena Experiment (unpublished data; hereafter: “Jena real-world”). These plots are usually mown twice per year; most are unfertilized and some are moderately fertilized.

The Cedar Creek biodiversity experiment e120 (hereafter: “BioDIV”; ^5, 36, 37, 62^) was selected as a North-American example of a long-term biodiversity experiment, along with a suite of other naturally-assembled grasslands at Cedar Creek that served as nearby real-world communities. BioDIV was established in 1994, when 1, 2, 4, 8 or 16 species were randomly drawn from an 18-species pool and sown across 168 13 m × 13 m plots at the Cedar Creek Ecosystem Science Reserve in Minnesota, USA.

Several datasets of local experiments and observation plots served as local real-world comparison for BioDIV. Experiments e001 (hereafter: “Fertilization 1”) and e002 (hereafter: “Fertilization 2”) were set up in 1982 to study the long-term effects of fertilization with nitrogen and other nutrients, ranging from low rates of nutrient inputs that are similar to atmospheric N deposition rates to high rates of fertilization similar to that used in agriculture. They consist of 324 plots located across three successional grassland fields (324 plots = 2 fertilization experiments × 3 old fields × 9 fertilization treatments × 6 replicates) that differ in their age since abandonment from agriculture and 45 plots in one never-plowed oak savannah in Fertilization 1 (45 plots = 9 nutrient treatments × 5 replicates)^41^. Plot sizes were 4 m × 4 m in the younger fields and 2 m × 4 m in the oak savannah. In contrast to Fertilization 1, Fertilization 2 plots were agriculturally disked before receiving nutrient addition treatments. Plot-level species richness in the two fertilization studies ranged from one to 28 species across all years used in our study. Established in 1983 and 1989, the Cedar Creek project e014 (hereafter “Old field succession chronosequence”) offers vegetation data from four to six observational transects in each of 23 different fields repeated seven times between 1983 and 2011 to study succession after agricultural abandonment^43^. Cedar Creek project e093 (hereafter: “Oak savannah”), established in 1991, offers data from 30 2 m × 2 m prairie opening plots of natural vegetation^63, 64^. This combination of Cedar Creek datasets was chosen to represent a variety of real-world plant communities that were comparable to the BioDIV experiment. Note that while Central European grasslands depend on anthropogenic management (mowing, grazing) to prevent succession to forest, the US prairies are naturally fire-disturbed, hence the selection of agricultural sites as the German real-world grassland. Please note that while all above-described datasets were used in the multivariate analysis of plant community property overlap (Fig. 1a,b,e,f), only a subset was used in constraining the biodiversity experiment data to realistic sites (Fig. 1c,d,g,h; see below). For an overview of the datasets used in this study and online resources to obtain the original data, see Table S1 in Supporting Information.

### 2. Plant-community properties

#### Vascular plant cover and biomass

In the Jena Experiment, vegetation surveys were performed annually in the second half of May on a 3 m × 3 m subplot of each plot and species-specific cover data was collected. Note that, in the Jena “main” plots, only target species (vascular plants originally sown in the respective plots) were recorded. Vegetation surveys of the invasion and succession plots were performed annually in 2 m × 2.25 m subplots (2003-2009) or 3 m x 3 m subplots (2010-2015), assessing all present species. We used Jena vegetation data from 2003–2015 (succession data only from 2003–2009). In the Biodiversity Exploratories (German real-world plots), species-specific vascular plant cover was estimated annually in a 4 m × 4 m subplot of each plot between Mid-May and Mid-June. Here, we used all data from 2008-2015. Data from the 3 m × 3 m vegetation surveys of Jena real-world plots was available for May 2011. To test if the different vegetation survey areas in Jena and the Biodiversity Exploratories might bias the relative abundance of vascular plant species and thus the calculation of abundance-weighted community properties, a separate survey of 27 Biodiversity Exploratories plots was performed by sampling species-specific cover in series of nested 4 m × 4 m, 3 m × 3 m and 2 m × 2 m subplots. As cover estimates did not show any sign of systematic variation (Supporting Information, Fig. S10), we concluded that the different survey areas were unlikely to bias our results.

For BioDIV, a combination of species-specific cover data (1996–2000) and species-specific aboveground peak biomass (2001–2015) data was used to calculate plant community relative abundance. Previous analyses have shown that this difference in methodology does not affect the conclusions of analyses investigating species-richness effects on biomass^65^. Cover estimates for BioDIV were obtained by averaging the estimates from four permanently-marked subplots (each 0.5 m × 1 m) within each plot. Species-specific biomass in BioDIV was obtained by annually clipping 0.1 m × 6 m strips on each plot, drying and sorting the resulting biomass to species.

For Fertilization 1 and Fertilization 2, species-specific plant aboveground biomass data was collected annually at peak biomass by clipping a 0.1 m × 3 m strip of vegetation per plot, sorting and drying it. Years 1982–2004 were used for Fertilization 1 and 1982–1991 for Fertilization 2 as these years maintained the original, balanced treatment design, which was later changed to add further treatments. For the old field succession chronosequence plots, species-specific cover values were used for seven years between 1983 and 2011. Each of the 23 fields had four transects (except for two fields with six transects) of 25 subplots each. For comparability to the other datasets, the 25 transect subplots of 0.5 m × 1 m in each transect were treated as one plot by averaging species-specific cover values across the subplots within transects resulting in four (or six) plots for each of the 23 fields (96 plots=21 fields × 4 plots + 2 fields × 6 plots). For the oak savannah dataset, only plant species cover from 1991 was used; later years were excluded because they were affected by a seed addition treatment. Species-specific cover was averaged across the 16 0.5 m × 0.5 m subplots per plot.

For comparative analyses, different years were chosen for these different datasets due to varying availability of measurements and in order to choose years with consistently balanced design of the experimental treatments in cases where treatments were added after the onset of the experiments. The transects in the old field succession chronosequence are likely to inflate certain community properties because their subplots span out further across the respective sites than a square plot of the same area would. Similarly, the averaging across subplots in the oak savannah dataset might influence the direct comparability to the biodiversity experiment data. As such, the data from the old field succession chronosequence and the oak savannah dataset are shown in Fig. 1e to put the BioDIV data into perspective by adding different kinds of real-world data. However, when it came to constraining biodiversity experiment data with the real-world data (Fig. 1g), we took a conservative approach and included only those real-world datasets that were most comparable in terms of plot types (Fertilizer 1 and 2; hereafter: Combined US real world). Similarly, for the Jena Experiment real-world counterparts, we considered only the German real world and Jena real-world sites as purely non-biodiversity experiment sites in Fig. 1c (hereafter: Combined German real world).

To enable direct comparisons of plant communities, species-specific cover and biomass values for all projects were transformed to relative abundance where the single abundance values within each community sum to 100. In order to do this, all Jena Experiment cover values (originally estimated on a decimal scale, ^66^) were first transformed to percent cover values^67^. Bare ground was ignored, so where vegetation covered <100% of the plot, it was scaled to 100% for the calculation of community properties.

#### Species synonyms and phylogeny

As we used plant species cover, biomass, and trait data from multiple sources based on research across decades and different geographic regions, there was considerable variation in the classification and nomenclature of species. Additionally, since the TRY database^68^ was queried for plant traits and we also used a phylogenetic backbone tree (see below), the various datasets contained species names that might not all be currently accepted names, challenging the linkage of the different datasets. This issue was dealt with by creating “code” data frames connecting all original spellings, outdated and synonym names to the names for which data was available and to the accepted species names obtained using The Plant List via function “TPL” in R package “Taxonstand”^69^.

To calculate phylogenetic diversity metrics and to use phylogenetic relatedness to assist the imputation of missing trait data, a phylogenetic tree of all plant species was created and included in our study. We adopted the nomenclatural criteria in The Plant List v. 1.1^70^ for the species in our dataset, and pruned the updated vascular plant megaphylogeny by Qian & Jin^71^ to include only the species in our study (n = 664). We used the software SUNPLIN^72^ to add the species lacking from the megaphylogeny (n=132 or 19.9% of all species in our study) at random within the crown nodes of the corresponding monophyletic genera. In a few cases where the genera of the missing species were polyphyletic (*Potentilla, Medicago, Solidago, Galium*) or paraphyletic (*Calamagrostis, Vicia*), we inserted the species at random within the nodes representing the most recent common ancestors that unequivocally contain them (see ^73^). We repeated this procedure iteratively to obtain 50 phylogenetic trees (see Supporting Information, Fig. S11 for one example tree and the distribution of randomly inserted species). When using the phylogenetic trees in the subsequent data analysis (calculation of phylogenetic diversity metrics and plant trait imputation), all 50 trees were used and results were averaged.

#### Functional trait data

In order to calculate community weighted mean trait values for all plant communities, functional trait data from the TRY database (see Supporting Information, Table S15) were complemented with in-situ collected trait data from Cedar Creek and not published in TRY. Plant species specific functional trait values were calculated separately for the German and US species subsets.

Trait data for leaf area (mm²), leaf dry mass (mg), leaf dry matter content (LDMC, g/g), leaf nitrogen concentration (leaf N, mg/g), leaf phosphorus concentration (leaf P, mg/g), plant height (m), specific leaf area (SLA, mm²/mg) and seed mass (dry mass in mg) were assembled (Cornelissen et al. 2003). These traits were selected as they are important for ecosystem functioning^44, 45^ and data for them was available. For the details of processing TRY and other trait data to generate species-level values, see Supporting Methods.

To fill gaps in trait data, trait values from same-genus species with available trait information were inferred. Subsequently, the “phylopars” function in the R package “Rphylopars”^74^ was employed to impute missing data based on available information on other traits and the phylogenetic tree^75^. Before imputation, all trait data was natural-log transformed. To account for phylogenetic uncertainty (see above), trait data for all 50 phylogenetic trees was imputed and averaged. Subsequently, the plant species and their trait values were visualized in a PCA for each region (Supporting Information, Fig. S12) to check for strong outliers and check the outlier-species’ ability to score extreme values.

#### Calculation of plant-community properties

Before calculating plant-community properties, tree species, occurred as seedlings, were removed from all datasets, because of their strong impact on the calculated CWM’s and functional metrics, and the fact that biodiversity experiments are mown annually thus preventing tree invasion. Plant-community properties were calculated for each plot-year combination so that the temporal development (succession) of plots was accounted for in our analysis. As taxonomic diversity indices, we calculated species richness (S), Shannon’s diversity (H), Simpson’s diversity index (D1), and inverse Simpson’s diversity index (D2) (calculated as D1=1-D and D2=1/D, where D is the sum over all pi^2 and pi are the relative abundances of all species i) with functions “specnumber” and “diversity” in R package “vegan”^76^ and Simpson’s evenness (SEve, by dividing D2 by S)^77–80^. As phylogenetic diversity indices, we used Faith’s phylogenetic diversity (PD), mean pairwise distance (MPD), and mean nearest taxon distance (MNTD)^81^ with functions “pd”, “mpd” and “mntd” in R package “picante”^82^, where MPD and MNTD were calculated with abundance-weighting. All three phylogenetic diversity properties were calculated for each of the 50 phylogenetic trees and averaged to account for phylogenetic uncertainty (see above). For the calculation of the functional diversity indices functional richness (FRic), functional evenness (FEve), functional divergence (FDiv), functional dispersion (FDis), Rao’s quadratic entropy (RaoQ)^83–85^ and community weighted mean traits (CWM’s) the function “dbFD” in the R package “FD”^84, 86^ was used. As function “dbFD” relies on the computation of a Gower dissimilarity matrix where zero-dissimilarity values between two species (identical trait values) are not allowed, we slightly altered the trait values of a small number of species by deliberately increasing all trait values by 0.001 to 0.002% for the function to run. For each of the respective species pairs, only the species with the lower overall cover (throughout the regional dataset) received this alteration (Supporting Information, Table S16). For all but FRic, the abundance-weighted versions of these indices were computed. Communities comprising less than three species were assigned a value of zero for FRic, FEve, FDiv, PD, MPD and MNTD, as their computation is not possible for such communities.

### 3. Multivariate analysis of experiment and real-world intersection

#### Multivariate comparison

All analyses were carried out in R version 3.4.2^87^. Here, a multivariate PCA approach was employed, based on a number of plant-community properties to assess the distribution, similarities and differences between plant communities of biodiversity experiments and real-world systems. Prior to the analysis, we tested for multicollinearity of community properties by calculating variance inflation factors (hereafter: vif; R function “corvif” provided by ^88^). In the German and US dataset, we sequentially removed the variables with the highest variance inflation factor until all vif values were <3. Only the last of the eight variables to remove differed between the German and US datasets, so for comparability between regional datasets, we removed all nine variables from both datasets (see Supporting Information, Table S17 and S18). Specifically, H, FDis, S, leaf area, D1, PD, MPD, RaoQ and FDiv were removed (in order of sequential removal) and only the following 12 community properties were employed in the PCA’s: D2, SEve, FRic, FEve, SLA, leaf dry mass, leaf N, leaf P, seed mass, height, LDMC, and MNTD (Fig. 1b and f). Separate community property PCA’s were computed for the German and USA data subsets using the “rda” function in R package “vegan” (with variables scaled to avoid bias due to different range-size of properties) and the data was visualized in biplots with 95% confidence ellipses (Fig. 1a and e).

#### Intersection-calculation methods

The intersection between experimental and real-world plots was calculated using three different methods of differing complexity, all based on the community-property PCA’s presented in Fig. 1a and e. Intersections were calculated between two groups of data per geographic region: a) all experimental communities across all years and b) a subset of the most comparable and data-rich real-world datasets (combined real-world datasets). For Jena, the related combined real-world communities were all German real-world communities (Biodiversity Exploratories) and the Jena real-world communities. For BioDIV, only Fertilization 1 and Fertilization 2 plots were used as the combined real-world counterparts when calculating the intersections as different vegetation-survey techniques in the old field succession chronosequence and the oak savannah datasets (transects and subplots) made these data incomparable. First, the first two PCA axes were used to assess the two-dimensional intersection of 95% confidence ellipses for experimental and real-world data using the functions “ellipse” and “point.in.polygon” in R packages “car”^89^ and “sp”^90, 91^, respectively (Supporting Information, Fig. S4). Second, the first three PCA axes were employed to compute the intersection of three-dimensional convex hull volumes using functions “convhulln” and “tsearchn” in R package “geometry”^92^ (Fig. 1c and g show 2-dimensional representation of 3-dimensional convex hull volume). Third, using the first three PCA axes, three-dimensional hypervolumes were computed using the “hypervolume” package in R^93^. The intersection hypervolume of the experimental and real-world hypervolumes was then calculated and function “hypervolume_inclusion_test” was used to assess which communities fall in the intersection hypervolume (Supporting Information, Fig. S4). For the subsequent analysis of diversity-functioning (hereafter: BEF) relationships, experimental plots were defined as realistic if their plant communities fell inside the intersection in at least one of the years present in the dataset. Given this threshold, each plot in the experiments was either defined as realistic (the plot’s plant community was within the intersection in at least one year) or unrealistic. Calculating the intersection based on three different methods of different complexity demonstrated that the selection of realistic communities was largely insensitive to the underlying methodology (Supporting Information, Table S4 and S5, Fig. S5a, c). Therefore, we focus our analyses on using three-dimensional convex-hull volumes, a method of intermediate complexity, and present results for the other methods in the Supporting Information.

### 4. Measurement of ecosystem-function variables

A range of above- and belowground ecosystem process rates and state variables was selected as ecosystem functions from the Jena Experiment and BioDIV in such a way that the functions of these experiments were as comparable as possible. Only function data obtained between 2006 and 2015 (at least 4 years after initiation of the experiments) was used because BEF relationships shortly after the initial establishment of experiments are often unrepresentative of longer-term trends^24, 94^. These selection criteria resulted in the following functions: Plant aboveground biomass (biomass), aboveground plant biomass C:N ratio (plant C:N), soil carbon (C; only organic fraction in Jena, total soil C in BioDIV) and root biomass were available for both experiments. As inorganic C is a significant proportion of total soil C at Jena, but not at Cedar Creek, soil organic C was used for Jena, but total soil C for BioDIV. Herbivory rate, soil microbial biomass C, phosphatase activity, and pollinator abundance were only available for Jena. For details regarding the measurement of these ecosystem functions in the Jena Experiment and BioDIV; please refer to the Supporting Methods section.

### 5. Statistical analysis of unconstrained and constrained experimental BEF relationships

In order to assess whether – and how much – BEF relationships change when excluding unrealistic plots from the analysis, each relationship was first analyzed in the unconstrained dataset with all experimental plots. Subsequently, biodiversity experiment datasets were constrained to only include realistic plots and the models were re-run. For ecosystem function variables with multiple years of data, values were averaged across years and simple linear models were fit that tested for the effect of realized target species richness (log_2_, averaged per plot between 2006 and 2015) on the individual functions. Where necessary, square-root or log_10_-transformation was applied to response variables to meet model assumptions of normality and homoscedasticity of variances. For each of the resulting relationships, slope estimates and their 95% confidence intervals (function “confint” in R) were calculated. Slopes and confidence intervals of each pair of constrained and unconstrained relationships were compared to decide if the slope or sign of the relationship had changed. If confidence intervals of unconstrained and constrained slopes included each other’s mean value, we concluded that they were not significantly different. Additionally, a paired t-test on differences between unconstrained and constrained slopes was performed.

### 6. Sensitivity analyses

Since our analysis involved many decisions on which variables to include and what exact analytical pathway to follow, and because we are aware that these decisions might affect our results, several sensitivity analyses were performed regarding different aspects of our analysis.

To test if different subsets of community properties entering the PCA affected our results, our analysis was re-run for all combinations of i) different subsets of community properties, i.e. (a) all 12 community properties (presented in the main text), b) just the eight CWM’s, or c) just the four functional diversity properties) and ii) all three methods to compute the intersection between experiment and real-world plots described above (Supporting Information, Fig. S4 and S8).

To test if shifts in significance of BEF relationships in Fig. 2 simply resulted from the strong reduction of error degrees of freedom associated with using data subsets, we performed a sensitivity analysis randomly selecting the same proportion of plots as realistic that was selected by our PCA-driven selection of realistic sites, 500 times for each relationship (Supporting Information, Fig. S9).

To gain further insight into our findings at Jena, data from experimental plots which were abandoned and allowed to undergo natural succession (Jena invasion plots) was more closely analyzed. Over time, these migrated towards the multivariate community property space occupied by real-world communities, thus showing that differences between real-world and biodiversity experiment communities were due to experimental manipulation and maintenance rather than differences in plot conditions, species pools or initially random versus natural community assembly (Supporting Information, Fig. S1).

## Acknowledgements

We thank the establishers, maintainers, coordinators, technical and research staff of all involved projects, data owners of all involved research projects, the TRY initiative and data owners. Santiago Soliveres, Eric Allan for discussion, Sven Thiel and Guangjuan Luo (both Jena) and Dan Bahauddin (Cedar Creek) for help with data extraction and handling of project databases, Florian Schneider for help with processing TRY data. Thanks to Robert Junker and Benjamin Blonder for assistance with the calculation of multidimensional hypervolumes and their intersection in R. This study was funded through Jena Experiment SP 7 (Swiss National Science Foundation 310030E-166017/1). Further support came from the German Centre for Integrative Biodiversity Research (iDiv) Halle-Jena-Leipzig, funded by the German Research Foundation (FZT 118). The Jena Experiment was funded by the Deutsche Forschungsgemeinschaft (FOR 456 and FOR 1451) with additional support from the Friedrich Schiller University Jena, the Max Planck Institute for Biogeochemistry in Jena, and the Swiss National Science Foundation. All of the studies located at Cedar Creek included herein are funded by the US National Science Foundation’s Long-Term Ecological Research (LTER) program (DEB-1234162) and FI acknowledges funding from the LTER Network Communications Office (DEB-1545288).

## Data accessibility

We provide aggregated datasets with plant-community properties and ecosystem function data at first submission to enable editors and referees to run our main analyses. Currently, these datasets partly underlie project-specific embargo periods and need to be treated confidentially. All data will be a) uploaded to an online repository, b) submitted as supplemental files upon acceptance of the article or c) be made available within project databases after the respective project-defined embargo periods. Upon request by editors or referees, we are happy to provide all data at an earlier stage.

## Code availability

We provide R-code for running the main analyses and creating Fig. 1 and Fig. 2 based on aggregated datasets at first submission. All R-code for data crunching and analyses will be a) uploaded to an online repository, b) submitted as supplemental files upon acceptance of the article or c) be made available within project databases after the respective project-defined embargo periods. Upon request by editors or referees, we are happy to provide all R-code at an earlier stage.

## Author contributions

MJ, PM, MF and FvdP conceived the idea and designed the study; all authors except for MJ, FvdP, RM-V, CP, AR and PM contributed data; MJ developed the analytical framework and analyzed the data; RM-V constructed the phylogenetic hypothesis trees; MJ and PM wrote the manuscript; all authors contributed to the discussion of results and to the writing of the manuscript.

## Supporting Information

**Supporting Information** for Jochum et al. submission entitled

“The results of biodiversity-ecosystem functioning experiments are realistic”

The following Supporting Information is available for this article online

## Supporting Methods

### 1. Details of ecosystem function measurement in the Jena Experiment and BioDIV

#### Jena and BioDIV plant aboveground biomass

In Jena, aboveground plant biomass was harvested bi-annually (late May and late August), just prior to mowing. Here, we used only the first harvest, which represents peak standing biomass in most years, from years 2006–2015. All vegetation was clipped at 3 cm above ground in up to four rectangles of 0.2 m × 0.5 m per plot with the location of these rectangles being randomly assigned each year. For BioDIV, aboveground peak plant biomass was harvested annually in August by clipping 0.1 m × 6 m strips (see above) each year from 2006–2015. For both studies, harvested target-species biomass was sorted into individual species, dried to constant weight at 70 °C for at least 48 h and weighed. Target plant community biomass was then calculated as the sum of the biomass of the individual sown species (g m^-2^).

#### Jena and BioDIV aboveground plant biomass C:N ratio

In Jena, the combined target species material from the spring biomass harvest (May) was shredded (Analysenmühle, Kinematica, Littau, Switzerland). A subsample of the shredded material was milled to fine powder in a ball-mill (mixer mill MM2000 Retsch, Haan, Germany) and 5–10 mg was used for CN analysis with an elemental analyzer. C and N content were calculated as percentage elemental concentration of dry material and C:N ratios as the ratio between those percentages for years 2007-2012.

In BioDIV, two strips of 0.1 m × 6 m were clipped, typically in late July or early August with clip strip locations rotated each year. Unsorted biomass was air-dried at 40 °C. Dried biomass samples were ground (standard Thomas Wiley mill) and the resulting sample homogenized. A sub-sample was re-ground in a Wiley Mini-Mill, stored in glass scintillation vials and re-dried prior to lab analysis. Percent C and N content in dry matter were determined using an elemental analyzer (NA1500, Carlo-Erba Instruments or ECS 4010, COSTECH Analytical Technologies Inc., Valencia, CA, USA) at University of Minnesota or at the Ecosystems Analysis Lab, University of Nebraska, Lincoln. Ratios of dry mass elemental content were then calculated from these results for year 2006.

#### BioDIV total soil C

Total soil C samples were taken at all BioDIV plots during summer 2006 at 0–20 cm depth on nine sites per plot^1^. Samples were sieved to remove roots, combined for each plot, mixed and ground. Subsequently, soil samples were dried at 40 °C for 5 days. For each plot, two soil samples were analysed for total C by combustion and gas chromatography (Costech Analytical ECS 4010 instrument, Costech Analytical Technologies Inc., Valencia, CA). We used the average of the two measurements of C in % total carbon of dry weight.

#### Jena soil organic C

Soil organic C in the Jena “main” experiment was determined in 2008, 2011 and 2014. Using a split-tube sampler (4.8 cm diameter), three soil cores per plot were taken to a depth of 30 cm ^2^. Soil cores were segmented into 5 cm depth sections and pooled per depth sections and plot. Soil was then dried, sieved and milled. Subsequently, total C was determined by combustion with an elemental analyzer at 1,150 °C (Elementaranalysator vario Max CN, Elementar Analysensysteme GmbH, Hanau, Germany). Inorganic C concentration was measured after oxidative removal of organic C for 16 h at 450 °C in a muffle furnace. Finally, organic C concentration was calculated as the difference between total and inorganic C for each 5-cm-layer^2^ and we averaged over the two uppermost layers to get organic C content for 0–10 cm depth. Subsequently, we averaged over the three samples to get soil organic C content per year and plot in g kg^-1^ soil.

#### Jena and BioDIV root biomass

In Jena, standing root biomass was sampled down to 40 cm depth in all plots in June 2011 and 2014. On each plot, three cores of 3.5 cm diameter were taken and immediately stored at 4 °C until further handling. The total sample was washed to determine root biomass. Bulk samples were carefully washed by hand over a sieve of 0.5 mm mesh size. Remaining soil particles and stones were removed with tweezers. Roots were dried at 60–70 °C and weighed subsequently^3^. Unit: g m^-2^ In BioDIV, root biomass was sampled in 2010 after aboveground biomass clipping by collecting three 5 cm diameter × 30 cm depth cores per clipped strip^1^. Roots were washed free of soil, sorted from other organic material, dried and weighed. Unit: g m^-2^

#### Jena herbivory rate

In Jena, invertebrate herbivory rates were assessed as proportional damage for every plant species × plot-combination. Herbivory rates of individual plant species were used to calculate community herbivory rates based on four different types of invertebrate herbivory: chewing, rasping, sap sucking and leaf mining. Samples of the Jena biomass harvest were used after sorting to species. For a maximum of 20 randomly chosen leaves per plant species, damage area was estimated in mm^2^ as total value of the four damage types and total leaf area of every leaf was measured with an area meter (LI-3000C Area Meter equipped with a LI3050C transparent belt conveyor accessory, LI-COR Biosciences, Lincoln, USA). For details on the methods used see^4^. Here, we used percentage herbivory of the target species community from the late harvest, as this was available for three years from 2010– 2012. Unit: % damage

#### Jena soil microbial biomass C

Soil sampling and measurement of basal and substrate-induced microbial respiration with an oxygen-consumption apparatus was done on each plot in September 2010^5^. Oxygen consumption of soil microorganisms in a fresh-soil equivalent to 3.5 g dry weight was measured at 22 °C. Substrate-induced respiration was determined by adding D-glucose to saturate catabolic enzymes of microorganisms according to preliminary studies (4 mg g^-1^ dry soil solved in 400 µl deionized water; ^6, 7^). Maximum initial respiratory response (µl O_2_ g^-1^ dry soil h^-1^) was calculated as mean of the lowest three oxygen consumption values within the first 10 h after glucose addition. Microbial biomass C (µg C g^-1^ dry soil) was calculated as 38 × maximum initial respiratory response as suggested by preliminary studies^8^. Previous work has shown that the 2010 microbial biomass data are representative for long-term plant diversity effects^7^.

#### Jena phosphatase activity

Nine soil cores (diam. 2 cm, 0–5 cm depth) were combined to one composite sample per plot to assess phosphatase activity in 2013^9^. Because of the alkaline pH of the soil, we measured alkaline phosphomonoesterase activity (phosphatase activity) according to the assay by ^10^. For each soil sample, one replicate and one blank value were included. One gram of field moist soil was mixed with toluene, modified universal buffer (MUB) and p-nitrophenylphosphate (pNP), and incubated at 37 °C for 1 hour. Subsequently, we added CaCl_2_ and NaOH. To blanks pNP was added after incubation. The solution was filtered through P-free filters (MN 619 G ¼, Macherey-Nagel GmbH & Co. KG, Düren, Germany). Directly after filtration, pNP concentrations [µg ml^-1^] were measured at 400 nm with a spectrophotometer (PU 8675 VIS spectrophotometer, Philips GmbH, Hamburg, Germany). The soil moisture was determined gravimetrically, i.e. by weighing before and after drying at 105 °C to convert phosphatase activities to dry matter (µg pNP g^-1^ DM h^-1^).

#### Jena pollinator abundance

In 2010 and 2012, hymenopterans were sampled by suction sampling using a modified commercial vacuum cleaner (Kärcher A2500, Kärcher GmbH, Winnenden, Germany). In each year, within each plot, two random subplots of 0.75 m x 0.75 m were chosen, covered with a gauze-coated cage of the same size, and arthropods within cages were sampled. The sampling was carried out between 9 a.m. and 4 p.m. within two 4-day sampling periods. The overall abundance of hymenopterans across the two samples per plot was used as a proxy of pollinator abundance and thus potential for pollination on each plot in the respective year. Unit: number of individuals

### 2. Processing TRY and other plant-trait data to generate species-level values

For each of the geographical species subsets, TRY trait data were processed separately following a standardized protocol: i) Removal of duplicate observations (e.g. duplicate entries of leaf mass from the same individual). ii) Removal of non-open data and removal of data obtained from outside the respective target continents. iii) Calculation of outliers for each trait-species combination (trait mean +/-1.96 SD as outlier definition). iv) Removal of observations with TRY ErrorRisk > 4. v) Averaging over trait-species values per TRY dataset. vi) Removal of TRY datasets with more than 5% of values identified as outliers. vii) Averaging over trait-species mean values of the remaining datasets. For the US species, TRY data was combined with additional trait data collected in naturally occurring polycultures at Cedar Creek (personal communication with J.A. Catford^11^, P.B. Reich, J. Cavender-Bares). Such Cedar Creek trait averages per dataset were included into the averaging process at step v). Finally, trait values of synonyms and accepted species names were averaged and assigned to the accepted plant-species names where necessary.

---

**Figure S1.**
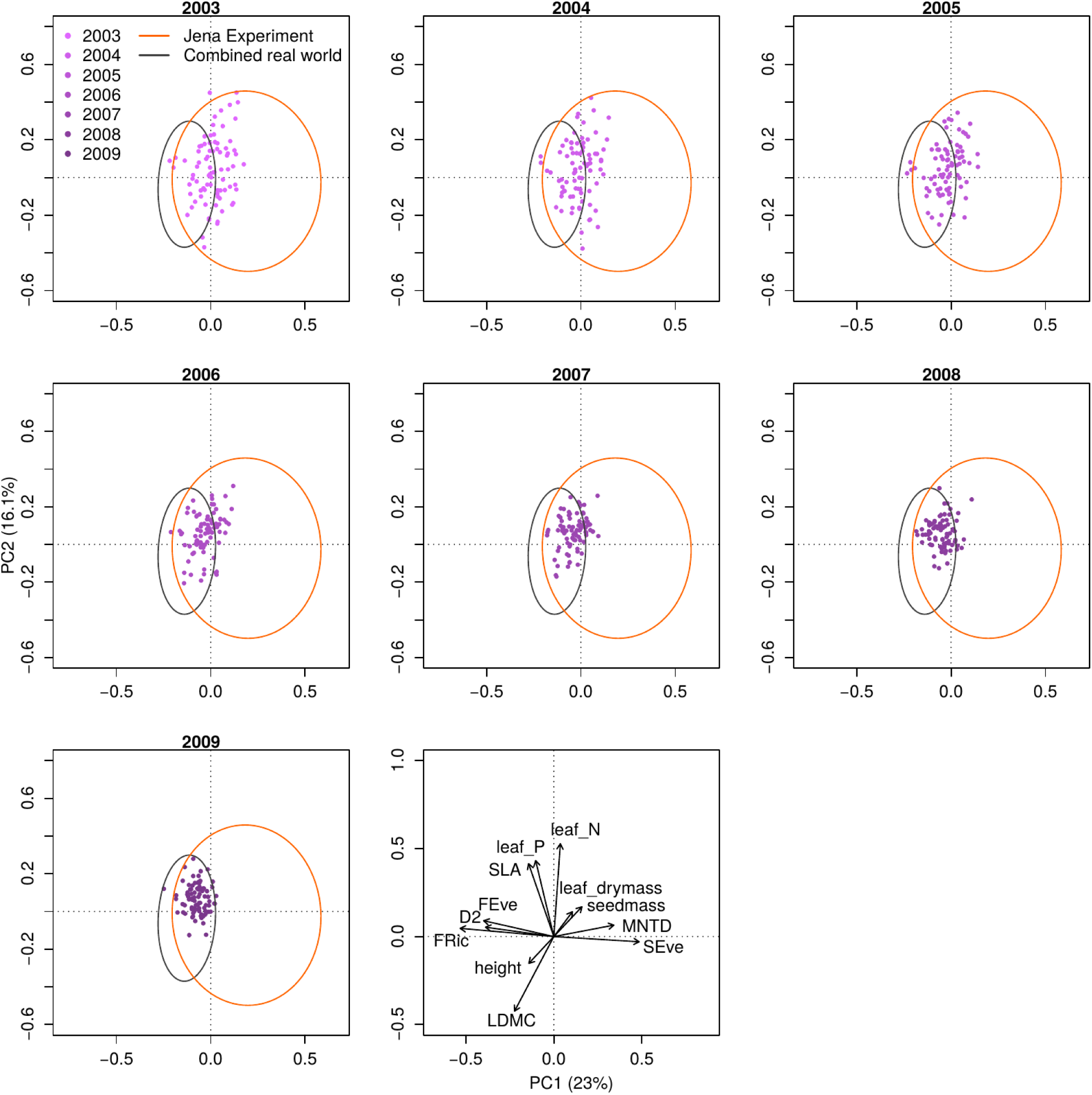
Temporal movement of Jena invasion plots into the real-world realm. Based on the PCA in Fig. 1a. Different shades of purple show Jena invasion plots across the years from 2003-2009. Orange and gray ellipses show 95% confidence intervals for Jena Experiment and combined real-world plots, respectively. Note that while the points in different panels are from single years, the ellipses are fixed to the across-year comparison in Fig. 1a. The last panel shows the PCA factor loadings for the full 12 community properties (arrows scaled to improve visibility). Within six years of succession, the plant communities of Jena invasion plots fully “moved” into the core of the community property space defined by the combined real-world plots (German real world and Jena real world, respectively).

**Figure S2.**
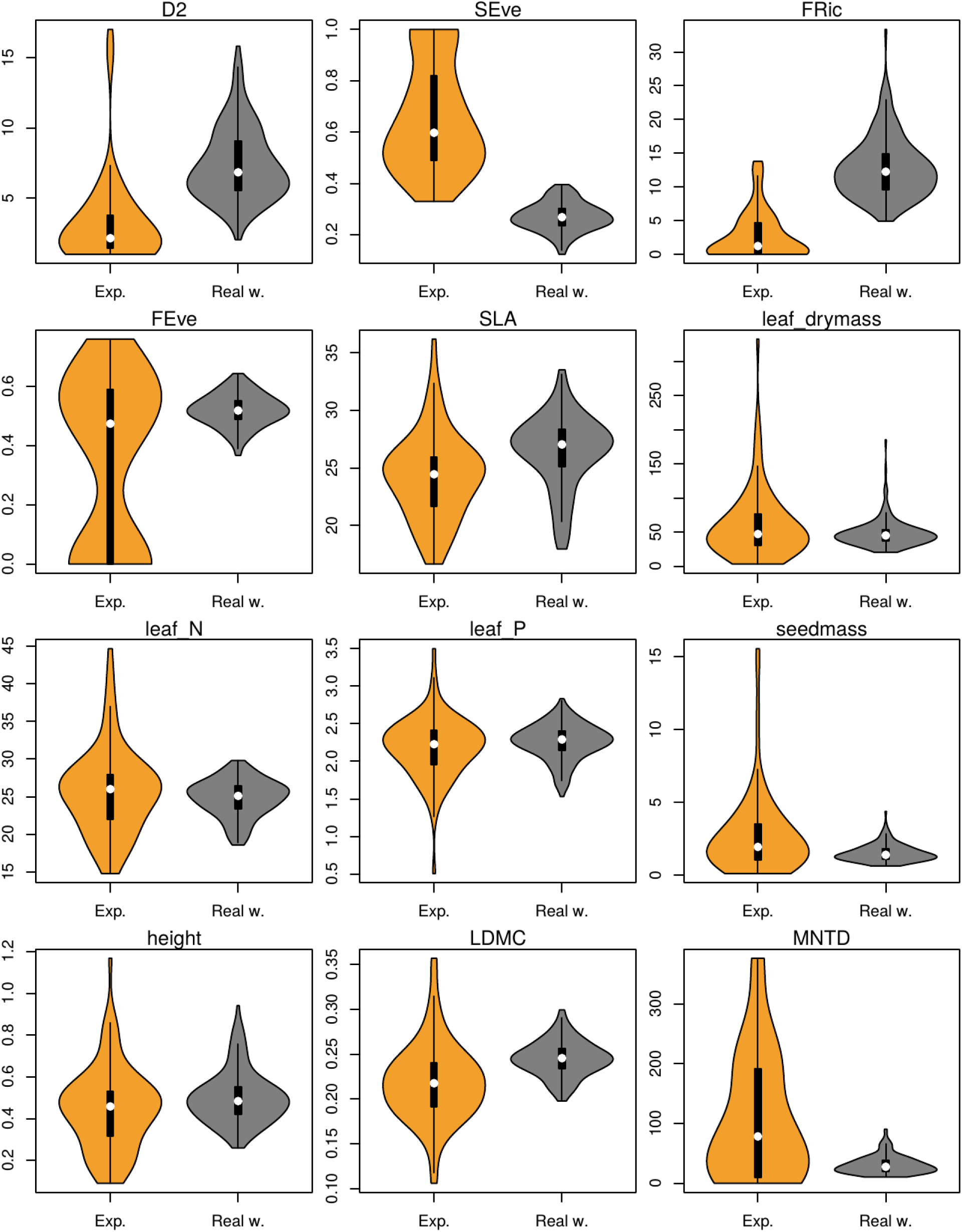
Violin plots of community properties of German experimental and real-world plots. Combination of boxplot and rotated kernel density plot (R package “vioplot”^20^). Jena Experiment (orange) and combined real-world properties (German real world, Jena real world, gray) averaged across all years per plot.

**Figure S3.**
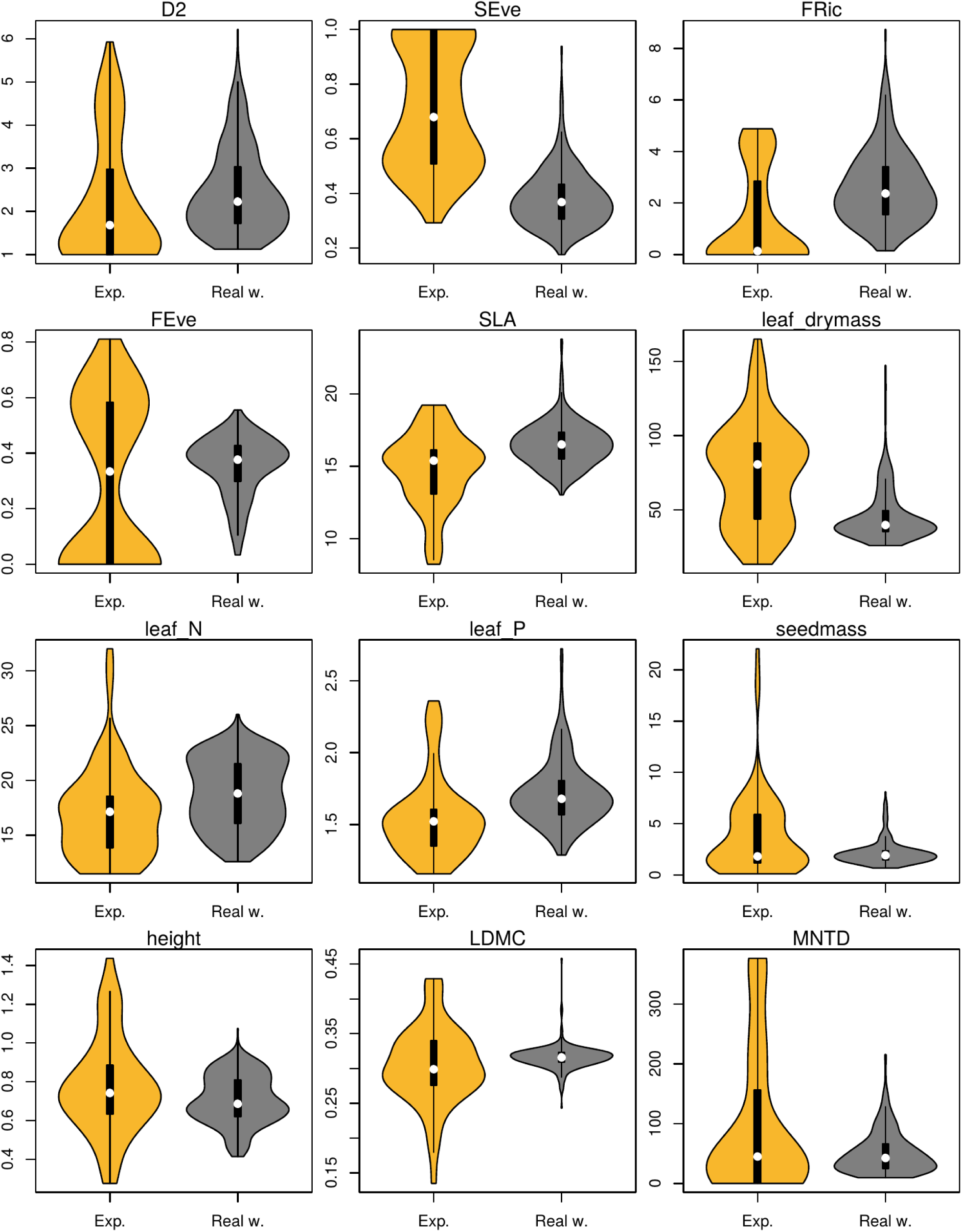
Violin plots of community properties of US experimental and real-world plots. Combination of boxplot and rotated kernel density plot (R package “vioplot”^20^). BioDIV (orange) and combined real-world data (Fertilization 1 & 2, gray) averaged across all years per plot.

**Figure S4.**
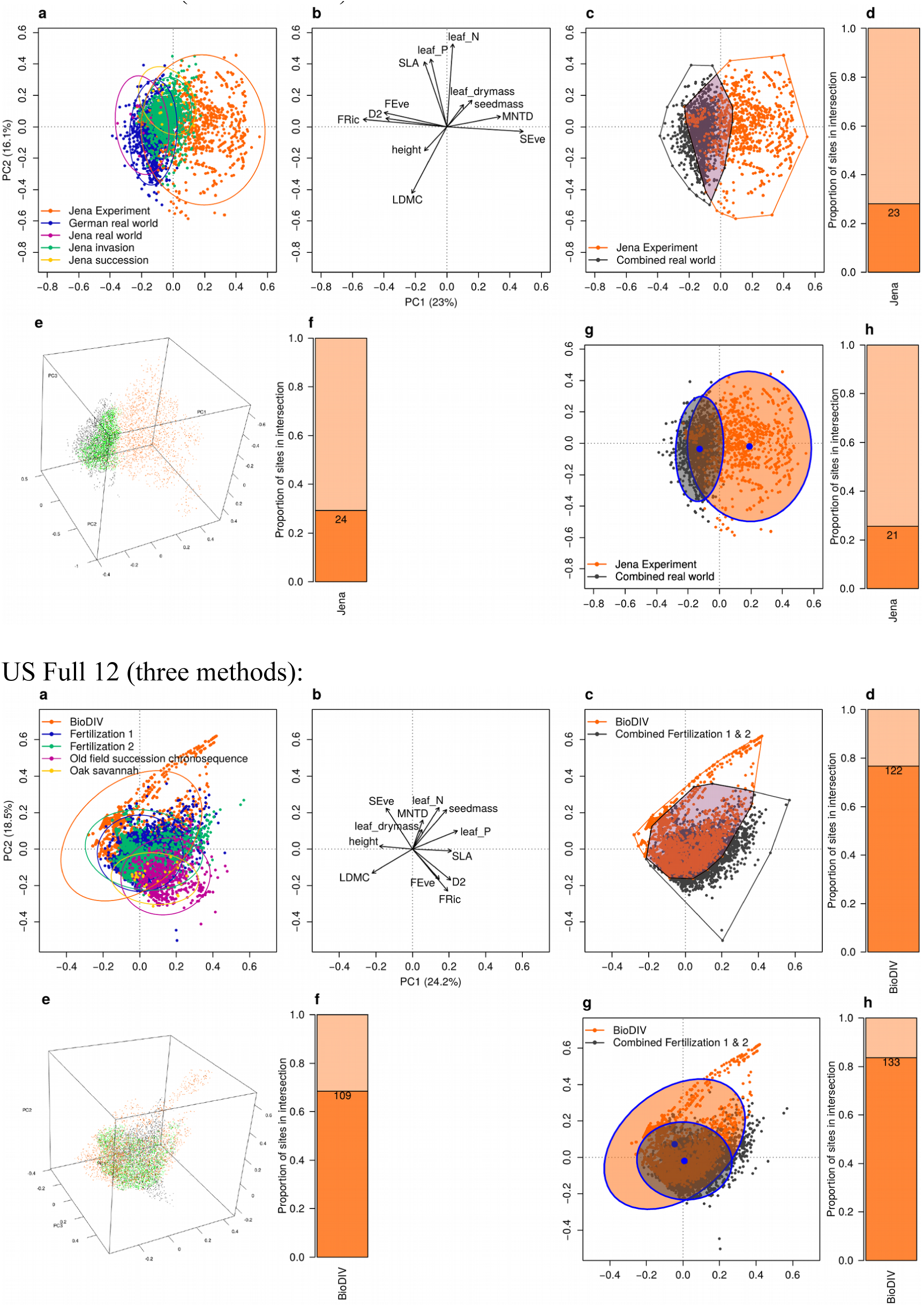
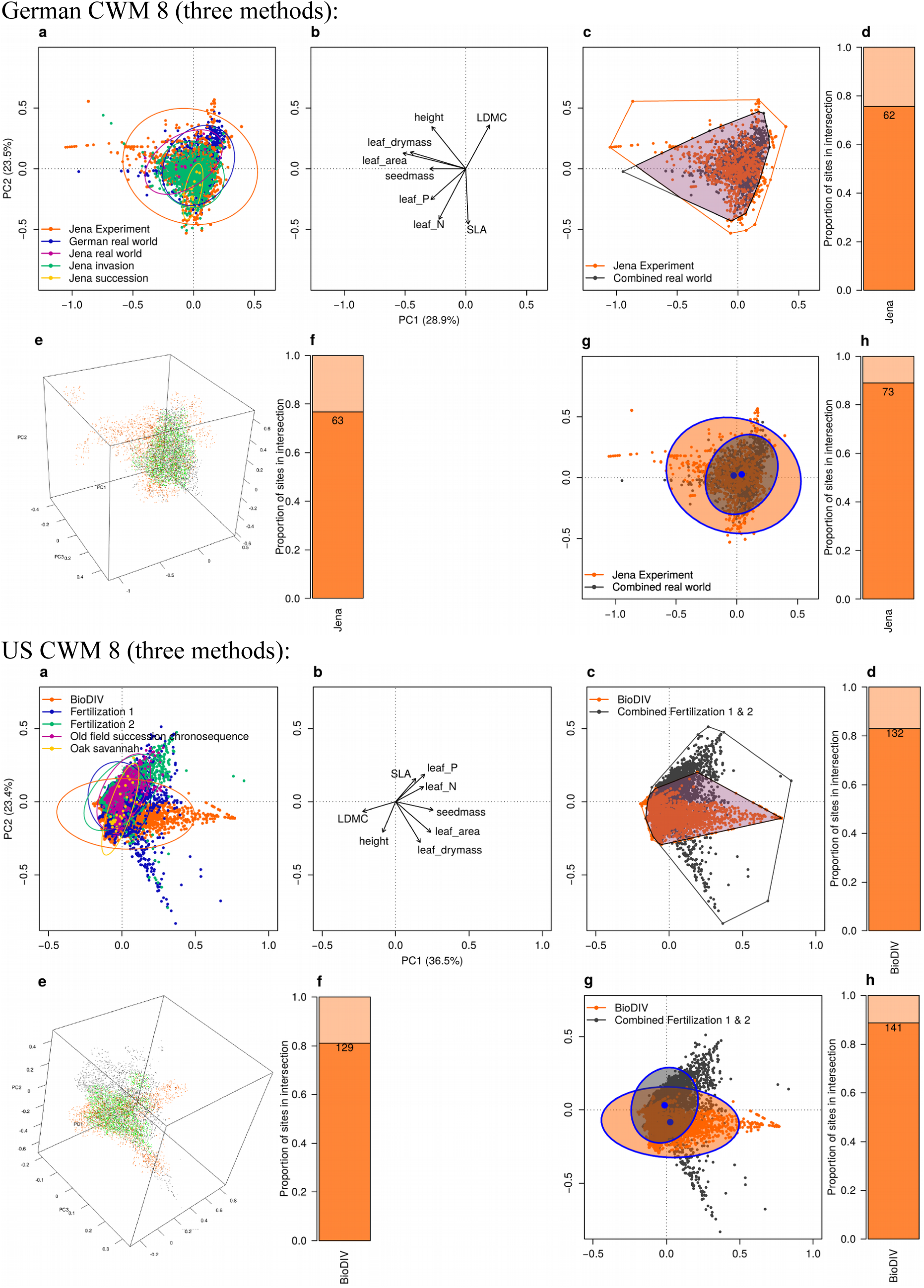
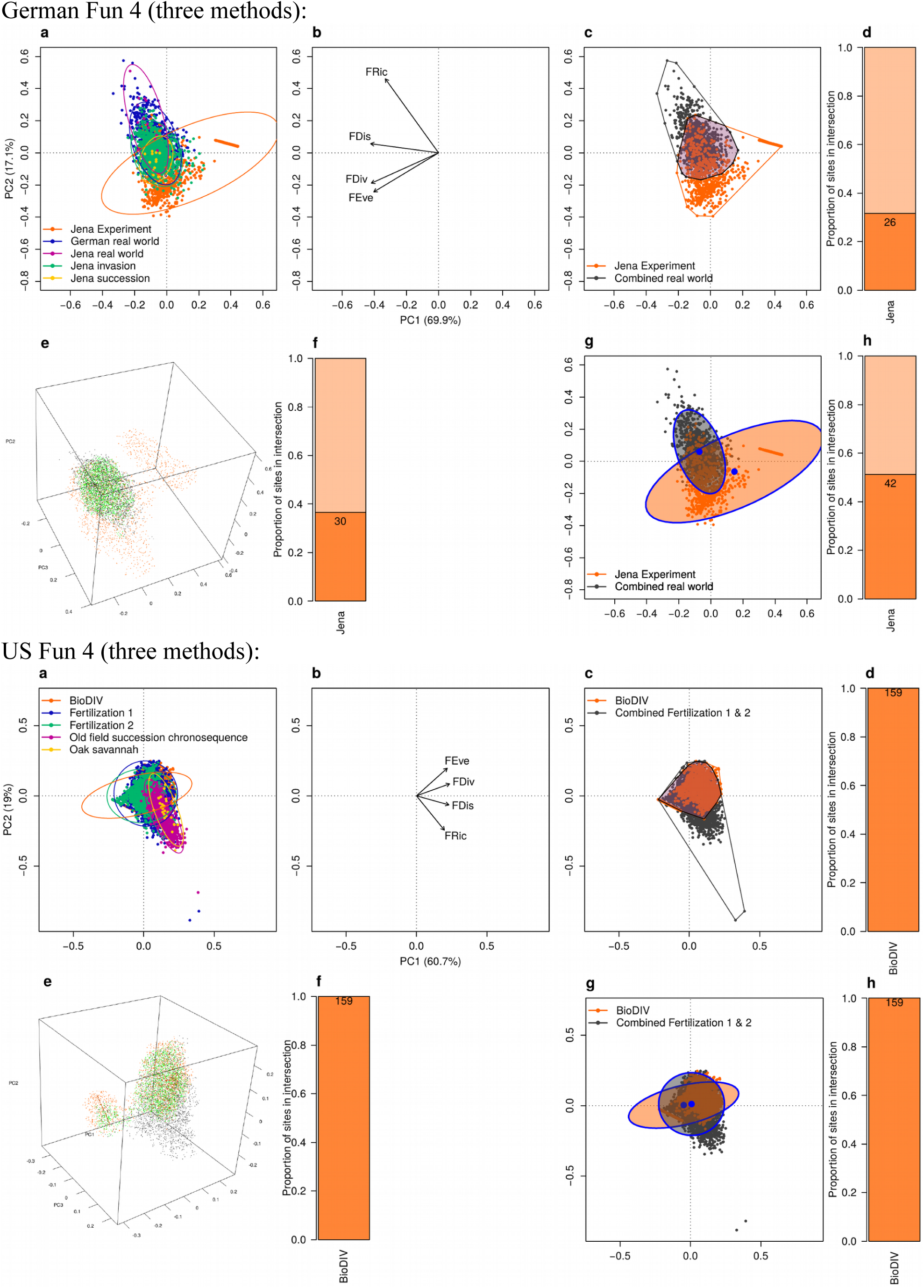
Alternative versions of Fig. 1 based on the alternative intersection scenarios. 6 Versions: One for each geographical dataset per community property subset, combining all three methods. Panels a & b: PCA and factor loadings; c & d: 3D convex hull volume, e & f: 3D hypervolume, g & h: 2D ellipse German Full 12 (three methods):

**Figure S5.**
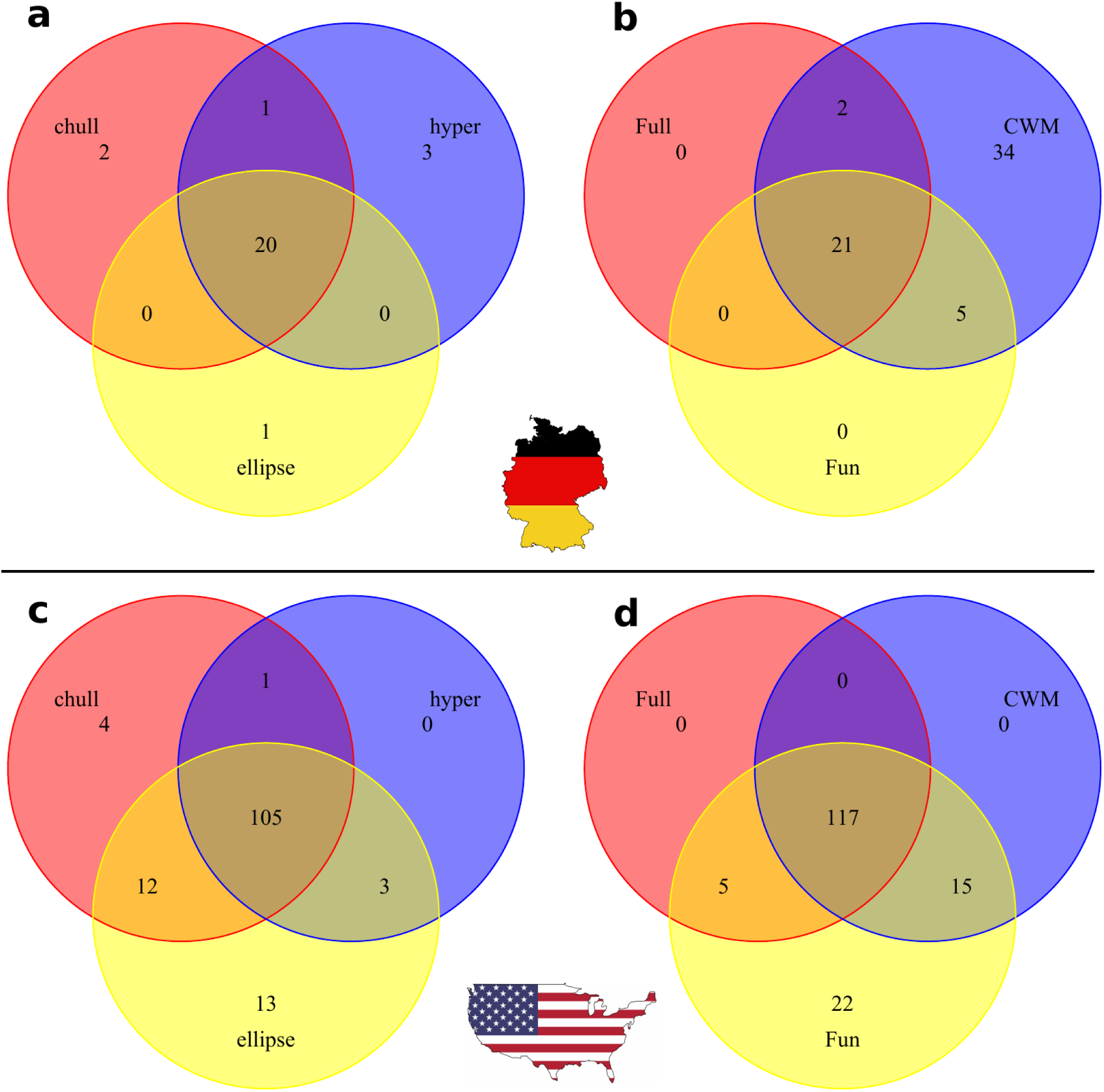
Venn diagrams illustrating overlap between plots defined as realistic for the Jena Experiment (upper row) and BioDIV (lower row) based on three different methods of calculating intersections (a and c) and three different subsets of community properties entering the PCA’s (b and d). a and c show three different methods for the PCA, all based on the full set of 12 properties. b and d show three different subsets based on just the convex hull method. Abbreviations: chull=convex hull volume approach, hyper= hypervolume approach, ellipse=confidence ellipse approach, Full=all 12 community properties, CWM=just the eight community weighted mean traits, Fun=just the four functional diversity properties. Diagrams were created with R package “VennDiagram”^21^.

**Figure S6.**
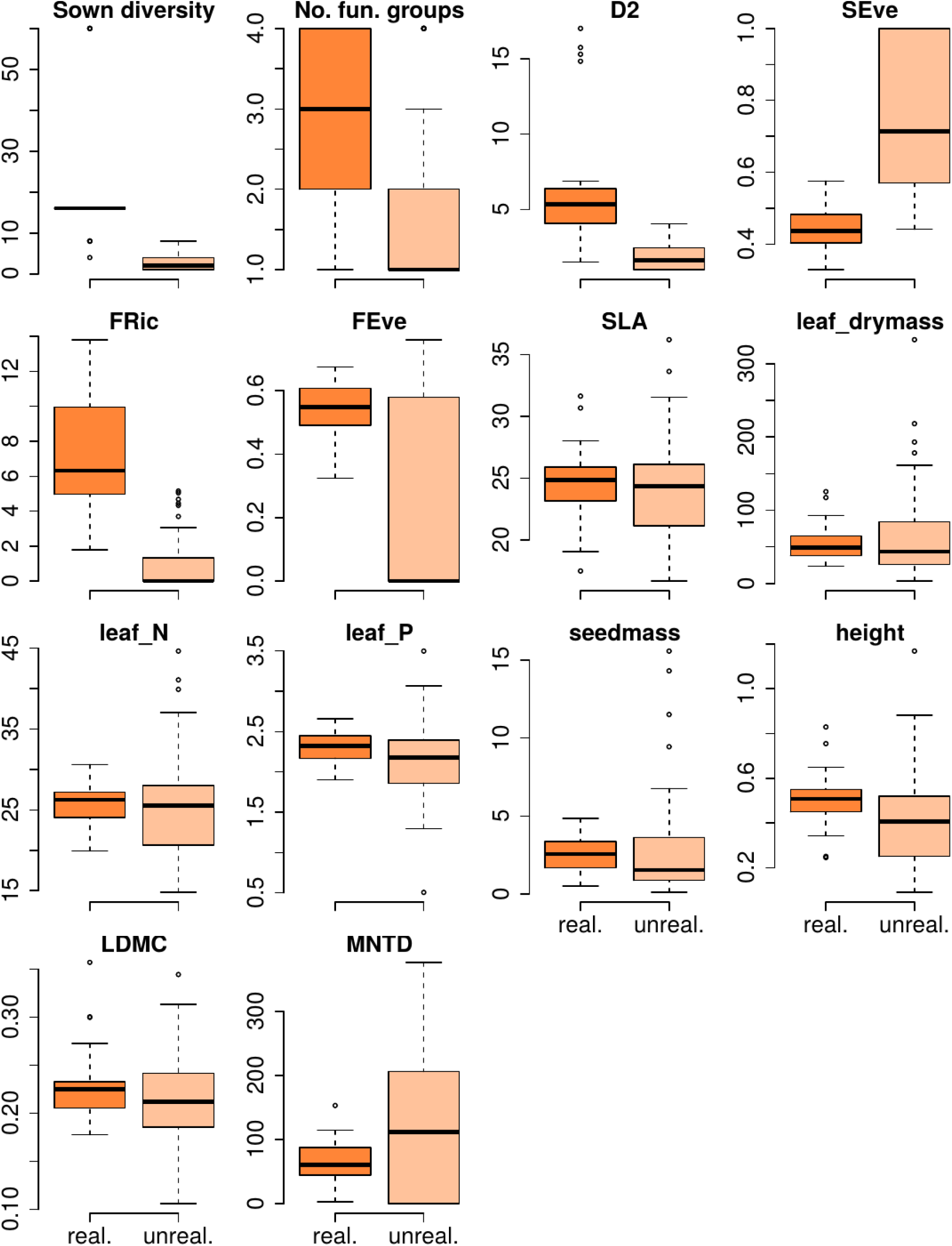
Boxplots of community properties of realistic (strong color) and unrealistic (weak color) plots for the Jena Experiment. Realistic plots were calculated based on the full set of community properties and the convex hull volume method. All properties were averaged across all available years per plot (23 realistic and 59 unrealistic plots).

**Figure S7.**
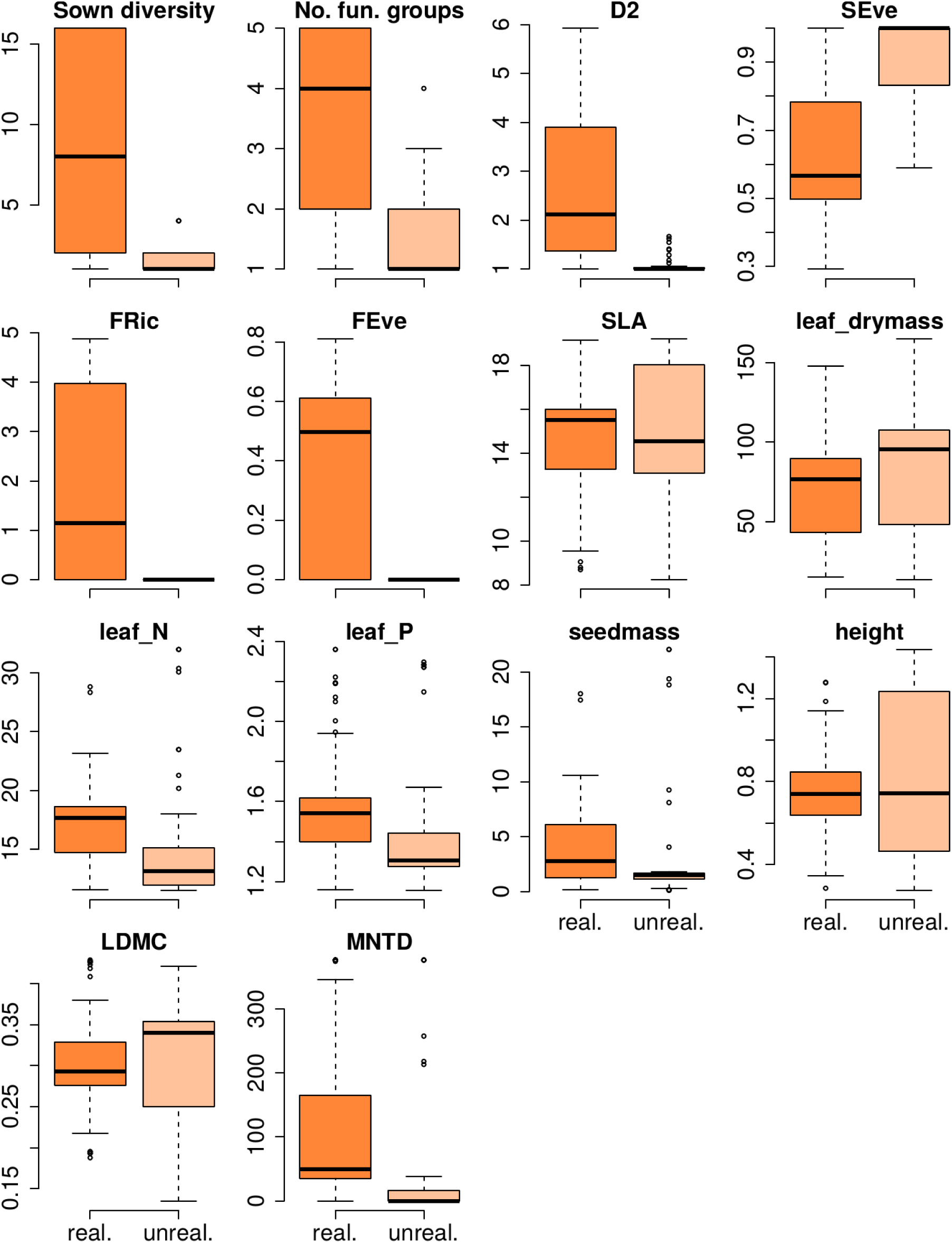
Boxplots of community properties of realistic (strong color) and unrealistic (weak color) plots for BioDIV. Realistic plots were calculated based on the full set of community properties and the convex hull volume method. All properties were averaged across all available years per plot (122 realistic and 37 unrealistic plots).

**Figure S8.**
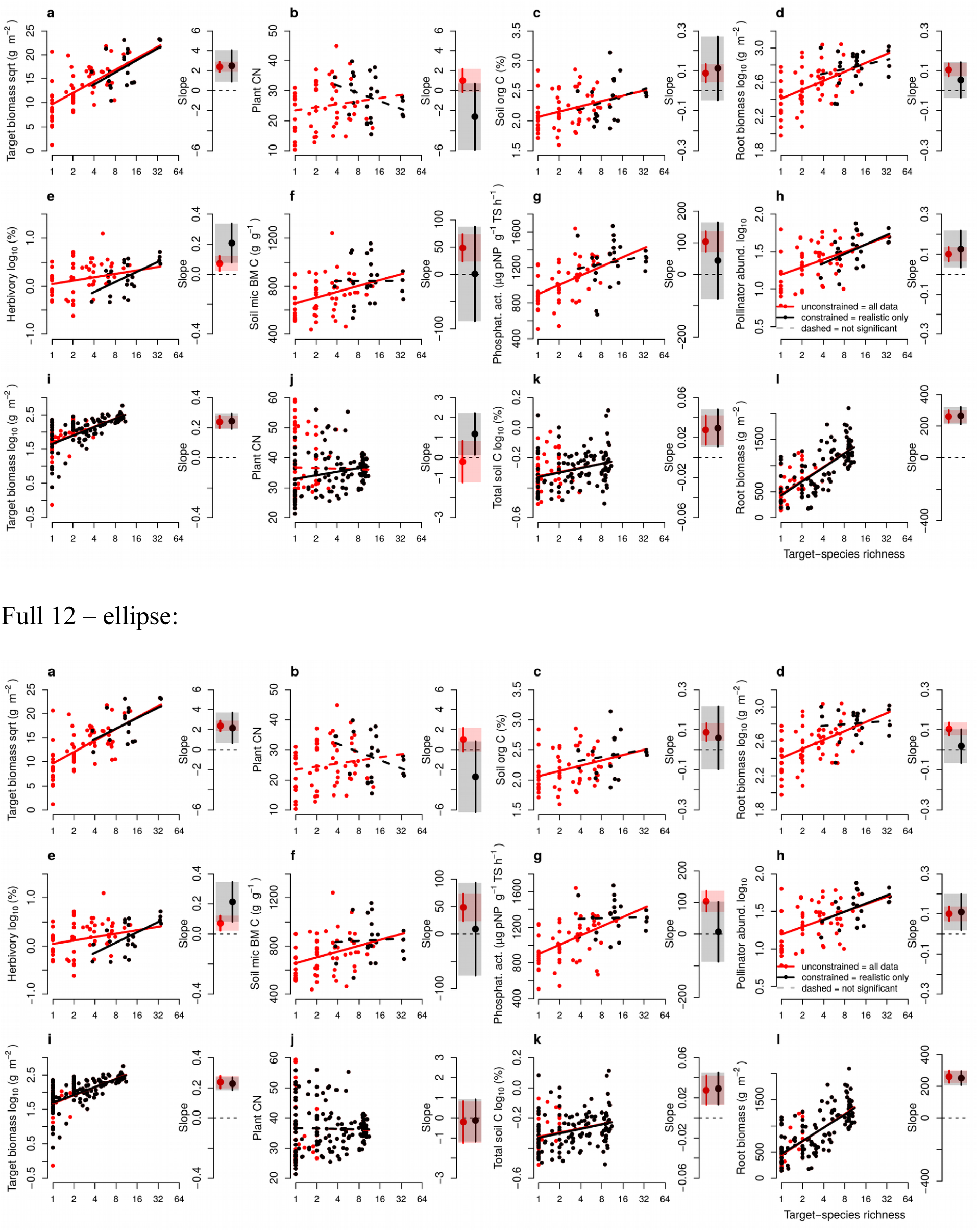
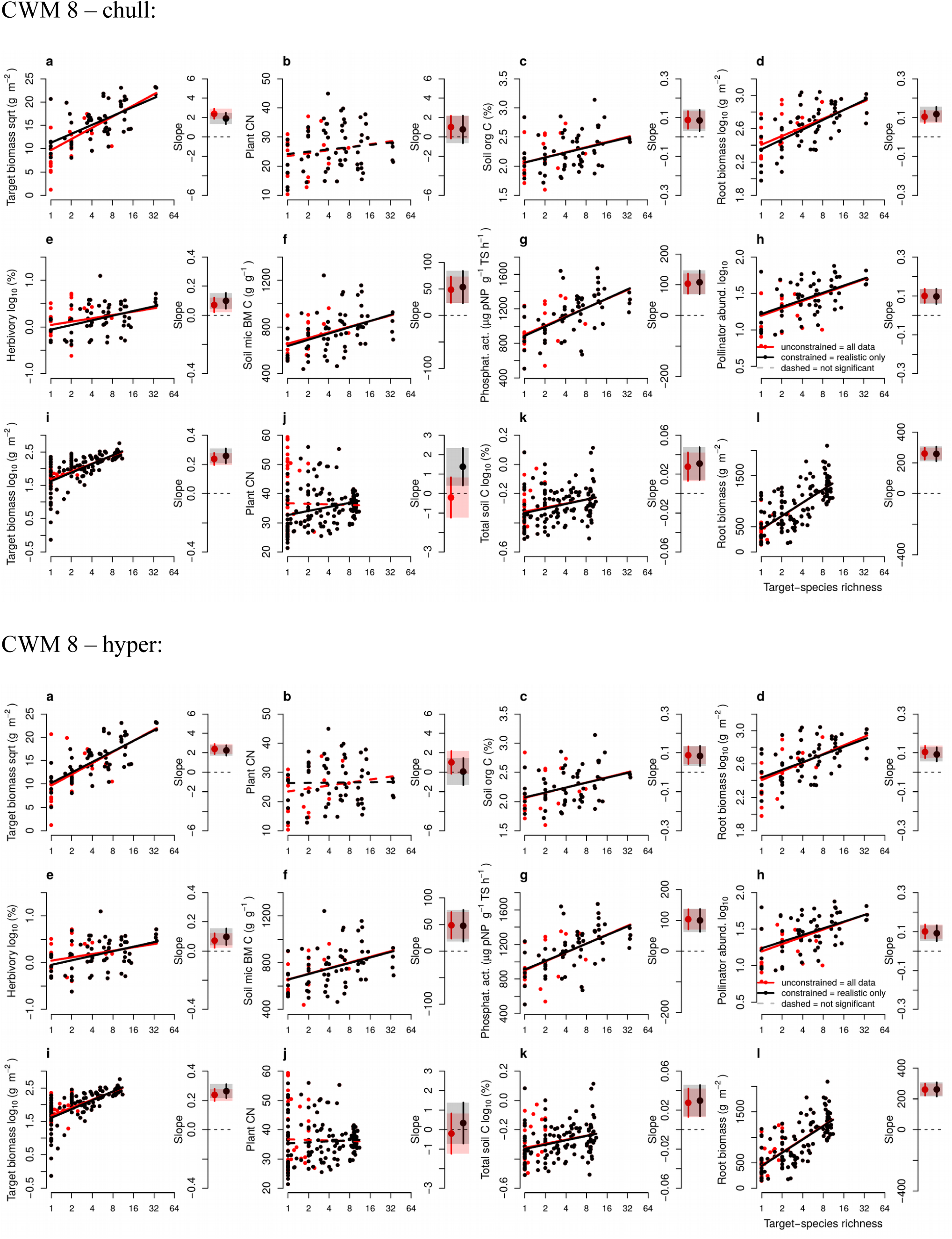
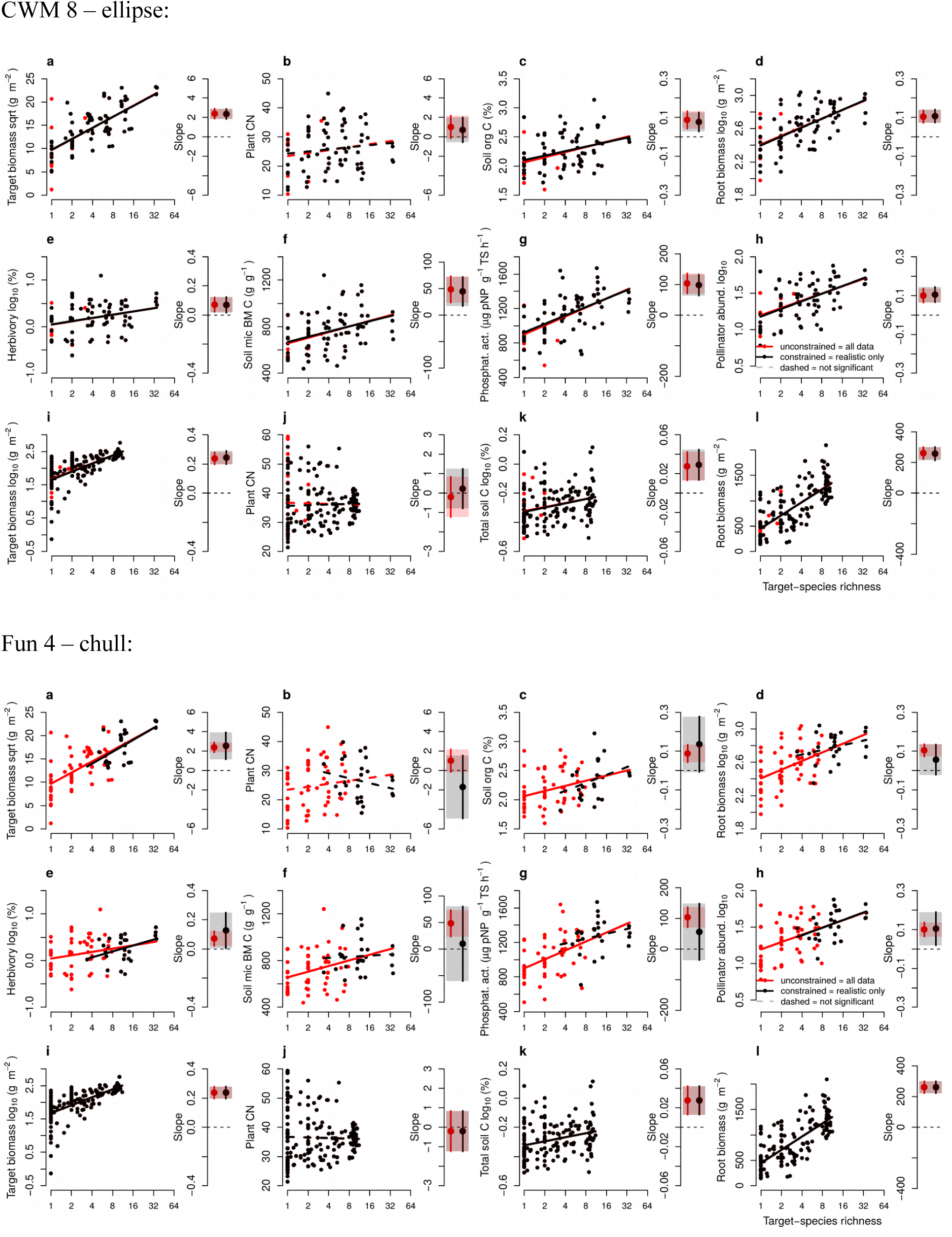
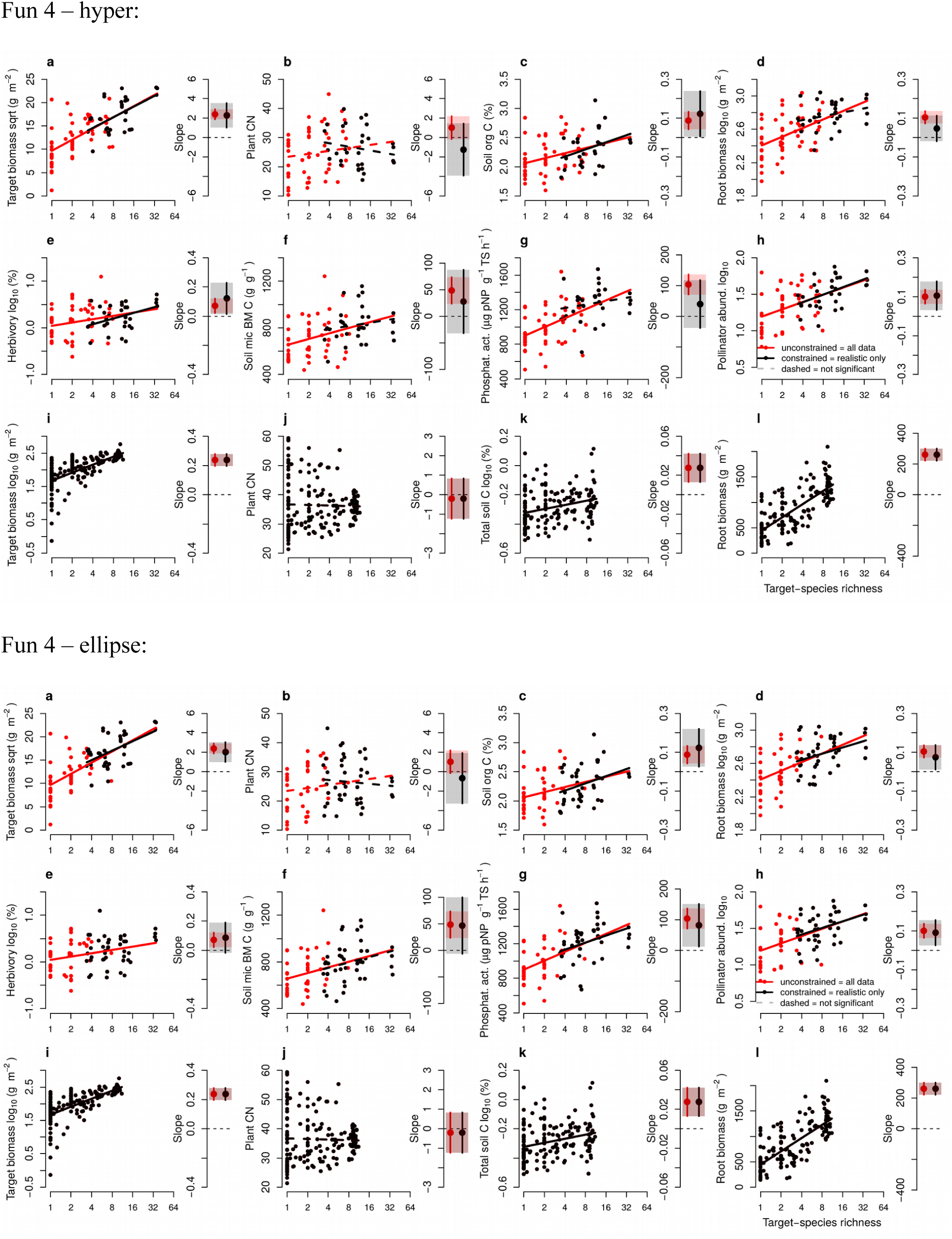
Alternative versions of Fig. 2 based on the alternative intersection scenarios. Panels a-h Jena, panels i-l BioDIV (see main text Fig. 2). 8 different versions: 3 methods and 3 community property subsets (but convex hull method with full 16 properties shown in main text already).

**Figure S9:**
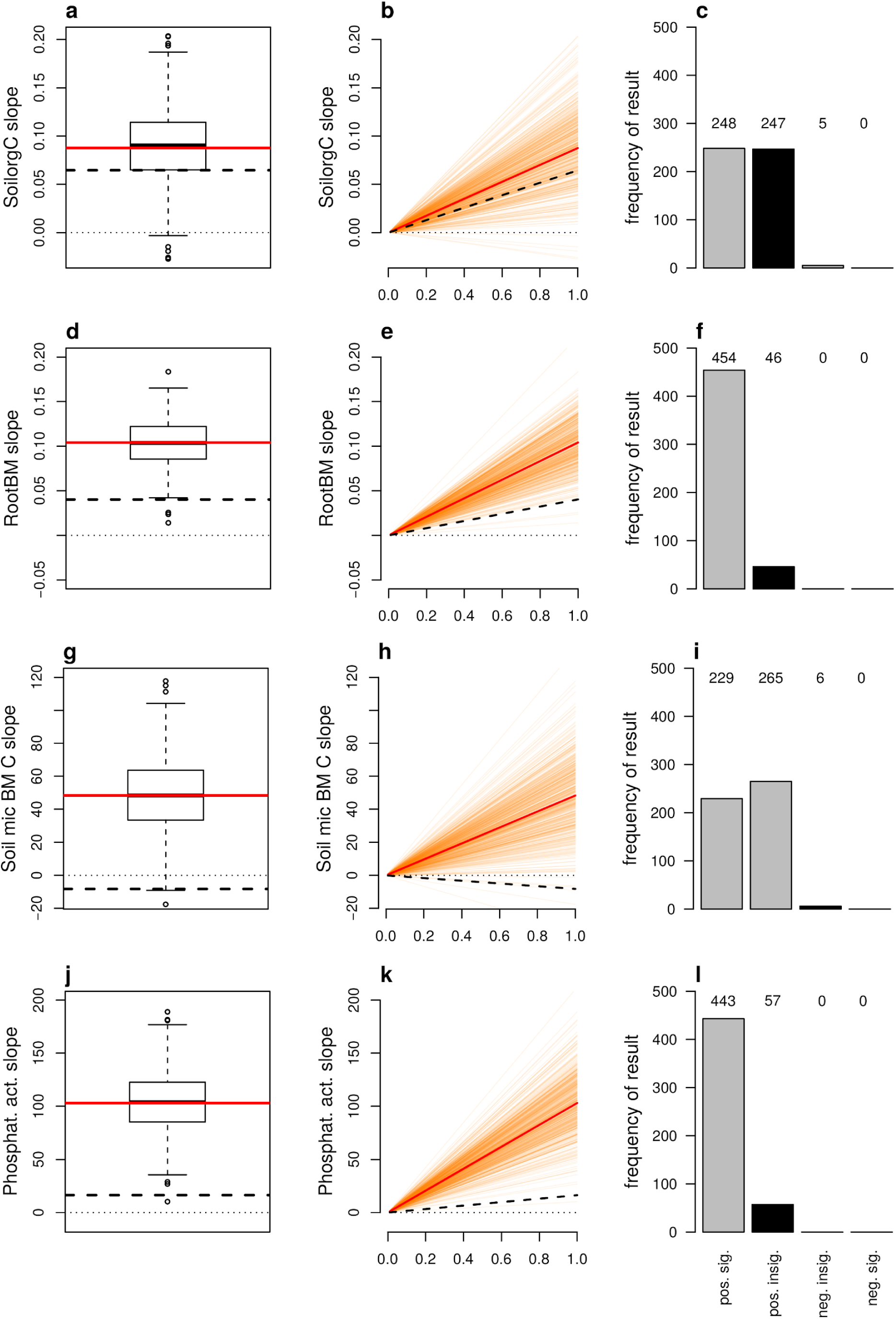
Random selection sensitivity analysis for Fig. 2 relationships turning insignificant. We performed a sensitivity analysis testing if changes in BEF relationships from being significant (all communities) to non-significant (realistic sites only) were likely caused by the related reduction in sample size or if a randomly-selected reduced number of plots was still likely to result in a significant relationship. Therefore, for each of the four BEF relationships found to switch significance (Jena soil organic C (a-c), root biomass (d-f), soil microbial biomass C (g-i) and phosphatase activity (j-l)), we repeatedly (500 times), randomly selected 23 Jena plots and re-ran the model testing for the BEF relationship and saving the slope estimates and p-values. This figure shows the distribution of these 500 random-selection slopes (boxplots in first column and orange lines in middle column) in comparison to the unconstrained (all sites, red lines) and constrained (PCA-selection based realistic sites only, black dashed lines) slopes from Fig. 2. Dotted black lines indicate zero slopes. The right column shows the frequency of positive significant, positive insignificant, negative insignificant and negative significant relationships obtained by the 500 random subsets of 23 plots with the black bar highlighting the PCA-based realistic result from Fig. 2. The sensitivity analysis shows that black dashed lines and the results of the PCA-based realistic subset divert relatively strongly from the 500 random-selection results. Specifically, the PCA-based realistic subset resulted in strikingly shallower slopes than the random choices and non-significantly positive or even negative relationships while a big part of the random subsets resulted in significant positive or at least non-significantly positive relationships. As such, our PCA-based selection of realistic sites is highly non-random in comparison to the random-selection of plots, thus indicating that our methodology is successful in finding a subset of plots based on prior knowledge (realistic plots based on the multidimensional, multivariate comparison of communities) and does not simply create a random subset of plots. Furthermore, these results show that, for these four Jena soil processes, experiment-derived BEF relationships might not be as important or strong in real-world systems, at least as long as plant communities in experiments deviate from those in real-world systems. Future developments of real-world plant communities due to global change drivers and increasing anthropogenic pressure might change this conclusion by rendering less diverse communities realistic, thus aligning the species richness gradients of biodiversity experiments and related real-world systems and increasing the slope of the BEF relationships.

**Figure S10.1.**
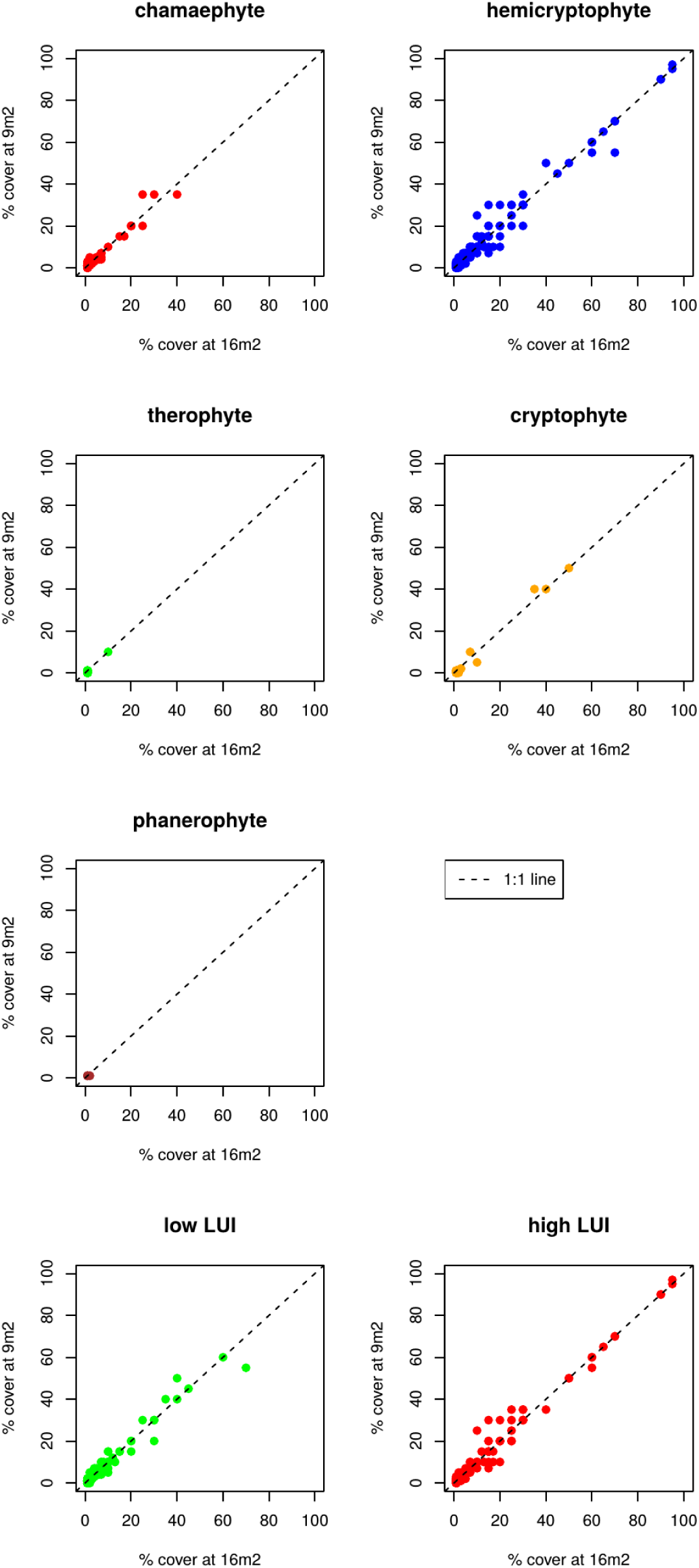
Cover versus vegetation survey size scaling sensitivity check for Biodiversity Exploratories (German real world). Here, 16 to 9 m², which is the vegetation survey area of the Jena main and Jena real world plots. For this figure, species were sorted into lifeforms using the R package “TR8”^23^ and information from The Ecological Flora Database^24^.

**Figure S10.2.**
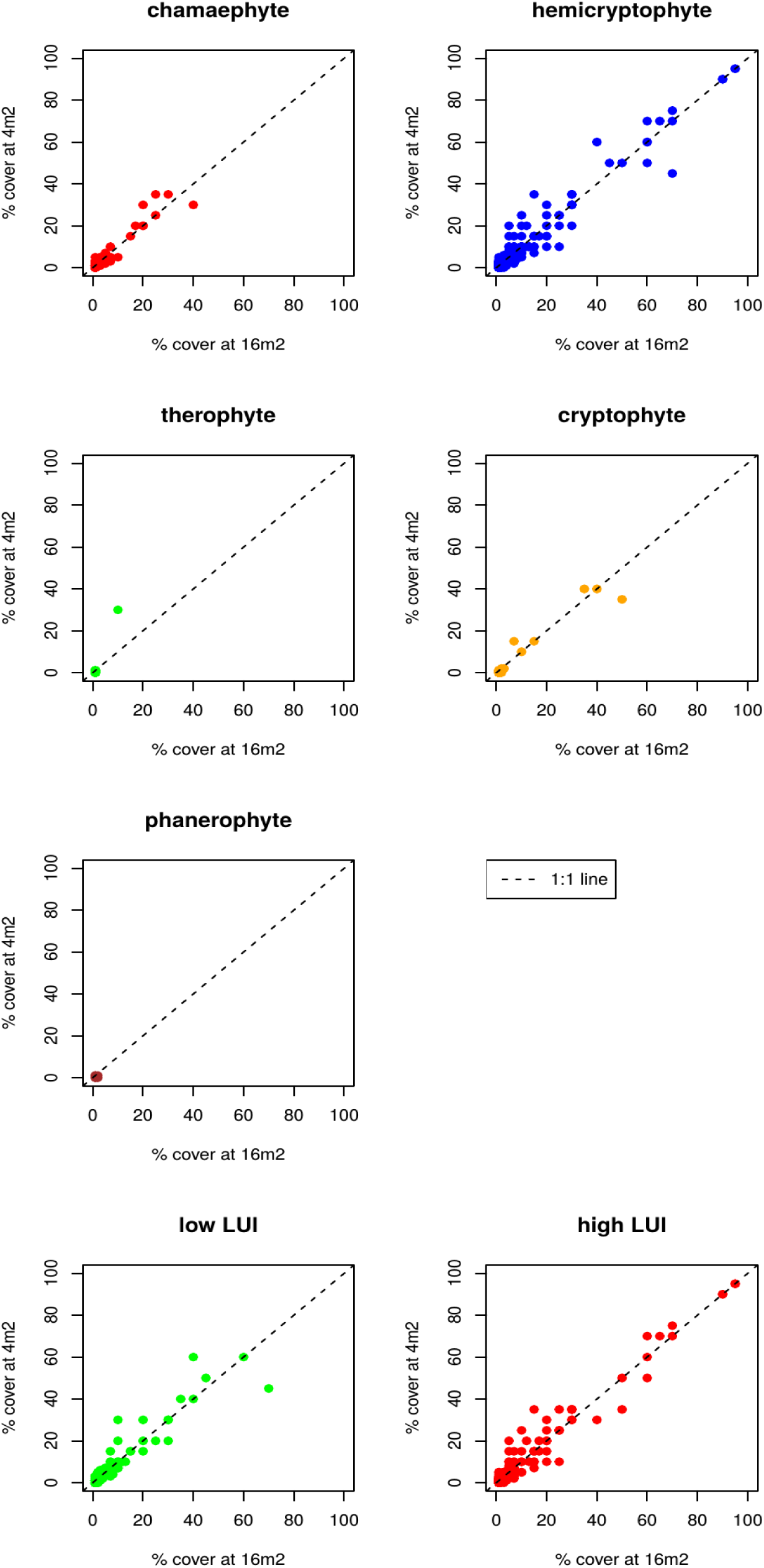
Cover versus vegetation survey size scaling sensitivity check for Biodiversity Exploratories (German real world). Here, 16 to 4 m², which resembles the vegetation survey area of the Jena invasion plots. For this figure, species were sorted into lifeforms using the R package “TR8”^23^ and information from The Ecological Flora Database^24^.

**Figure S11.**
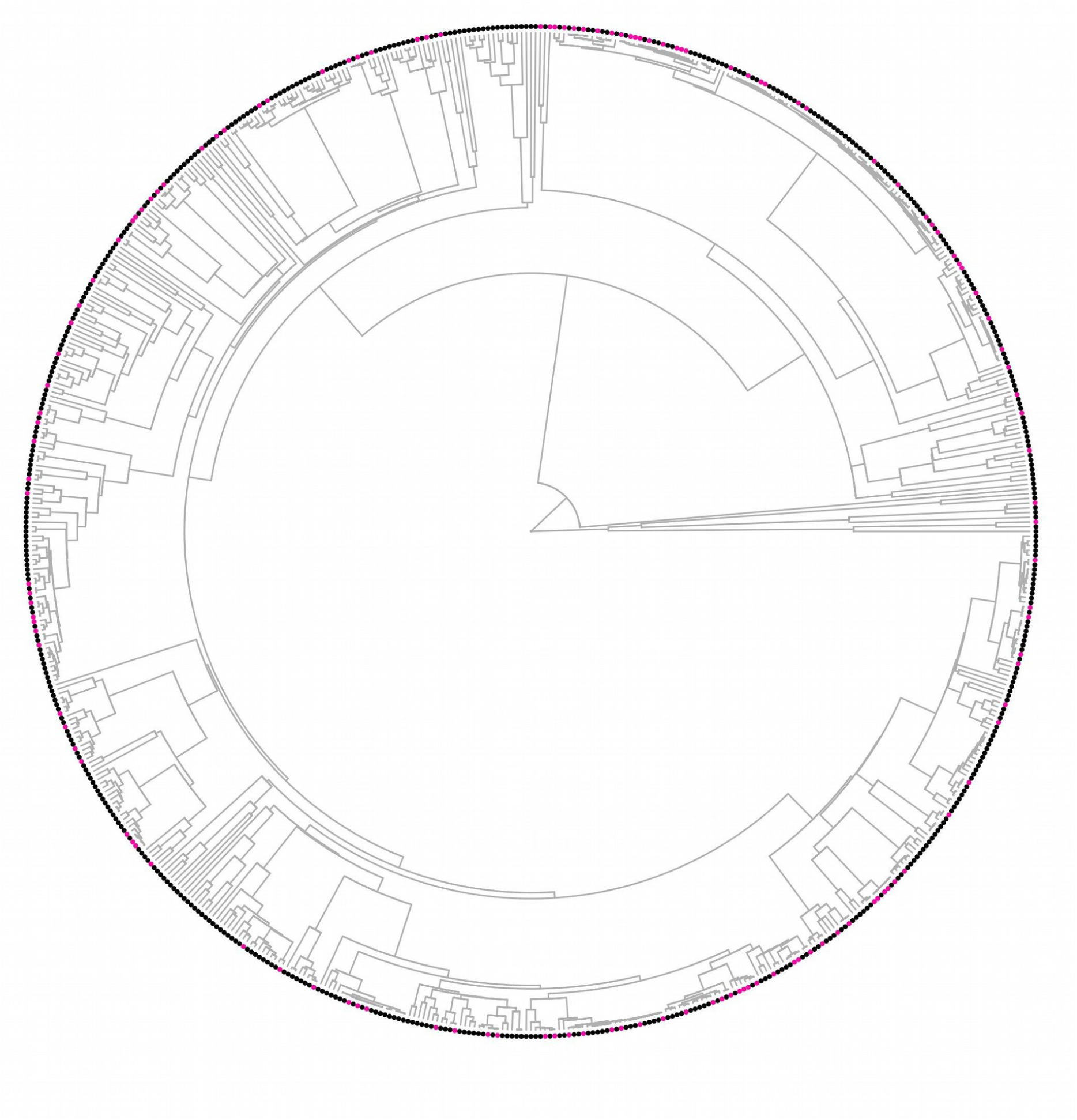
Phylogenetic backbone tree (one example of the 50 replicates). Overall 664 species. 132 species (19.9%, pink dots) that were not present in the backbone phylogeny used to build this tree were randomly inserted into their genera (see methods for details)

**Figure S12.**
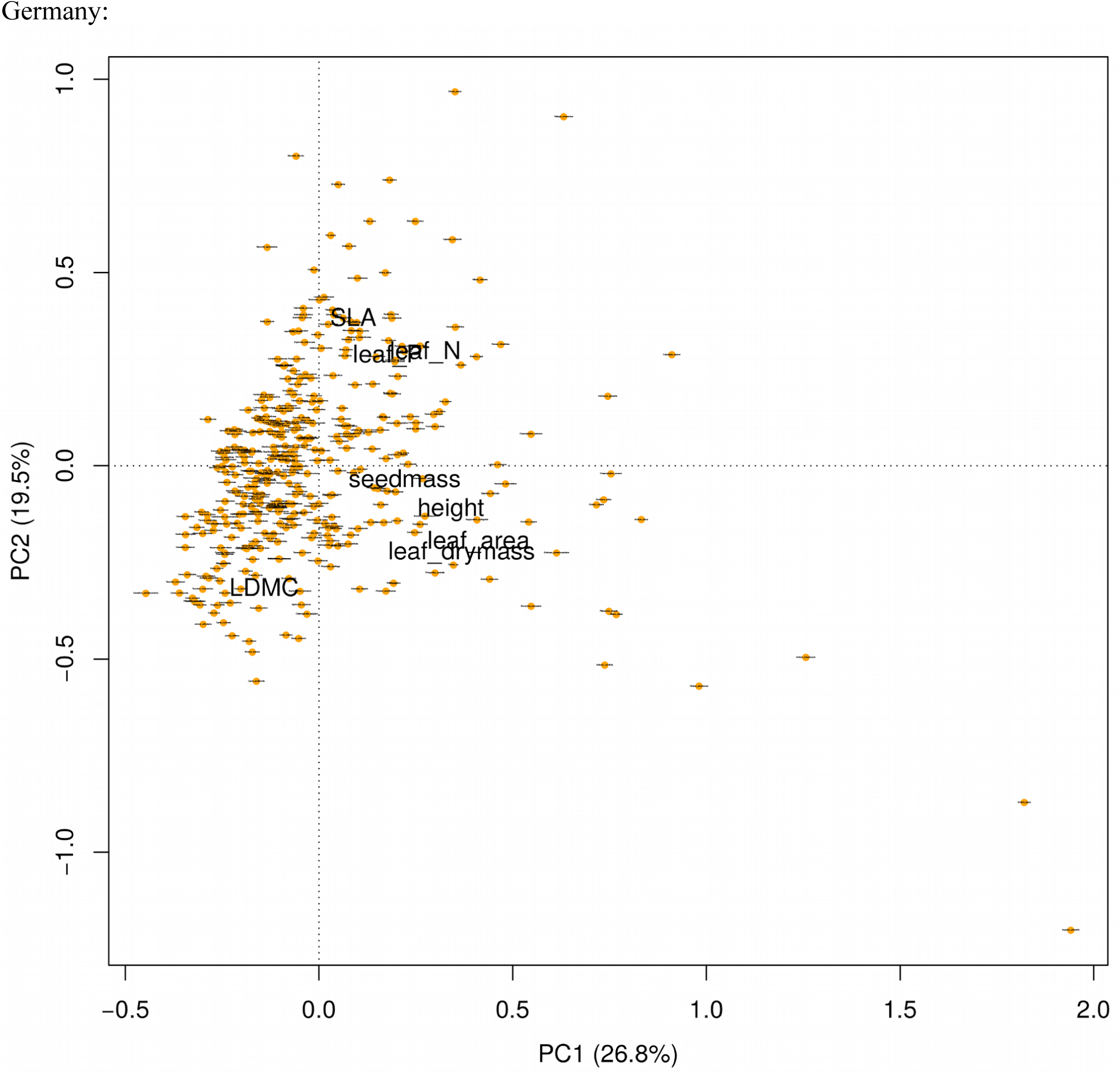
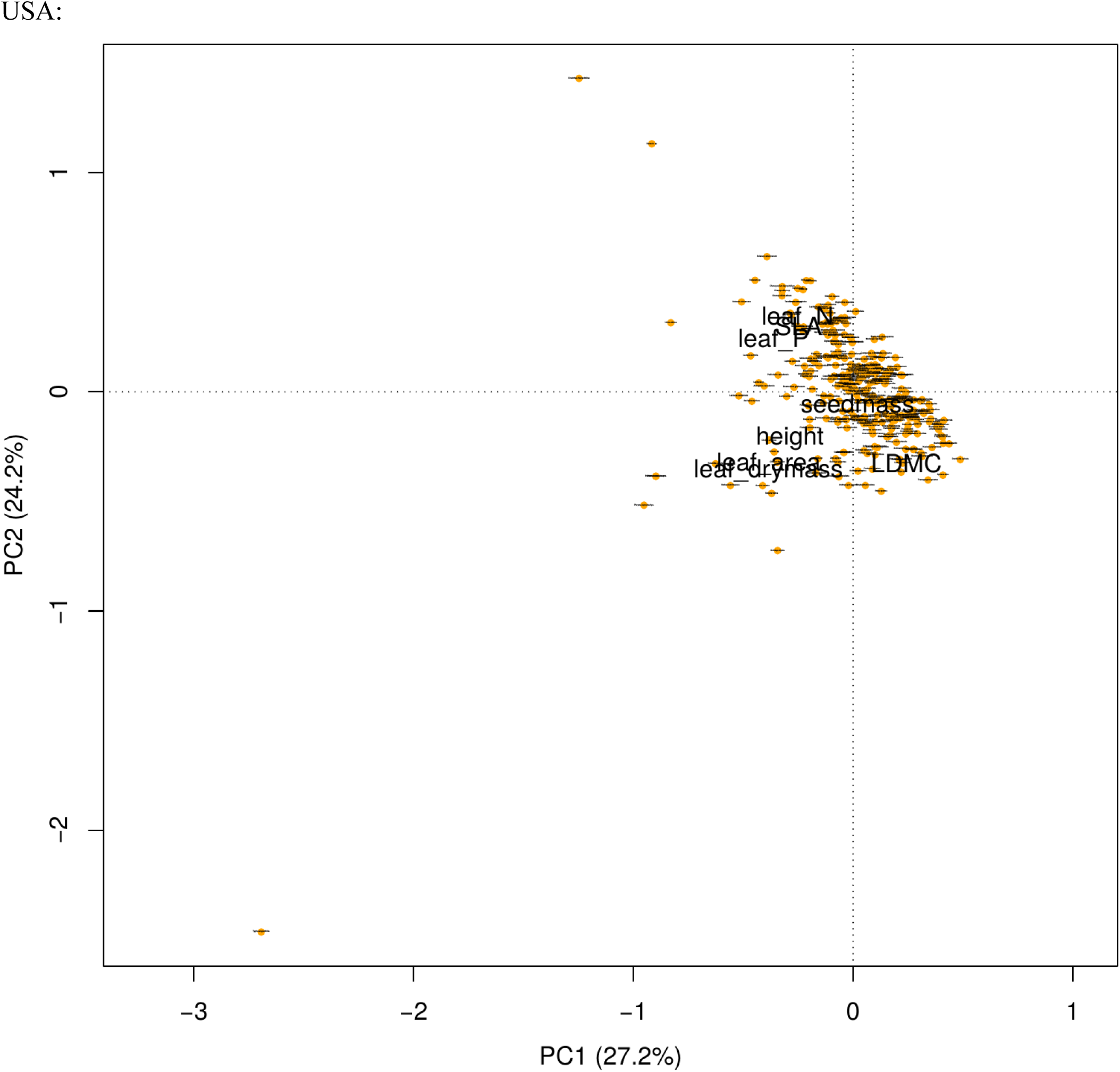
PCA of plant species and their traits for German and US comparison. Each point represents the traits of a single species in the German or US dataset. For obvious outliers, the ability of each species to score such extreme values was individually confirmed e.g. by checking that certain species have unusually large leaf area or leaf nitrogen content. Note that since most of the calculated community properties are relative-abundance weighted, these single outliers do not necessarily have significant impact on the community properties of a given plant community.

**Table S1.** List of German and US datasets for vegetation and ecosystem function data. Ideally, lists project name, main reference, short name used in this paper, number of sites we used, years we have vegetation data for, functions we used including years. Most of the data is openly available in various online repositories (except for data from recent years that are, in some cases, still covered by project-specific embargo periods): Jena Experiment (http://www.the-jena-experiment.de/Data.html), Biodiversity Exploratories (https://www.bexis.uni-jena.de/Login/Account.aspx), Cedar Creek (https://www.cedarcreek.umn.edu/research/data). Data from the Saale grasslands (Jena real world) was provided by Christiane Roscher and is currently not openly available.

| Country | Project name | Project code | Main reference | Number of sites used | Vegetation data years | Ecosystem functions & years |
| --- | --- | --- | --- | --- | --- | --- |
| G | Jena Experiment | Jena Experiment | <sup>14</sup> | 82 | 2003-2015 | plant aboveground biomass (2006-15), plant CN (2007-12), soil organic C (2008, 2011, 2014), root biomass (2011, 2014), herbivory (2010-12), soil microbial biomass C (2010), phosphatase activity (2013), pollinator abundance (2010, 2012) |
| G | Biodiversity Exploratories | German real world | <sup>15</sup> | 150 | 2008-2015 | NA |
| G | Saale grasslands | Jena real world | Roscher unpubl. | 14 | 2011 | NA |
| G | Jena invasion sub-experiments | Jena invasion | <sup>16,17</sup> | 82 | 2003-2015 | NA |
| G | Jena mown succession plots | Jena succession | <sup>16</sup> | 2 | 2003-2009 | NA |
| U | e120 | BioDIV | <sup>18</sup> | 159 | 1996-2015 (not 2009) | plant aboveground biomass (2006-15, not 2009), plant CN (2006), total soil C (2006), root biomass (2010) |
| U | e001 | Fertilization 1 | <sup>19</sup> | 207 | 1982-2004 | NA |
| U | e002 | Fertilization 2 | <sup>19</sup> | 162 | 1982-1991 | NA |
| U | e014 | Old field succession chronosequence | <sup>12</sup> | 23 | 1983, 1989, 1994, 1997, 2002, 2006, 2011 | NA |
| U | e093 | Oak savannah | <sup>11,13</sup> | 30 | 1991 | NA |

**Table S2.** T-test results for differences between German experimental and real-world plots. Welsh t-tests with unequal variances. Full set of 12 community properties averaged across all years per plot for Jena Experiment (82 plots) and combined real-world data (German real world: 150 plots, Jena real world: 14 plots). T-statistic, degrees of freedom (df), experimental (Exp) data mean and real world (RW) data mean are rounded to two, p-values to 5 decimal places.

| model | t_statistic | df | mean_Exp | mean_RW | p_value |
| --- | --- | --- | --- | --- | --- |
| D2 | -10.17 | 133.06 | 3.23 | 7.42 | 0 |
| SEve | 16.35 | 86.69 | 0.65 | 0.27 | 0 |
| FRic | -19.04 | 192.57 | 2.74 | 12.84 | 0 |
| FEve | -5.57 | 83.98 | 0.34 | 0.52 | 0 |
| SLA | -4.54 | 140.03 | 24.26 | 26.44 | 0.00001 |
| leaf_drymass | 2.12 | 93.49 | 61.03 | 48.28 | 0.03662 |
| leaf_N | 1.31 | 96.71 | 25.66 | 24.81 | 0.19315 |
| leaf_P | -1.44 | 109.38 | 2.19 | 2.26 | 0.15242 |
| seedmass | 3.95 | 84.7 | 2.76 | 1.52 | 0.00016 |
| height | -2.36 | 115.5 | 0.45 | 0.5 | 0.01988 |
| LDMC | -5.27 | 98.28 | 0.22 | 0.25 | 0 |
| MNTD | 6.62 | 82.65 | 109.29 | 31.2 | 0 |

**Table S3.** T-test results for differences between US experimental and real-world plots. Welsh t-tests with unequal variances. Full set of 12 community properties averaged across all years per plot for BioDIV (159 plots) and combined real-world data (Nutrient 1 & 2; 207 and 162 plots, respectively). T-statistic, degrees of freedom (df), experimental (Exp) data mean and real world (RW) data mean are rounded to two, p-values to 5 decimal places.

| model | t_statistic | df | mean_Exp | mean_RW | p_value |
| --- | --- | --- | --- | --- | --- |
| D2 | -1.83 | 226.06 | 2.23 | 2.46 | 0.06831 |
| SEve | 18.02 | 188.98 | 0.71 | 0.38 | 0 |
| FRic | -7.59 | 243.07 | 1.4 | 2.62 | 0 |
| FEve | -2.73 | 176.06 | 0.29 | 0.35 | 0.00693 |
| SLA | -8.01 | 204.34 | 14.77 | 16.5 | 0 |
| leaf_drymass | 10 | 193.36 | 74.23 | 45.67 | 0 |
| leaf_N | -4.9 | 237.85 | 16.97 | 18.77 | 0 |
| leaf_P | -6.68 | 248.46 | 1.55 | 1.71 | 0 |
| seedmass | 5.21 | 169.51 | 3.88 | 2.18 | 0 |
| height | 3.37 | 193.75 | 0.77 | 0.7 | 0.0009 |
| LDMC | -2.57 | 169.1 | 0.3 | 0.32 | 0.01105 |
| MNTD | 4.71 | 169.23 | 97.1 | 50.95 | 0.00001 |

**Table S4.**
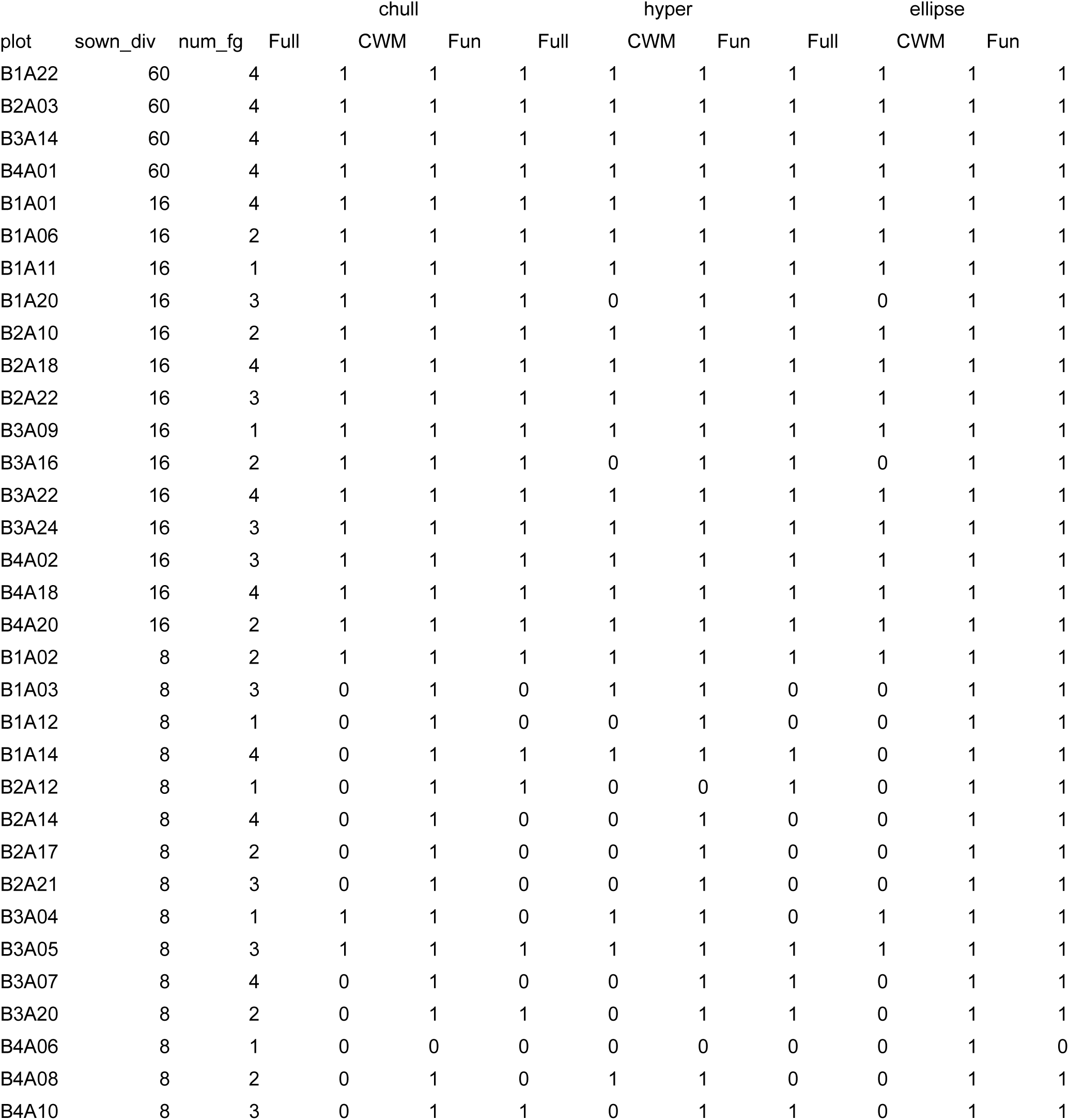
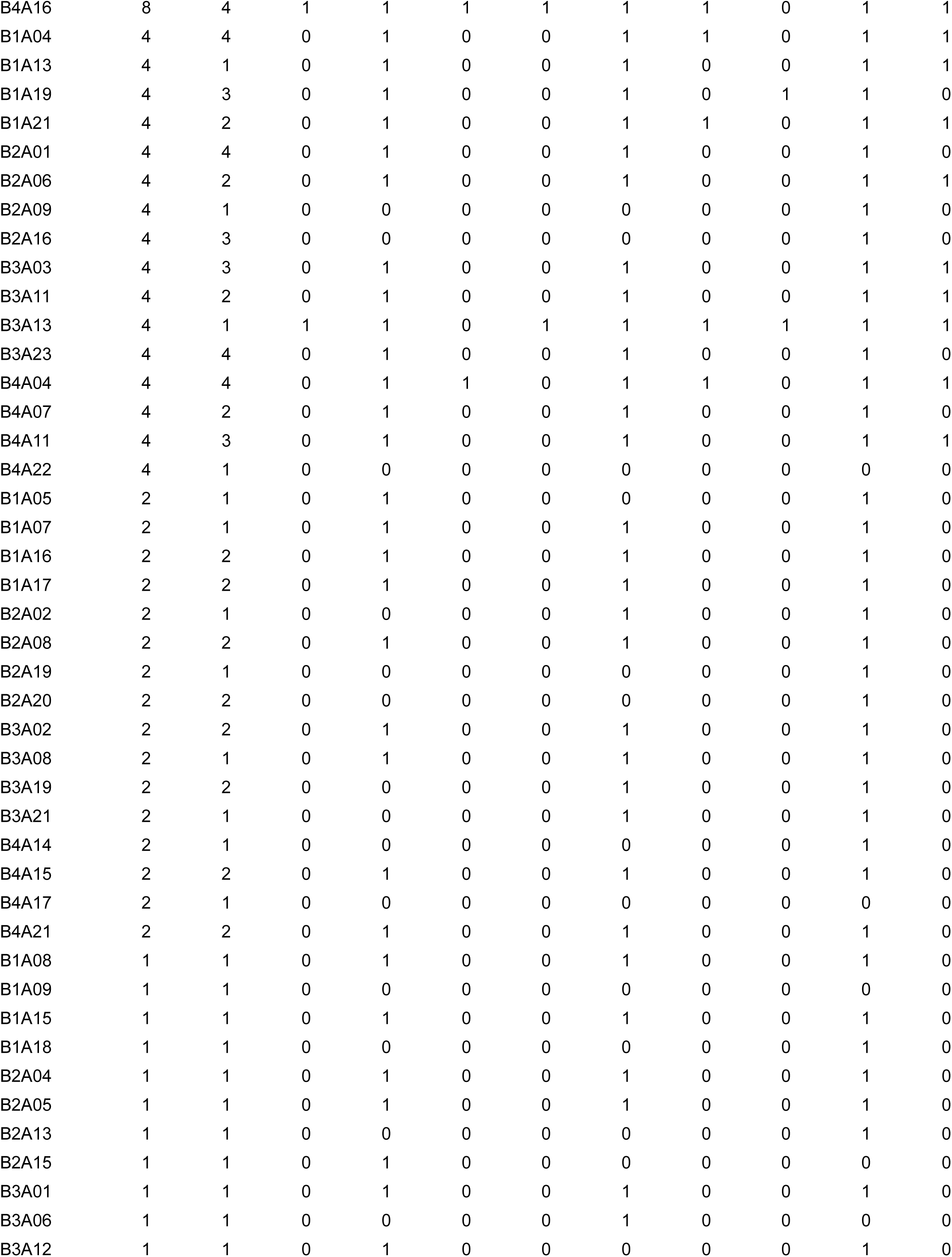
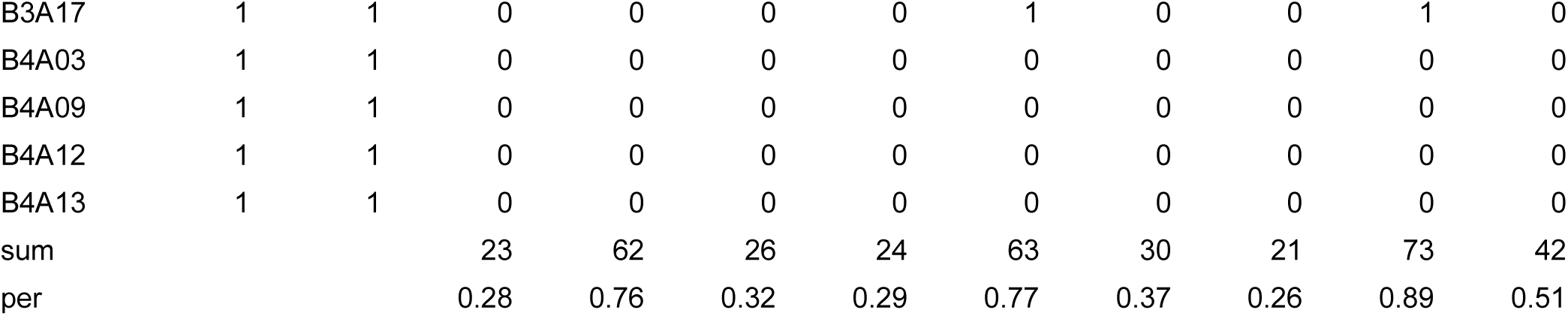
Jena Experiment plots with their sown diversity (sown_div), number of functional groups (num_fg) and their selection as realistic plots (1) based on three different methods of calculating the intersection and based on three different subsets of community properties entering the PCA’s. Methods: Intersection of three-dimensional convex hull volumes (chull), hypervolumes (hyper) and 95% confidence ellipses (ellipse). Subsets: Full (all 12 community properties), CWM (8 community weighted means) and Fun (4 functional diversity metrics). Additionally, the number of realistic plots (sum) and the percentage (per) of realistic plots from the overall number of plots (82) are given for each combination of methodology and community property subset. Plots are sorted by sown diversity levels.

**Table S5.**
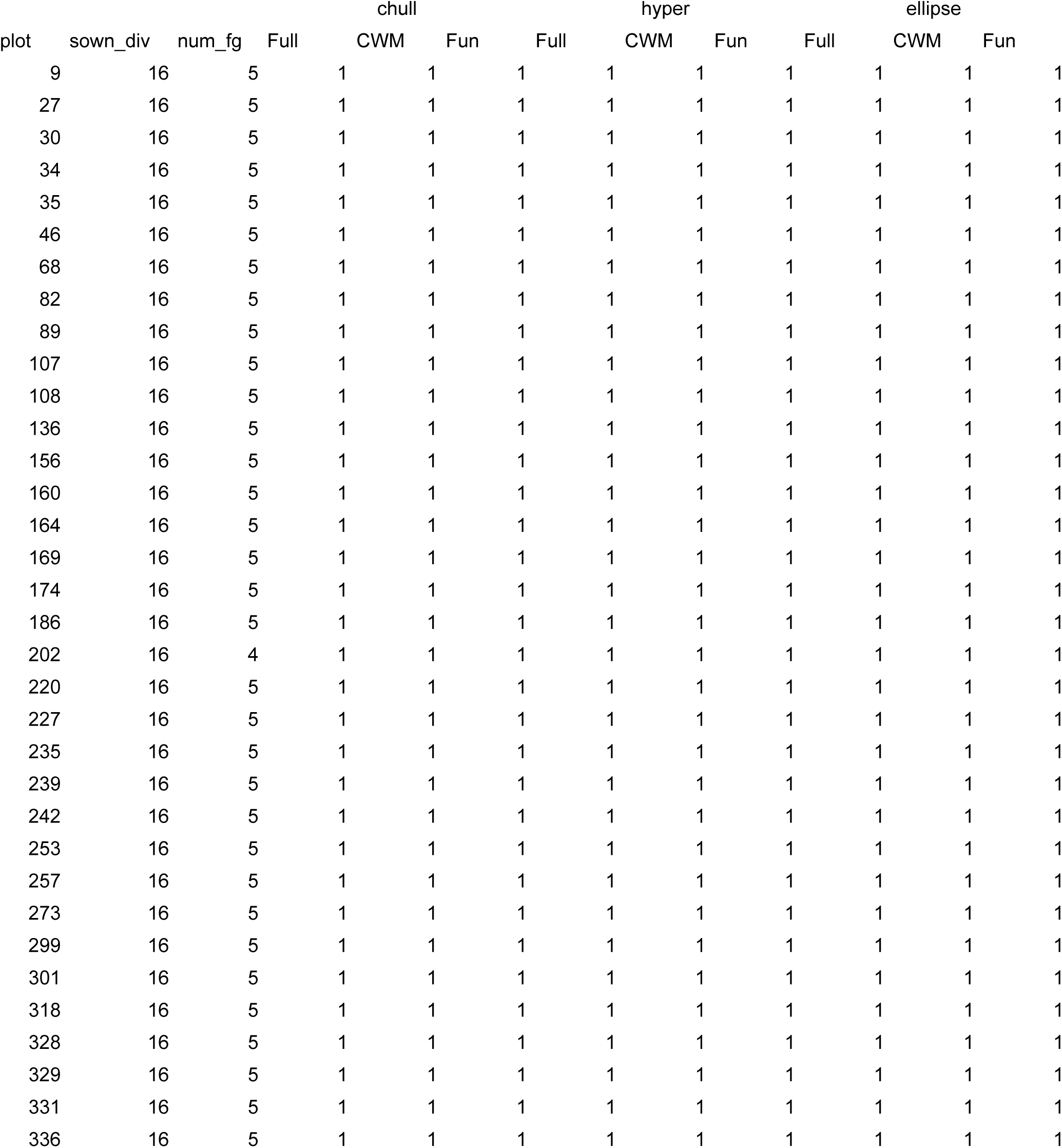
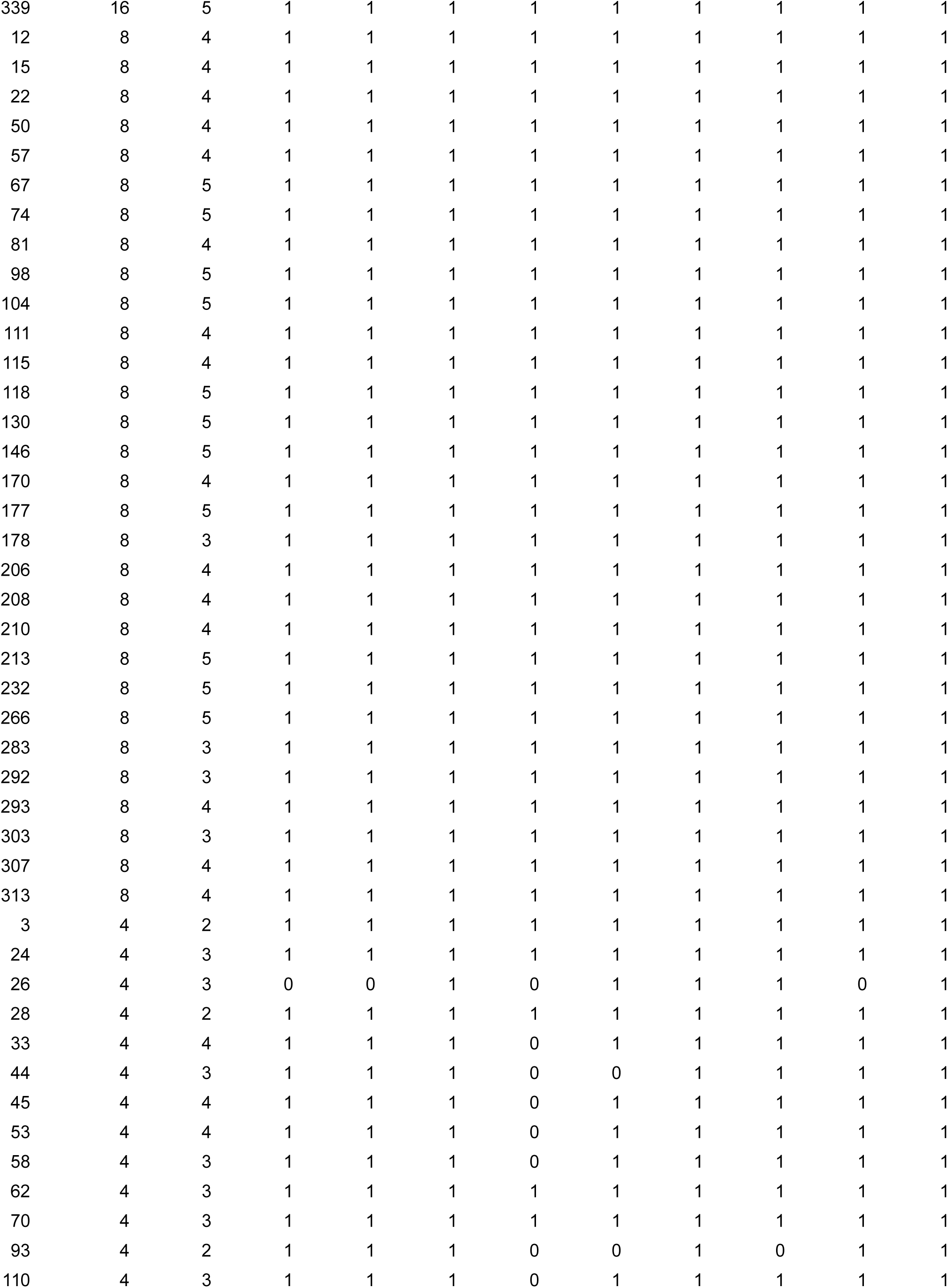
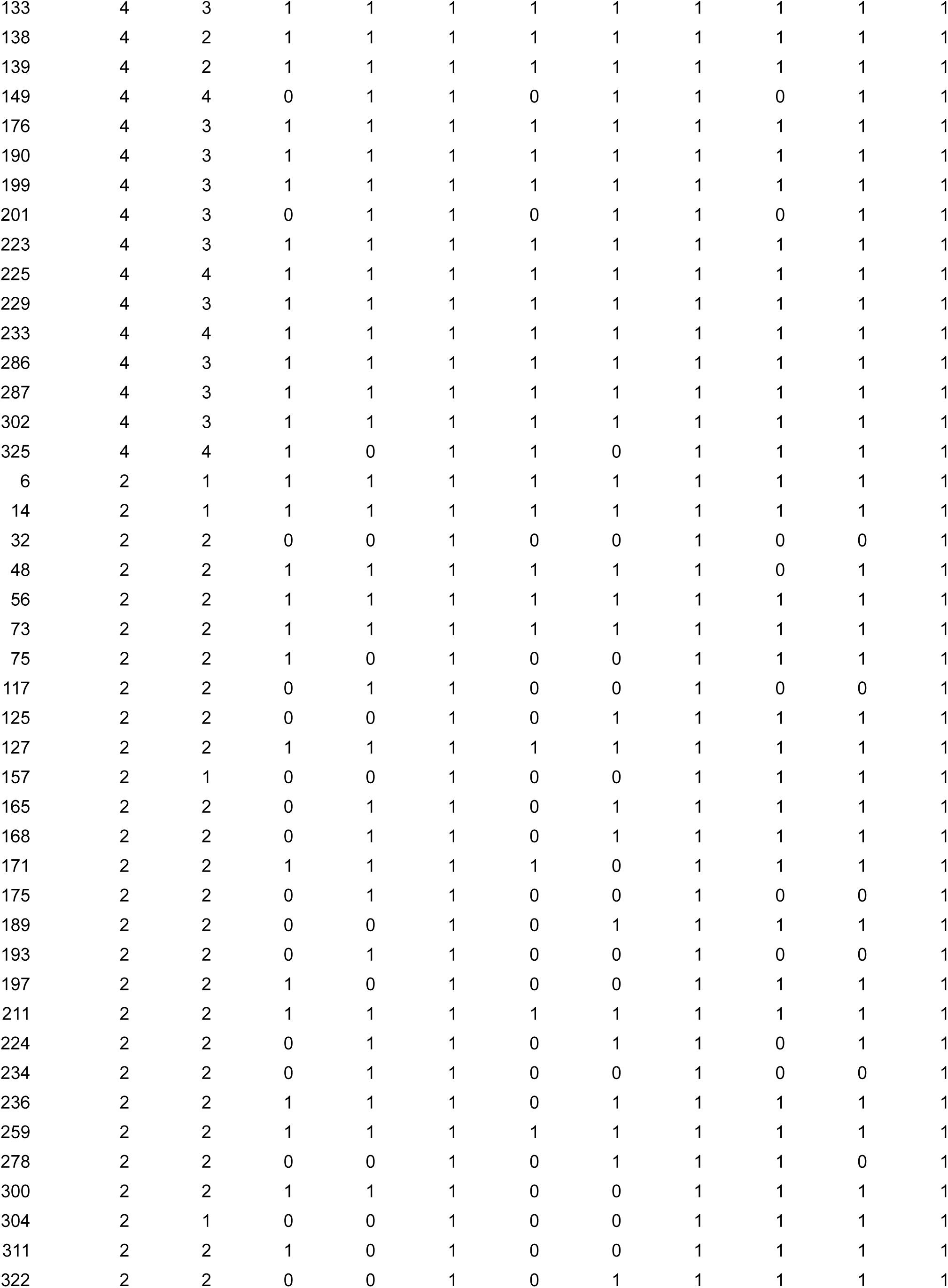
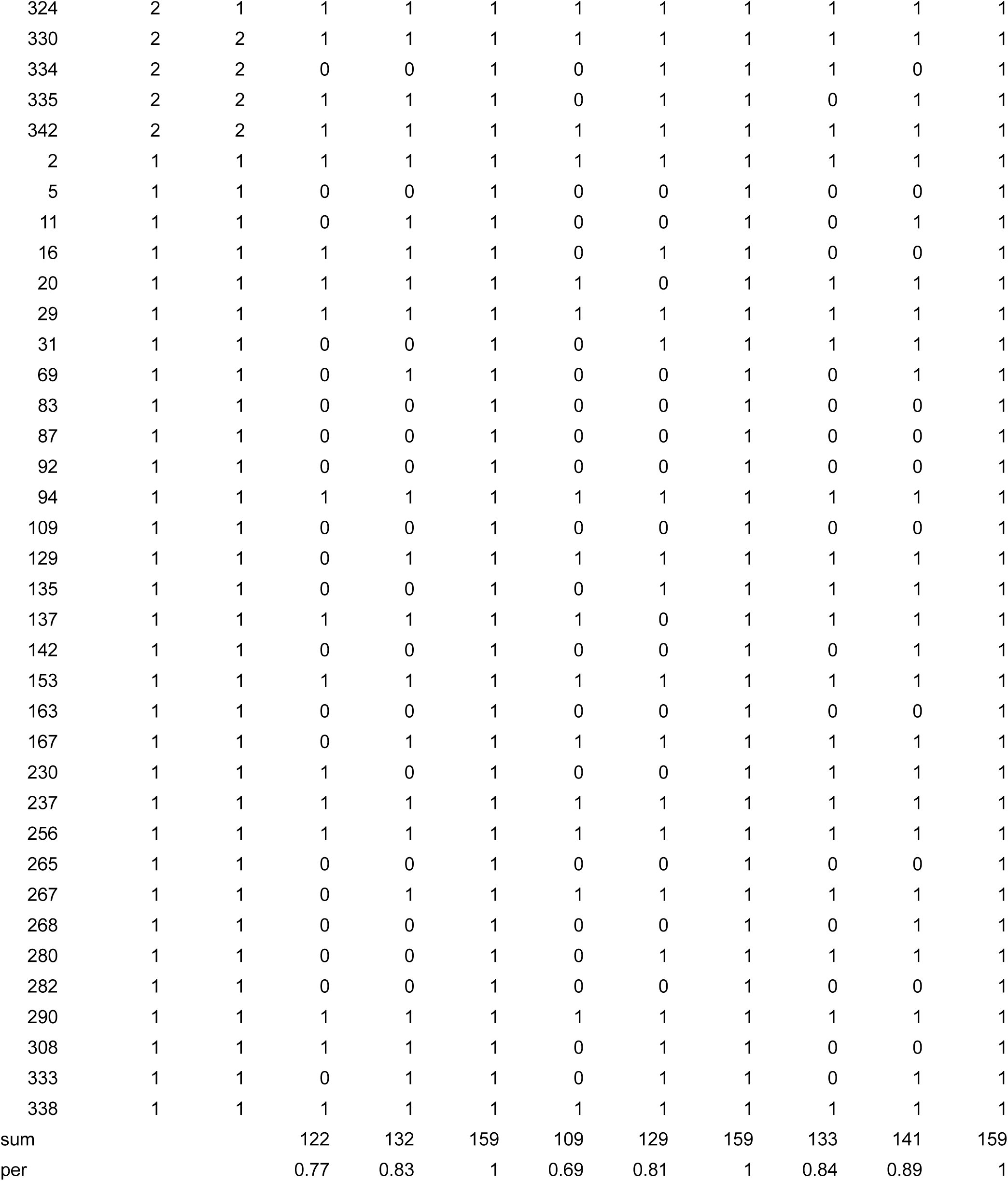
BioDIV plots with their sown diversity (sown_div), number of functional groups (num_fg) and their selection as realistic plots (1) based on three different methods of calculating the intersection and based on three different subsets of community properties entering the PCA’s. Methods: Intersection of three-dimensional convex hull volumes (chull), hypervolumes (hyper) and 95% confidence ellipses (ellipse). Subsets: Full (all 16 community properties), CWM (9 community weighted means) and Fun (4 functional diversity metrics). Additionally, the number of realistic plots (sum) and the percentage (per) of realistic plots from the overall number of plots (159) are given for each combination of methodology and community property subset.

**Table S6.** T-test results for differences between realistic and unrealistic plots for the Jena Experiment. Welsh t-tests with unequal variances. Realistic plots were calculated based on the full set of community properties and the convex hull volume method. All properties were averaged across all available years per plot (23 realistic and 59 unrealistic plots). T-statistic, degrees of freedom (df), means of realistic (real) and unrealistic communities (unreal) are rounded to two, p-values to four decimal places.

| model | t_statistic | df | mean_real | mean_unreal | p_value |
| --- | --- | --- | --- | --- | --- |
| sowndiv | 4.77 | 22.34 | 21.74 | 3.46 | 0.0001 |
| numfg | 3.54 | 36.95 | 2.83 | 1.85 | 0.0011 |
| D2 | 5.06 | 22.72 | 6.65 | 1.9 | 0 |
| SEve | -10.33 | 78.5 | 0.45 | 0.74 | 0 |
| FRic | 8.25 | 24.88 | 7.32 | 0.96 | 0 |
| FEve | 6.46 | 74.97 | 0.54 | 0.27 | 0 |
| SLA | 0.63 | 49.5 | 24.64 | 24.11 | 0.5334 |
| leaf_drymass | -0.55 | 78.6 | 57.26 | 62.5 | 0.5845 |
| leaf_N | 0.3 | 78.09 | 25.88 | 25.58 | 0.7679 |
| leaf_P | 2.24 | 77.01 | 2.31 | 2.14 | 0.0281 |
| seedmass | -0.35 | 79.48 | 2.64 | 2.81 | 0.7303 |
| height | 1.87 | 61.58 | 0.5 | 0.42 | 0.0663 |
| LDMC | 1.75 | 42.61 | 0.23 | 0.21 | 0.0865 |
| MNTD | -3.56 | 77.95 | 64.82 | 126.63 | 0.0006 |

**Table S7.** T-test results for differences between realistic and unrealistic plots for BioDIV. Welsh t-tests with unequal variances. Realistic plots were calculated based on the full set of community properties and the convex hull volume method. All properties were averaged across all available years per plot (122 realistic and 37 unrealistic plots). T-statistic, degrees of freedom (df), means of realistic (real) and unrealistic communities (unreal) are rounded to two, p-values to four decimal places.

| model | t_statistic | df | mean_real | mean_unreal | p_value |
| --- | --- | --- | --- | --- | --- |
| sowndiv | 11.51 | 137.13 | 7.81 | 1.65 | 0 |
| numfg | 10.95 | 122.28 | 3.49 | 1.54 | 0 |
| D2 | 11.23 | 133.74 | 2.58 | 1.09 | 0 |
| SEve | -10.43 | 102.79 | 0.64 | 0.92 | 0 |
| FRic | 10.7 | 121 | 1.82 | 0 | 0 |
| FEve | 14.65 | 121 | 0.38 | 0 | 0 |
| SLA | 1.04 | 46.22 | 14.91 | 14.3 | 0.3043 |
| leaf_drymass | -1.27 | 46.88 | 71.95 | 81.76 | 0.2119 |
| leaf_N | 1.2 | 42.28 | 17.27 | 15.99 | 0.2366 |
| leaf_P | 1.42 | 46.02 | 1.57 | 1.48 | 0.1633 |
| seedmass | 0.15 | 41.7 | 3.92 | 3.76 | 0.8815 |
| height | -1.81 | 40.78 | 0.75 | 0.87 | 0.0773 |
| LDMC | -0.66 | 44.55 | 0.3 | 0.31 | 0.5149 |
| MNTD | 1.95 | 55.87 | 107.8 | 61.82 | 0.056 |

**Table S8.** Model parameters for BEF relationships presented in Fig. 2. Values are presented for unconstrained (u) and constrained (c) models of Jena (J) and BioDIV BEF relationships. Constraining was done using all 12 community properties and the convex hull method. Sample size (n), slope estimates (slop), lower (low) and upper (upp) 95% confidence intervals, p-values (p) and adjusted R^2^ values (R2). All values are rounded to two decimal places.

| mod | u_n | c_n | u_slop | c_slop | u_low | c_low | u_upp | c_upp | u_p | c_p | u_R2 | c_R2 |
| --- | --- | --- | --- | --- | --- | --- | --- | --- | --- | --- | --- | --- |
| J_biomass | 82.00 | 23.00 | 2.39 | 2.03 | 1.87 | 0.45 | 2.91 | 3.61 | 0.00 | 0.01 | 0.51 | 0.22 |
| J_plantCN | 82.00 | 23.00 | 1.02 | -2.12 | -0.15 | -5.92 | 2.20 | 1.67 | 0.09 | 0.26 | 0.02 | 0.02 |
| J_soilorgC | 82.00 | 23.00 | 0.09 | 0.06 | 0.04 | -0.10 | 0.13 | 0.23 | 0.00 | 0.42 | 0.14 | -0.01 |
| J_rootbiomass | 80.00 | 23.00 | 0.10 | 0.04 | 0.07 | -0.05 | 0.14 | 0.13 | 0.00 | 0.37 | 0.34 | -0.01 |
| J_herbivory | 80.00 | 23.00 | 0.07 | 0.24 | 0.02 | 0.11 | 0.12 | 0.38 | 0.01 | 0.00 | 0.08 | 0.36 |
| J_micBMC | 80.00 | 23.00 | 48.40 | -8.18 | 23.35 | -92.06 | 73.46 | 75.71 | 0.00 | 0.84 | 0.15 | -0.05 |
| J_Jphosphatase | 80.00 | 23.00 | 103.17 | 16.33 | 70.38 | -98.35 | 135.97 | 131.01 | 0.00 | 0.77 | 0.33 | -0.04 |
| J_pollinators | 79.00 | 23.00 | 0.10 | 0.13 | 0.06 | 0.03 | 0.14 | 0.23 | 0.00 | 0.02 | 0.27 | 0.21 |
| BioDIV_biomass | 159.00 | 122.00 | 0.24 | 0.26 | 0.20 | 0.22 | 0.28 | 0.31 | 0.00 | 0.00 | 0.44 | 0.49 |
| BioDIV_plantCN | 158.00 | 122.00 | -0.20 | 1.22 | -1.24 | 0.16 | 0.84 | 2.28 | 0.71 | 0.02 | -0.01 | 0.03 |
| BioDIV_soilC | 158.00 | 122.00 | 0.03 | 0.03 | 0.01 | 0.01 | 0.04 | 0.05 | 0.00 | 0.00 | 0.08 | 0.08 |
| BioDIV_rootbiomass | 150.00 | 117.00 | 261.03 | 266.43 | 221.71 | 216.44 | 300.35 | 316.41 | 0.00 | 0.00 | 0.53 | 0.49 |

**Table S9.**
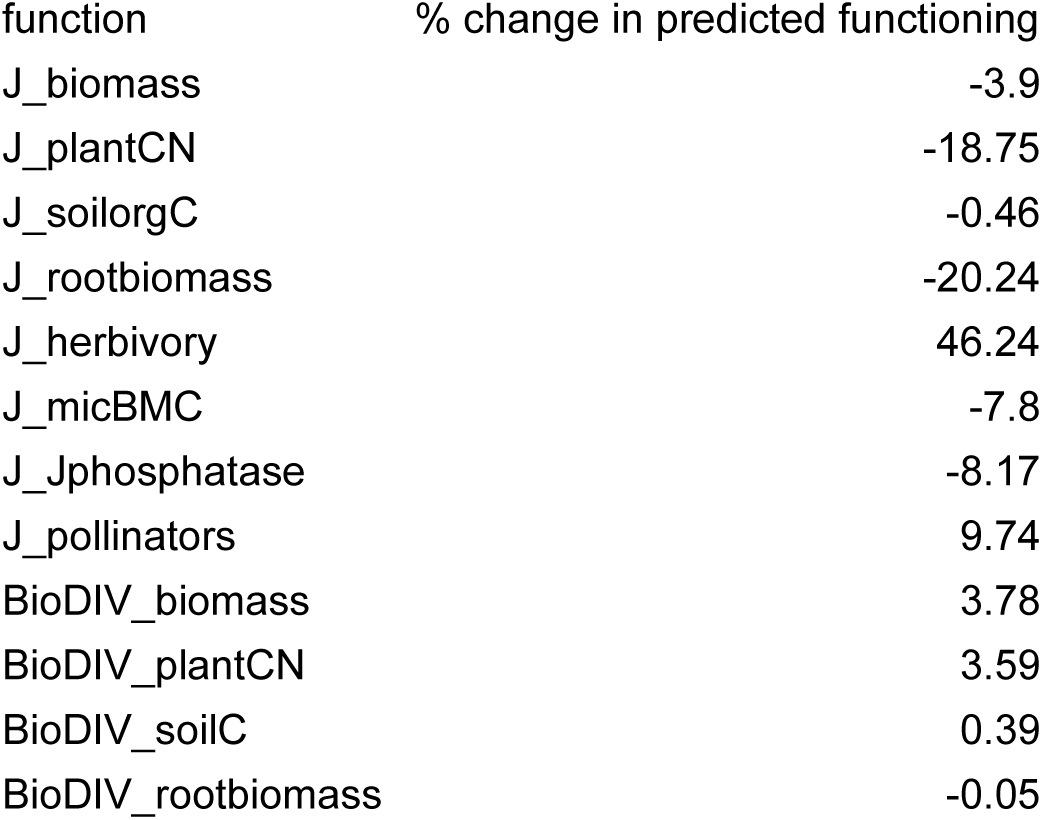
Constraining-related change in functioning at maximum species richness. For each of the 12 BEF relationships from the Jena Experiment (J) and BioDIV presented in Fig. 2, the table shows the constraining-related percentage change in the model-predicted function variable at maximum species richness (the proportional difference in the un-transformed function value at the right-hand tip of the black and red lines in Fig. 2). The average absolute percentage function change is 10.3% (SE: 4%).

**Table S10.** Differences between range in function unconstrained and constrained data underlying the BEF relationships in Fig. 2. Values are presented for unconstrained and constrained datasets of Jena (J) and BioDIV BEF relationships. Constraining was done using all 12 community properties and the convex hull method. Ranges were calculated based maximum and minimum function performance in unconstrained and constrained datasets. Range changes were calculated as the proportion of unconstrained functioning still covered by constrained functioning. Changes are caused by the removal of unrealistic plots which changes the distribution of function values for a given species richness level, but also by the reduction of the species richness gradient that is caused by the removal of plots. The across-year species richness gradient in Jena changed from 1-35.2 species (unconstrained) to 3.7-35.2 species (constrained). The BioDIV species richness gradient was 1-11.1 species and did not change from unconstrained to constrained datasets.

| model_name | uncon_range | con_range | range_change |
| --- | --- | --- | --- |
| J_biomass | 22 | 9.7 | 0.44 |
| J_plantCN | 34.59 | 24.33 | 0.7 |
| J_soilorgC | 1.54 | 1.26 | 0.82 |
| J_rootbiomass | 1.06 | 0.66 | 0.63 |
| J_herbivory | 1.72 | 1.05 | 0.61 |
| J_micBMC | 800.54 | 624.56 | 0.78 |
| J_Jphosphatase | 1159.23 | 956.63 | 0.83 |
| J_pollinators | 1.1 | 0.82 | 0.74 |
| BioDIV_biomass | 2.91 | 2.38 | 0.82 |
| BioDIV_plantCN | 38.04 | 34.59 | 0.91 |
| BioDIV_soilC | 0.62 | 0.62 | 1 |
| BioDIV_rootbiomass | 1952.87 | 1952.87 | 1 |
| Jena_avg |  |  | 0.69 |
| BioDIV_avg |  |  | 0.93 |
| Overall_avg |  |  | 0.77 |

**Table S11.**
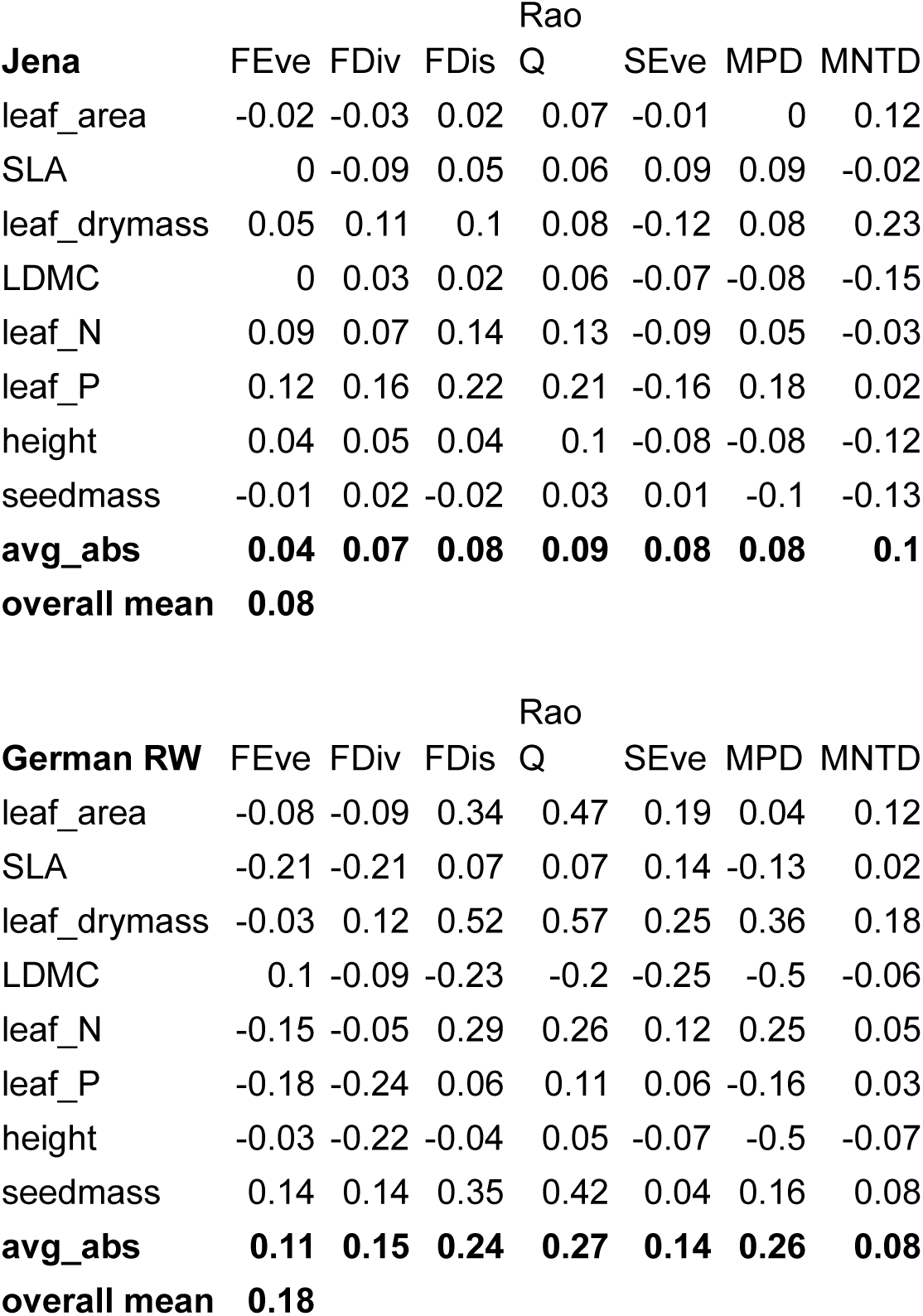
Correlation coefficients for CWM’s versus functional, phylogenetic metrics and evenness, German dataset. Pearson correlation coefficients for Jena Experiment (upper part) and combined German real world community properties (lower part). Bold values are mean absolute correlation coefficients for the columns, the overall mean is the absolute mean across all column averages.

**Table S12.**
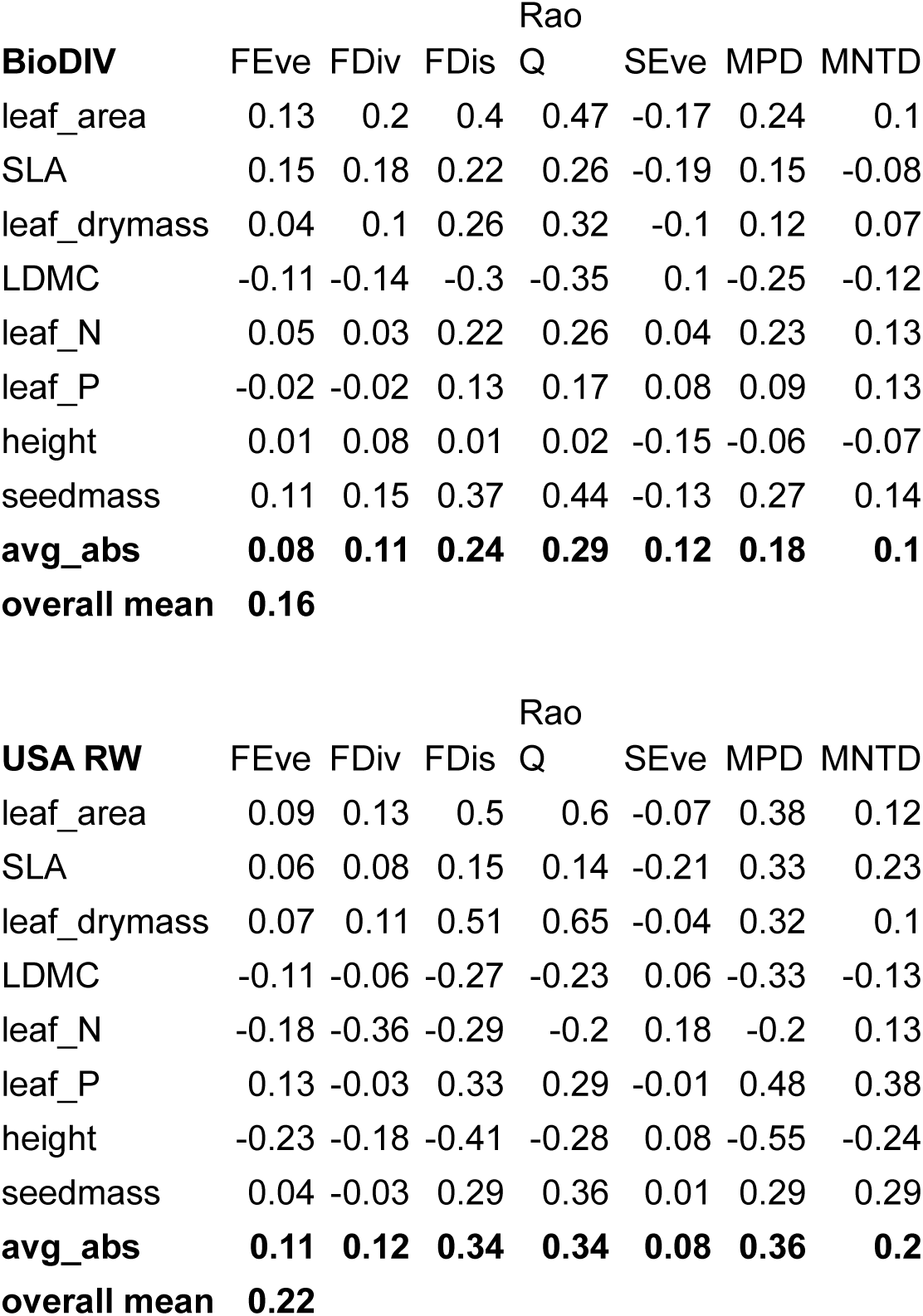
Correlation coefficients for CWM’s versus functional, phylogenetic metrics and evenness, US dataset. Pearson correlation coefficients for BioDIV (upper part) and combined US real world community properties (lower part). Bold values are mean absolute correlation coefficients for the columns, the overall mean is the absolute mean across all column averages.

**Table S13.** PCA scores for full 12 community properties on PCA’s in Fig. 1. Scores have been produced using the scores() command of the “vegan” package^22^ in R and have been rounded to 2 decimal places.

|  | Germany |  | USA |  |
| --- | --- | --- | --- | --- |
|  | PC1 | PC2 | PC1 | PC2 |
| FRic | -3.47 | 0.28 | 2.90 | -3.24 |
| FEve | -2.60 | 0.54 | 2.19 | -2.33 |
| SLA | -0.93 | 2.47 | 3.19 | -0.14 |
| leaf_drymass | 0.69 | 0.84 | 0.76 | 1.50 |
| LDMC | -1.45 | -2.53 | -3.32 | -1.86 |
| leaf_N | 0.24 | 3.15 | 2.18 | 3.21 |
| leaf_P | -0.67 | 2.58 | 3.69 | 1.41 |
| height | -0.93 | -0.91 | -2.71 | 0.22 |
| seedmass | 1.04 | 1.01 | 2.80 | 3.04 |
| D2 | -2.53 | 0.32 | 3.13 | -2.39 |
| SEve | 3.16 | -0.18 | -2.16 | 3.15 |
| MNTD | 2.22 | 0.39 | 0.83 | 2.24 |

**Table S14.** Full dataset of community properties for all plots used in the PCA’s over all years (submitted along with R-code at first submission).

**Table S15.**
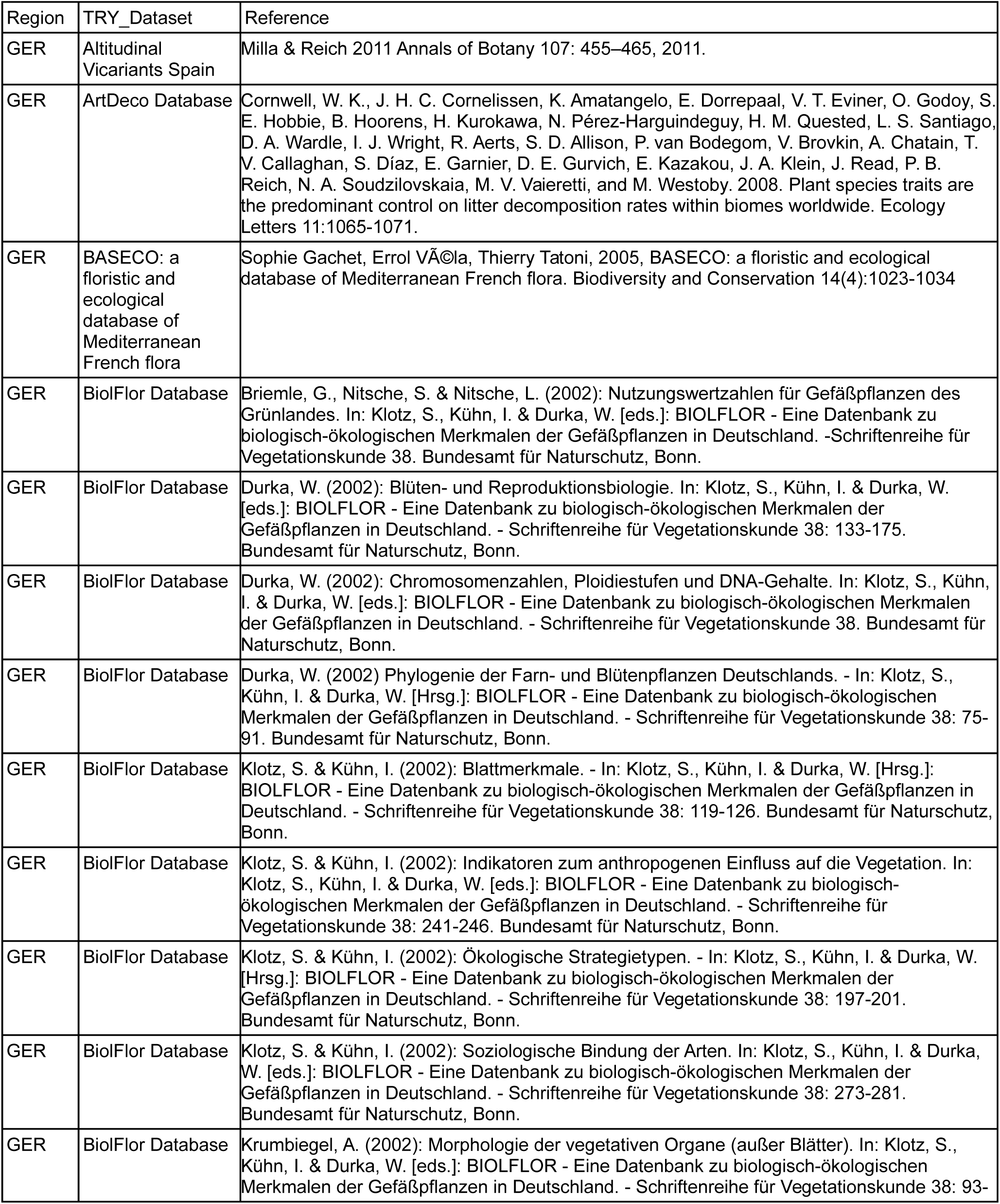
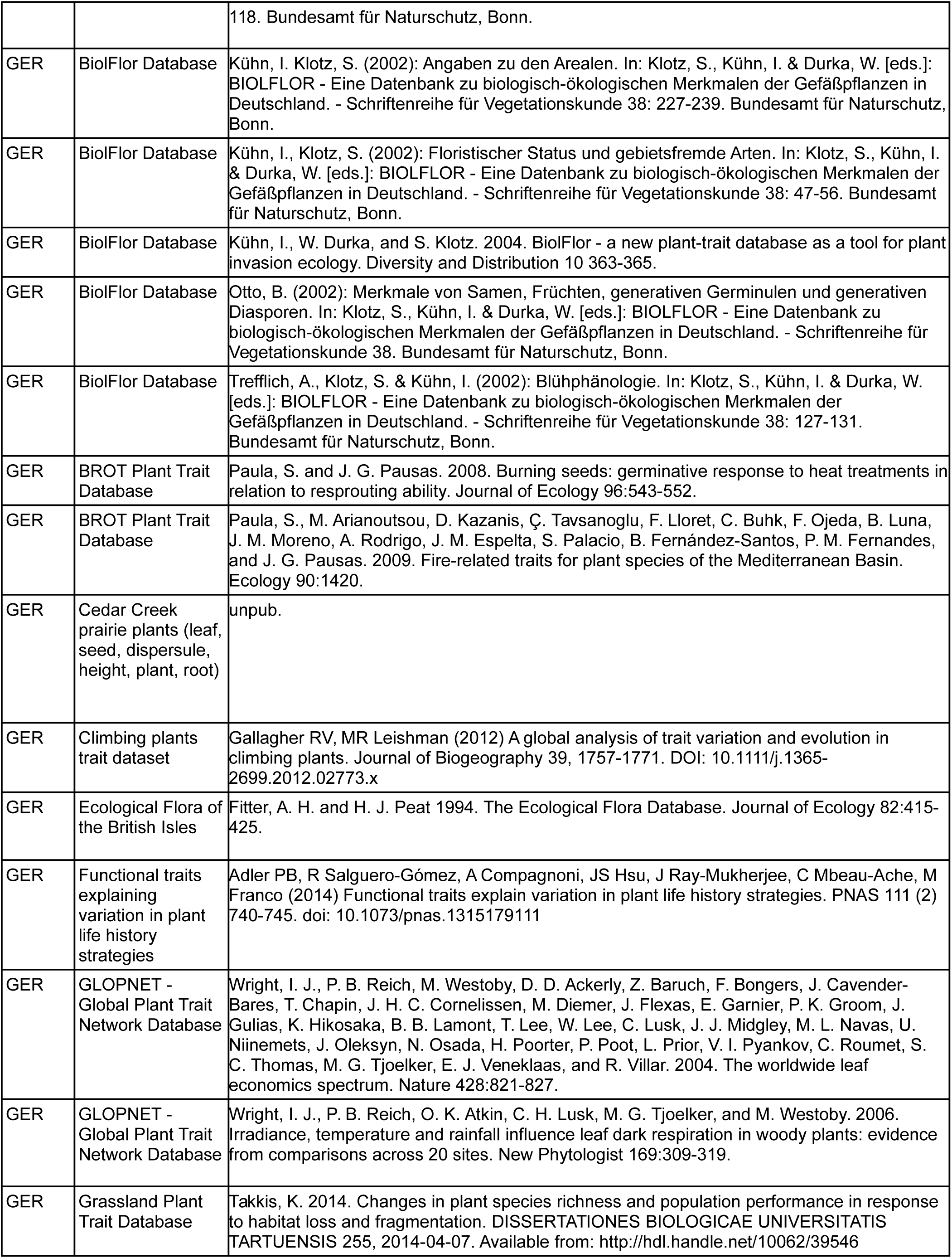
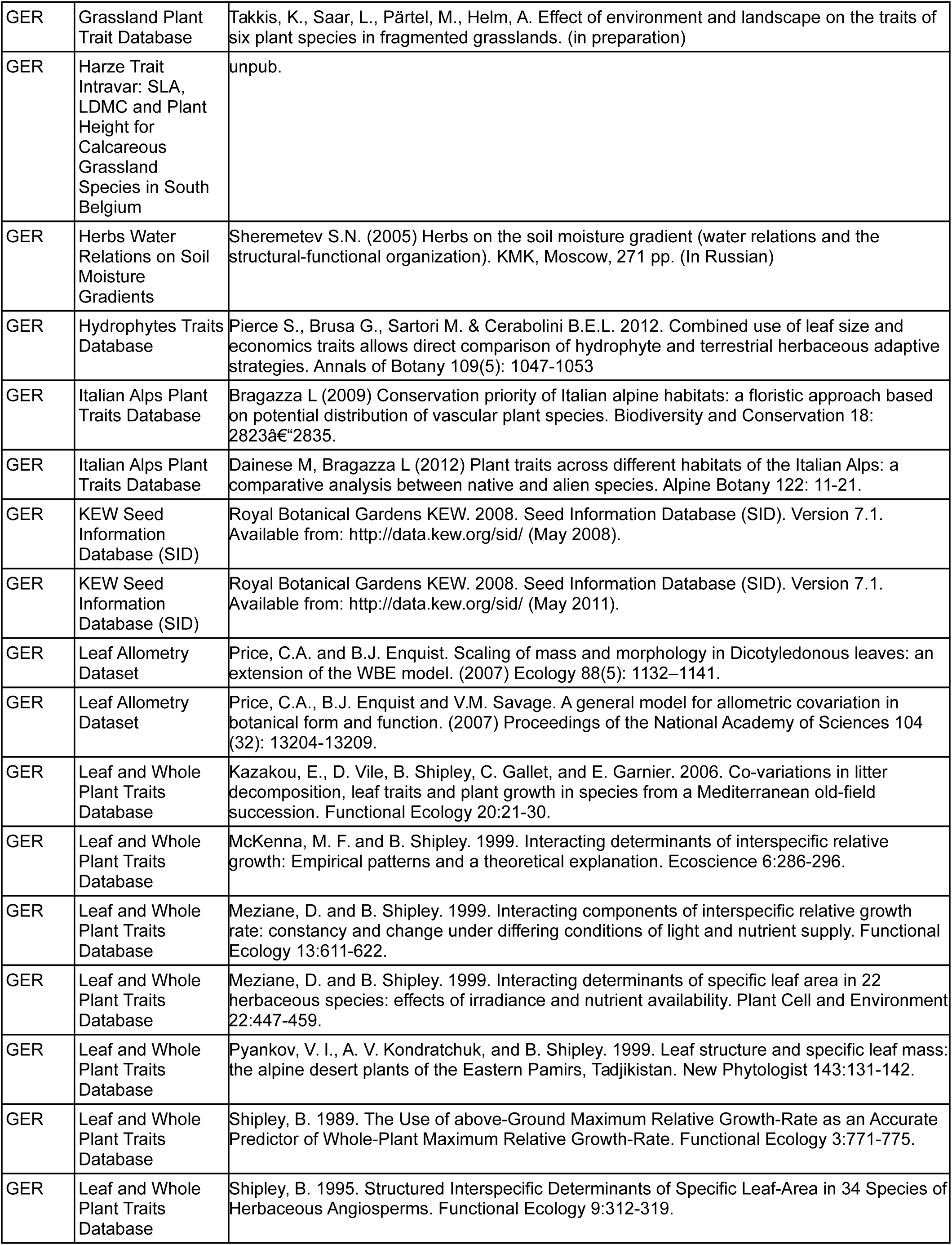
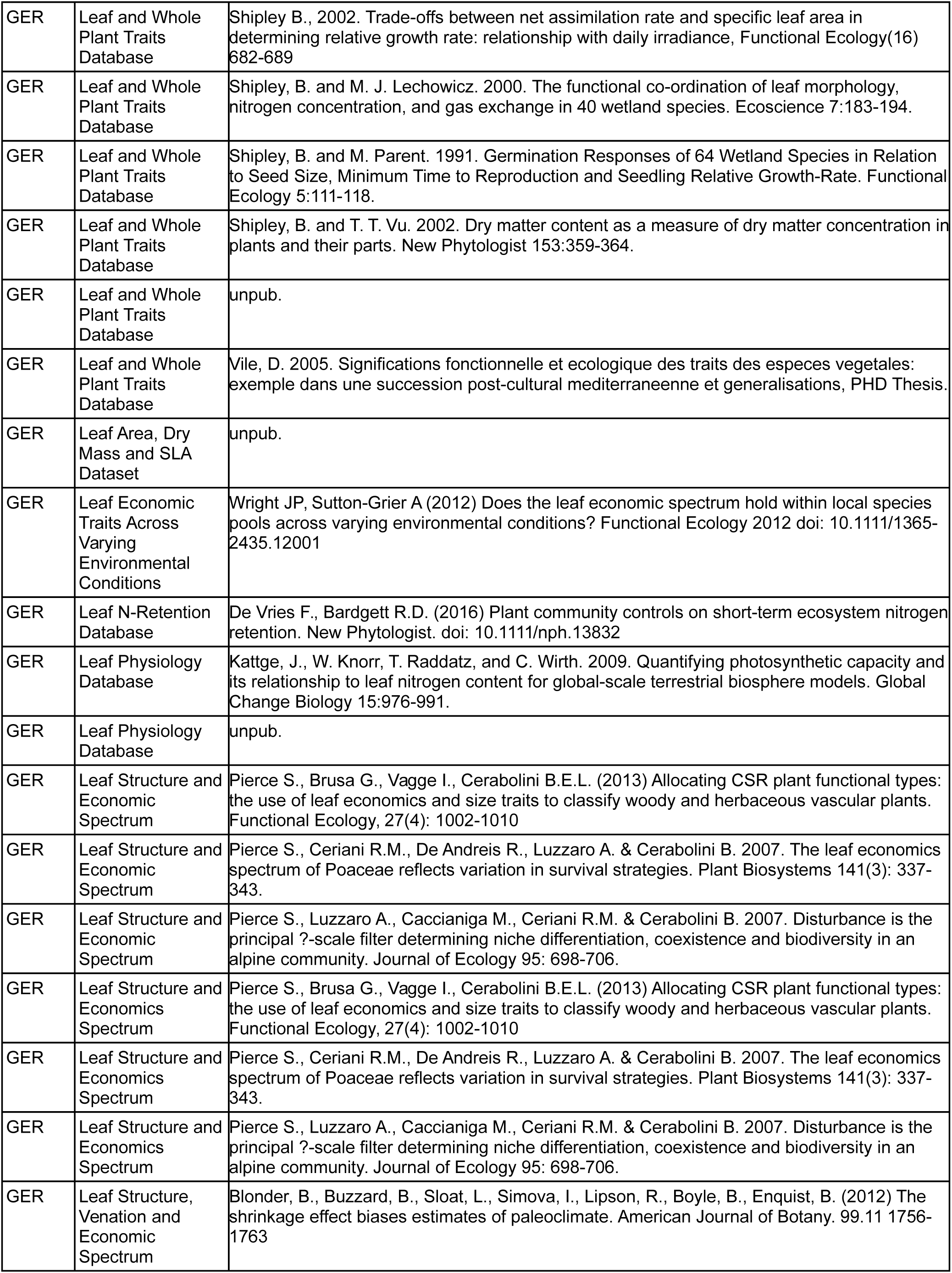
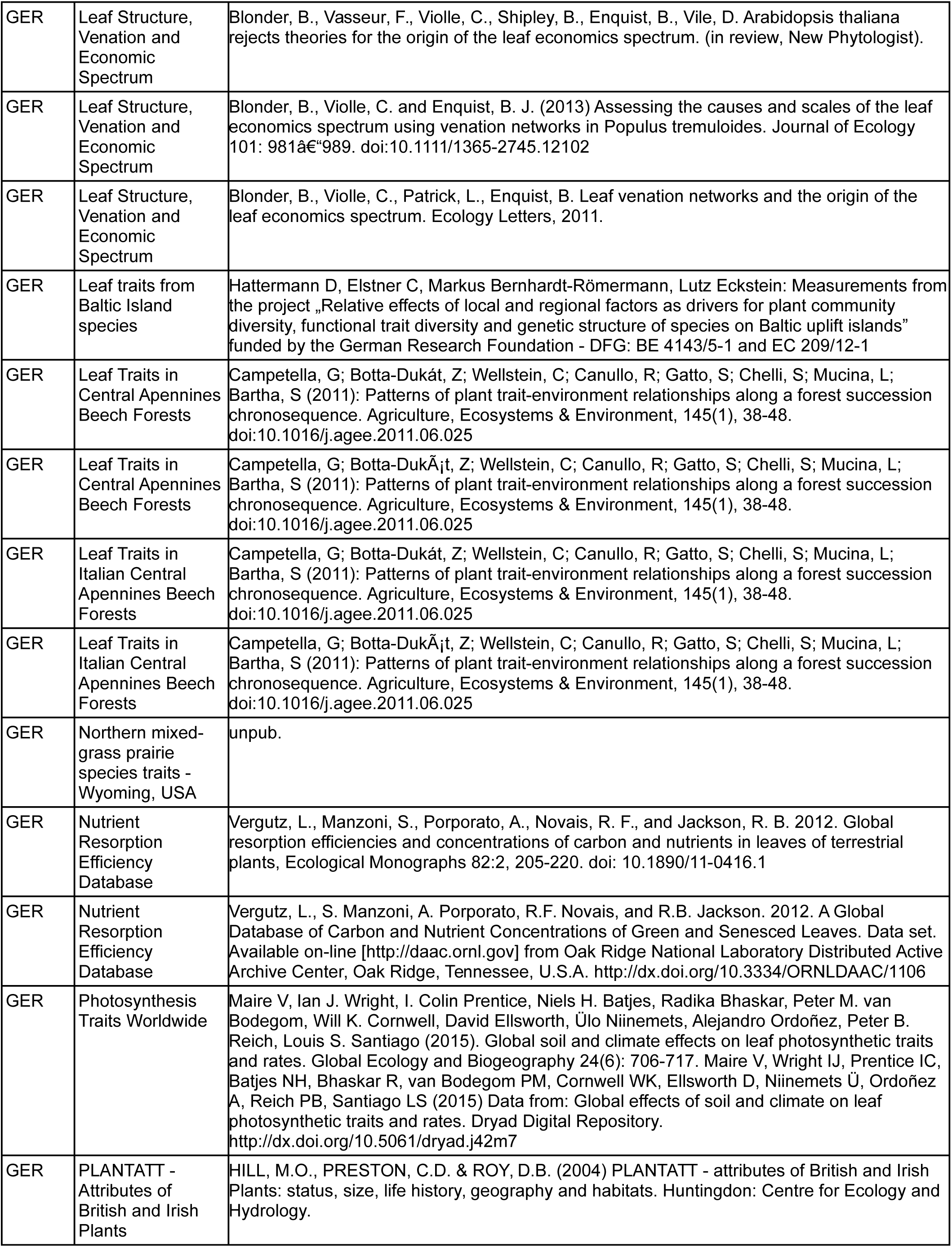
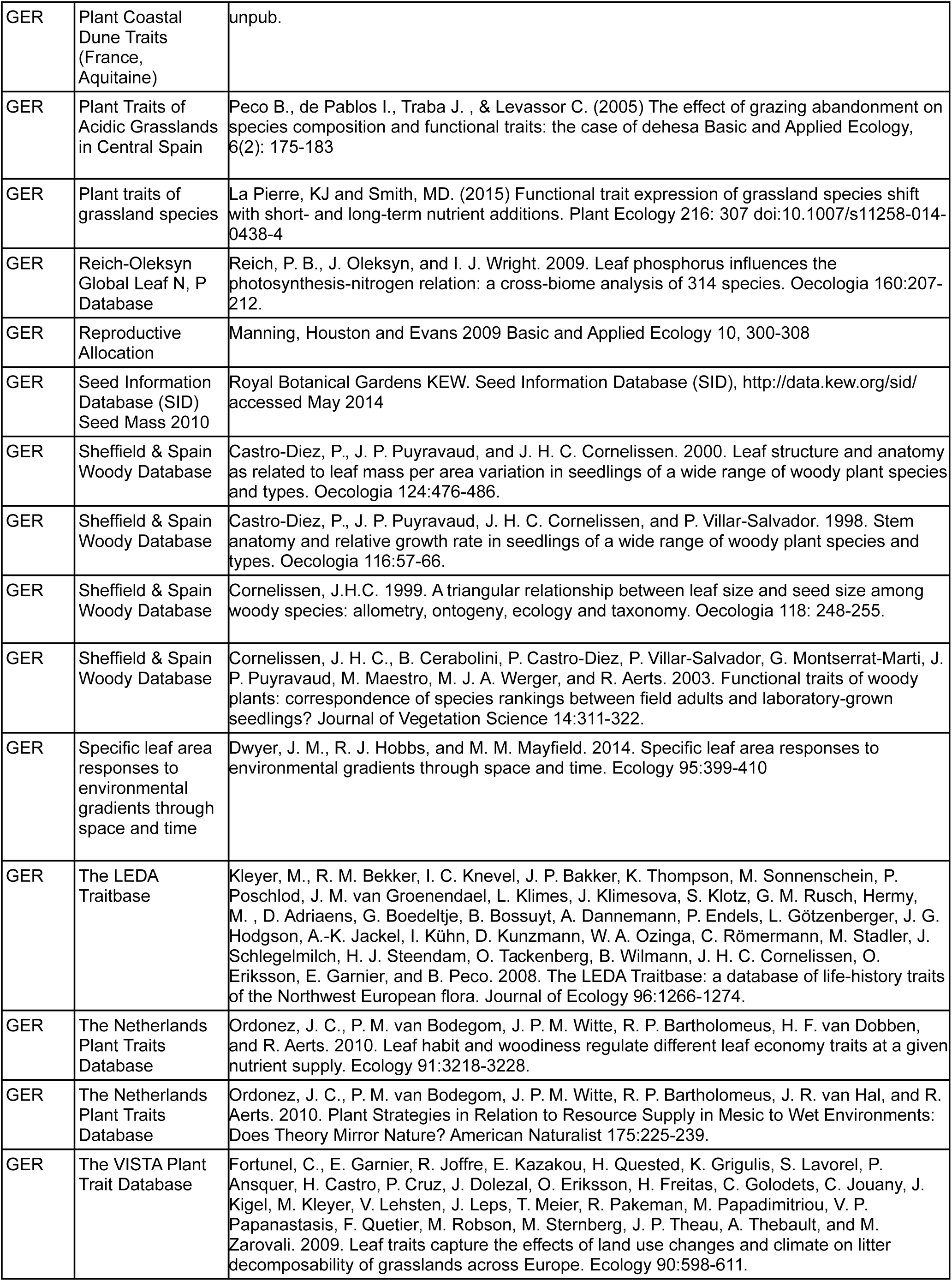
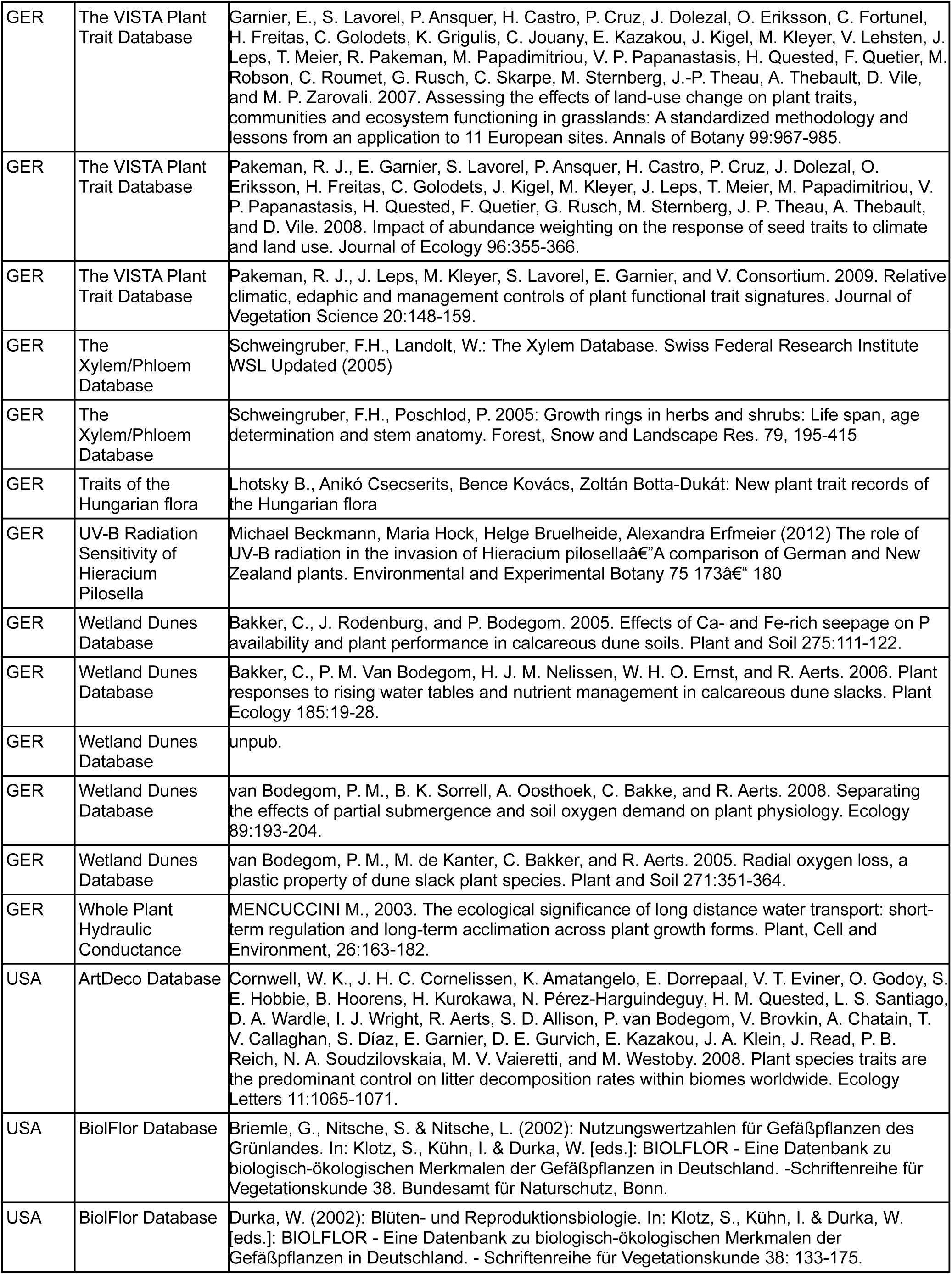
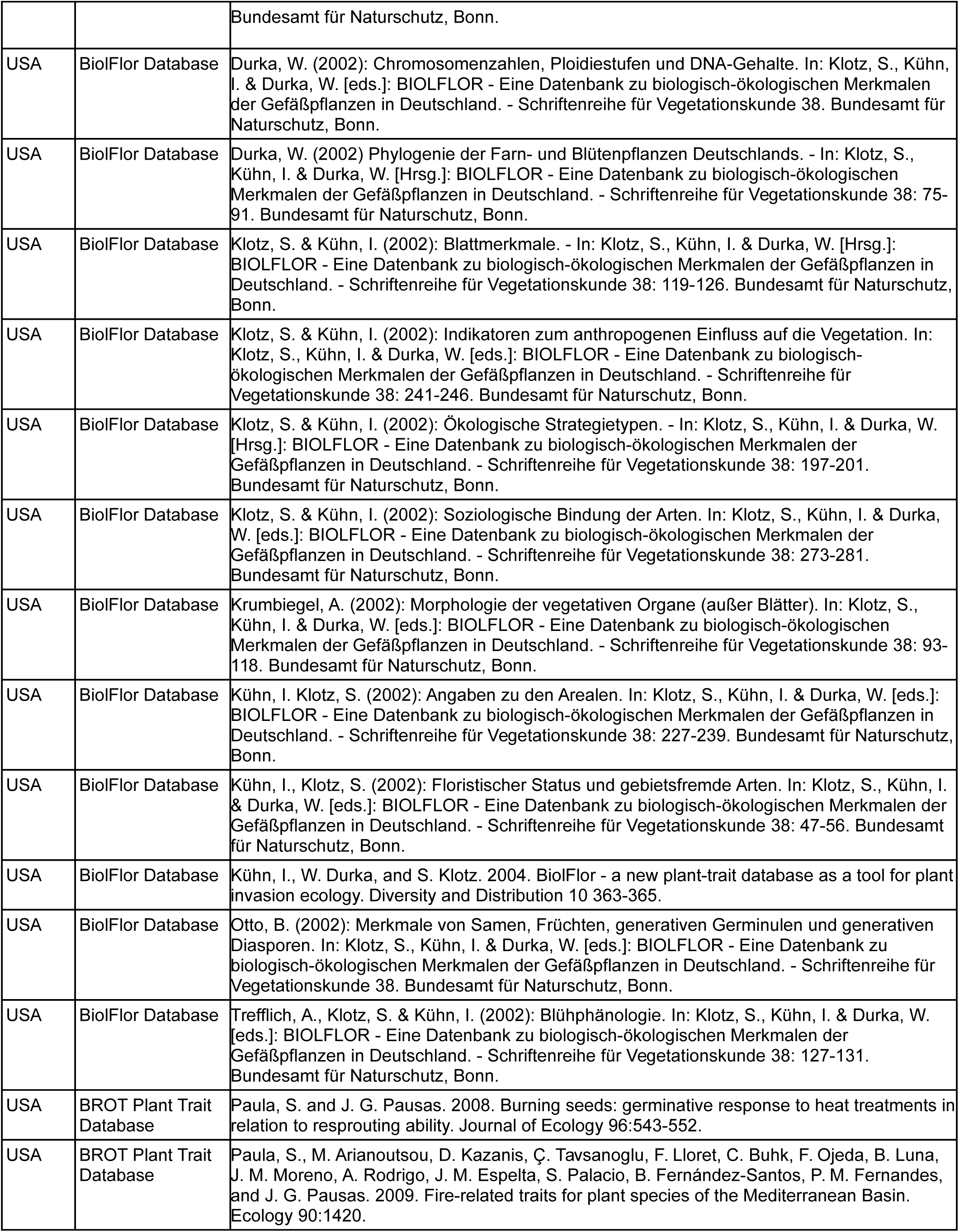
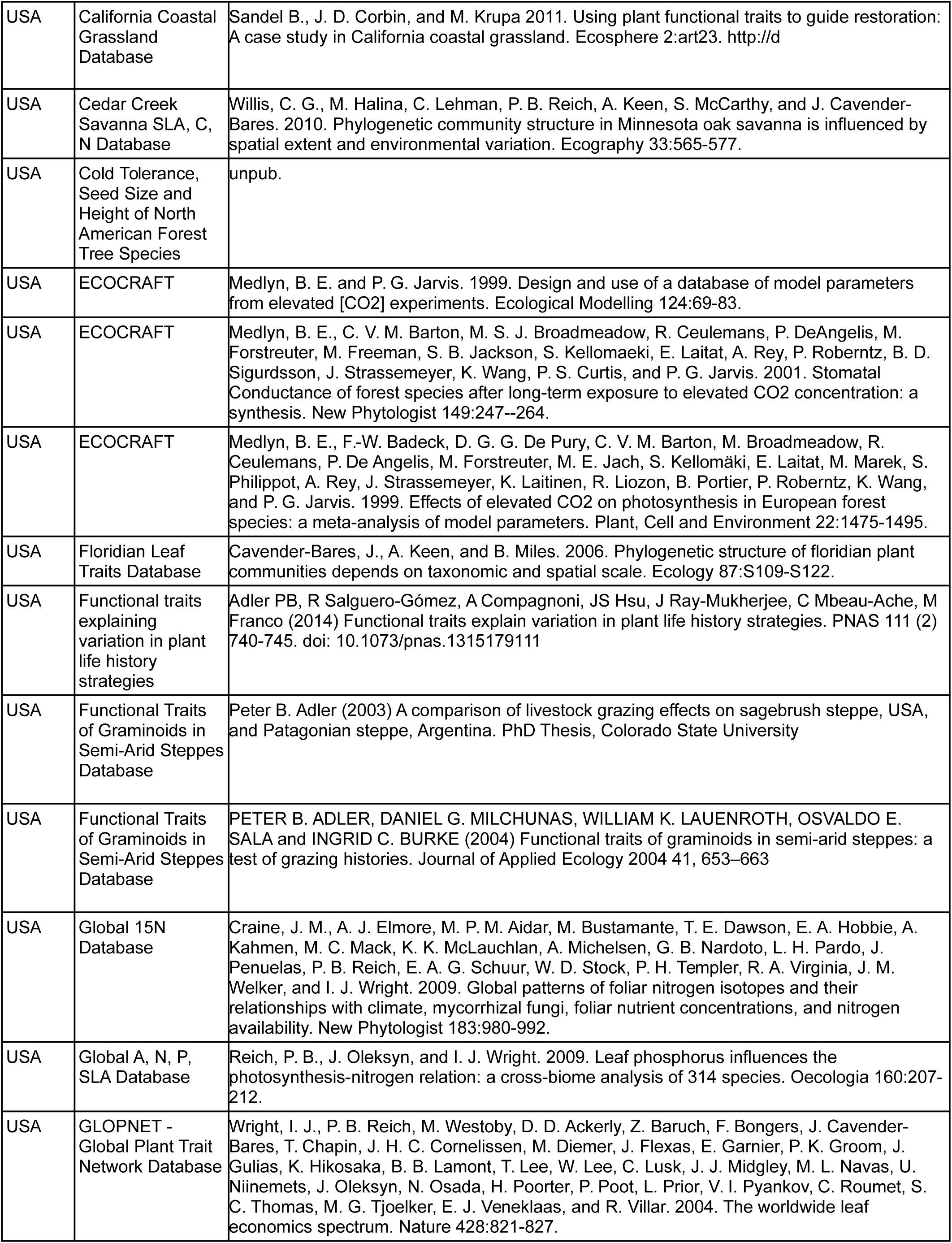
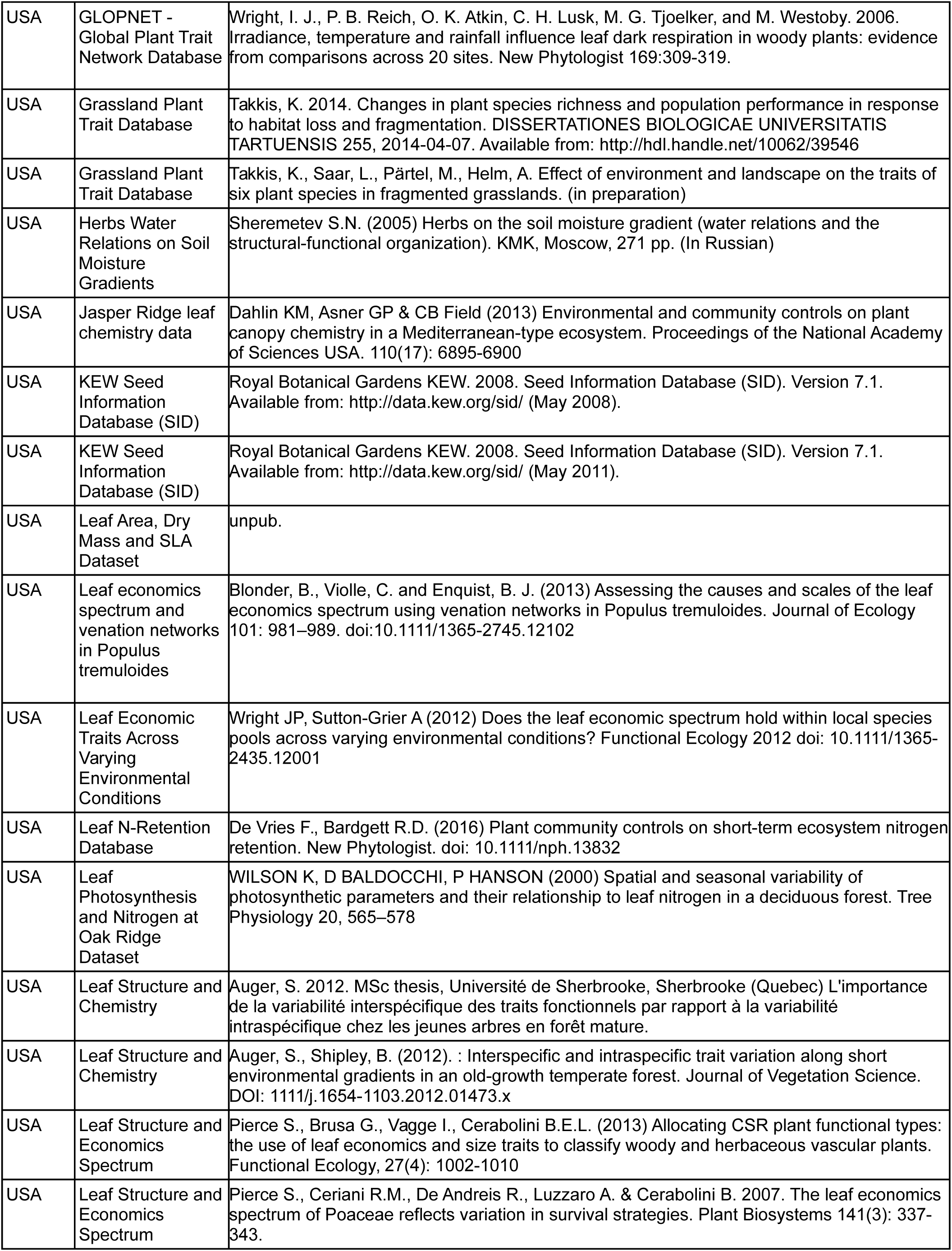
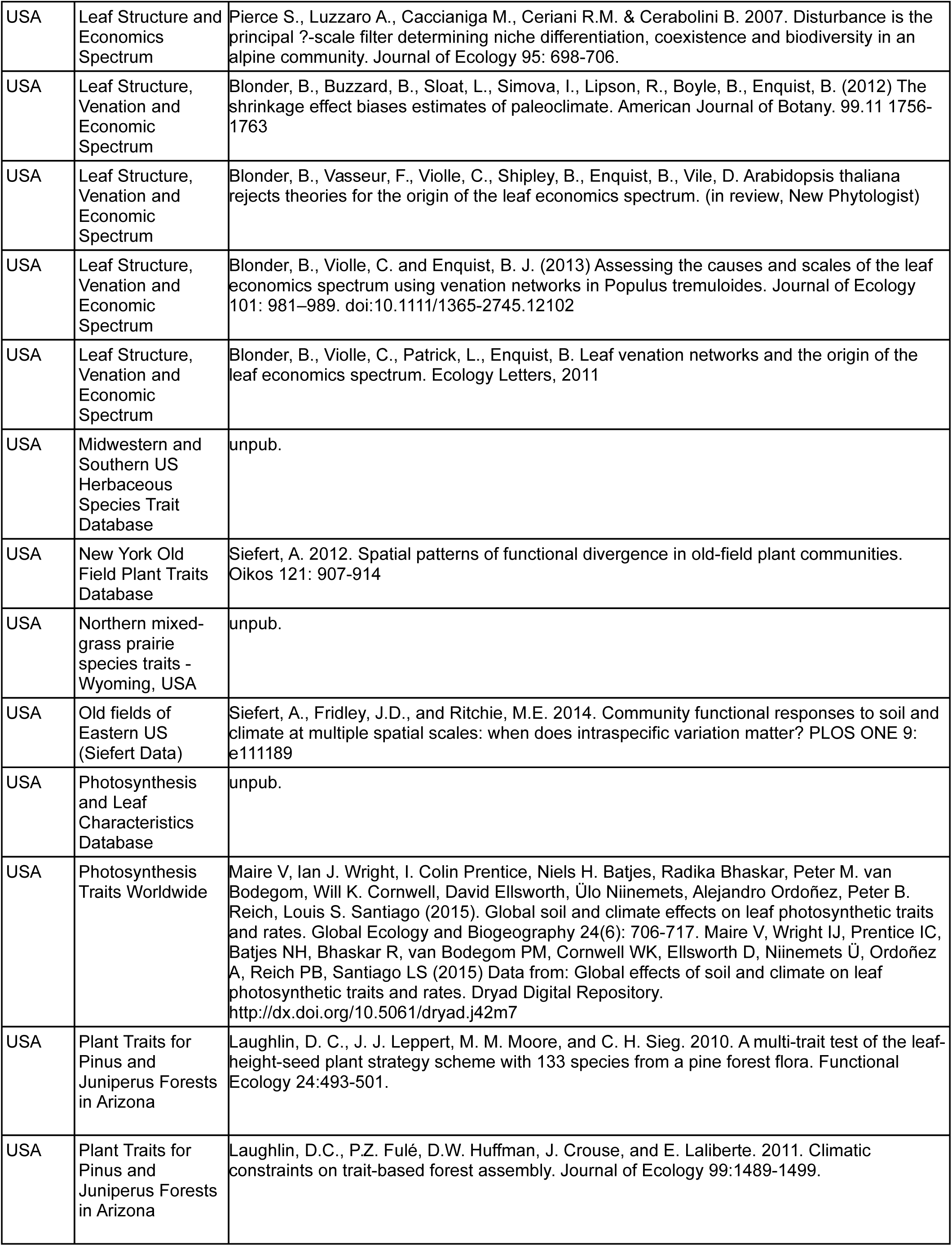
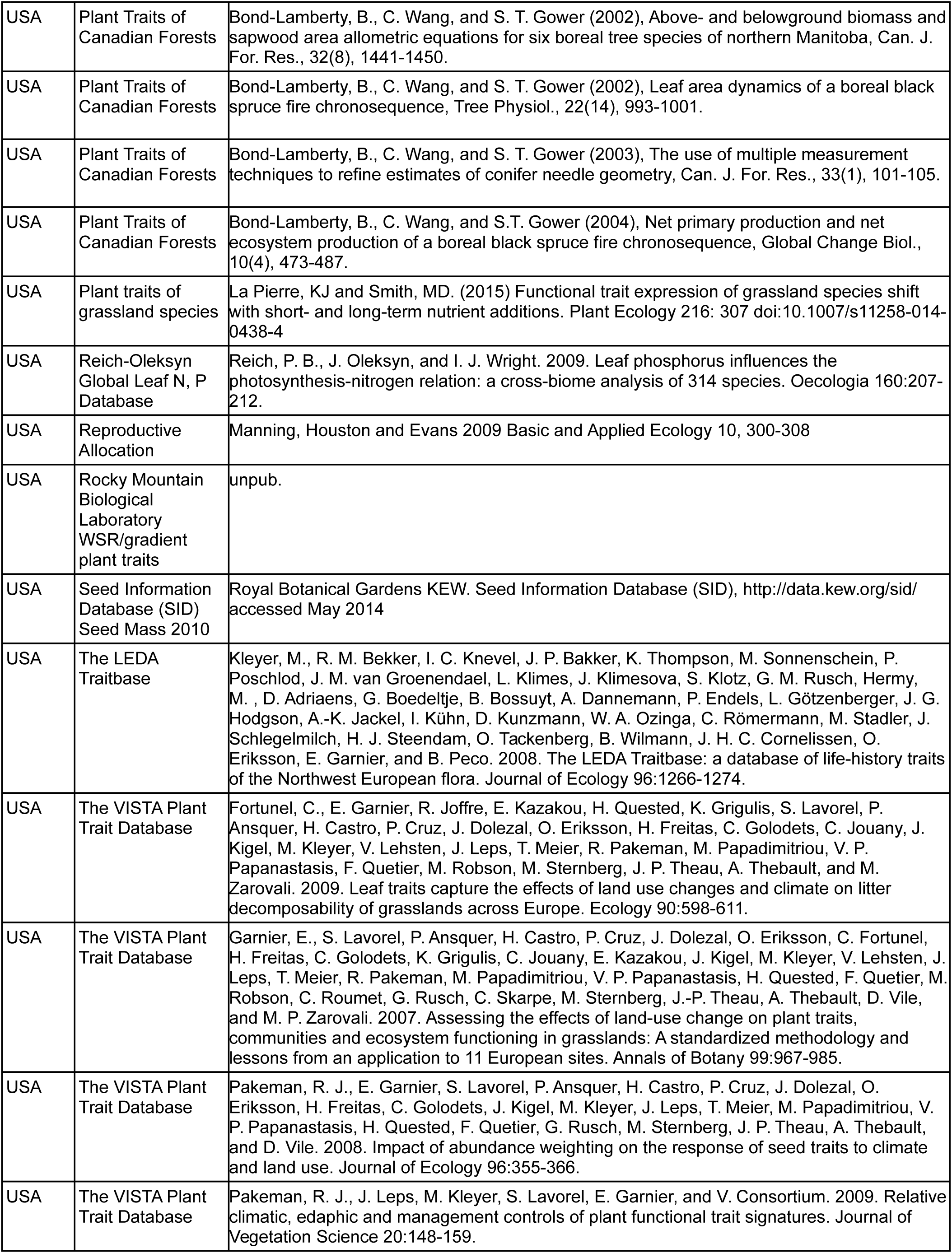
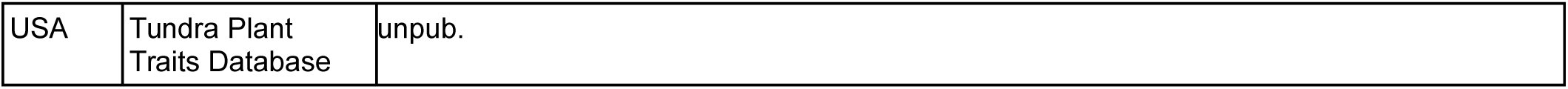
TRY references for plant species trait data from TRY^25^ requests 2968 and 4106. Data sources are sorted by the region their trait data have been used for (Germany=GER or USA). Note that, as mentioned in the main text, trait data for the USA dataset have been complemented by data from Cedar Creek plant trait assessments by Jane Catford, Peter Reich and Jeannine Cavender-Bares.

**Table S16.**
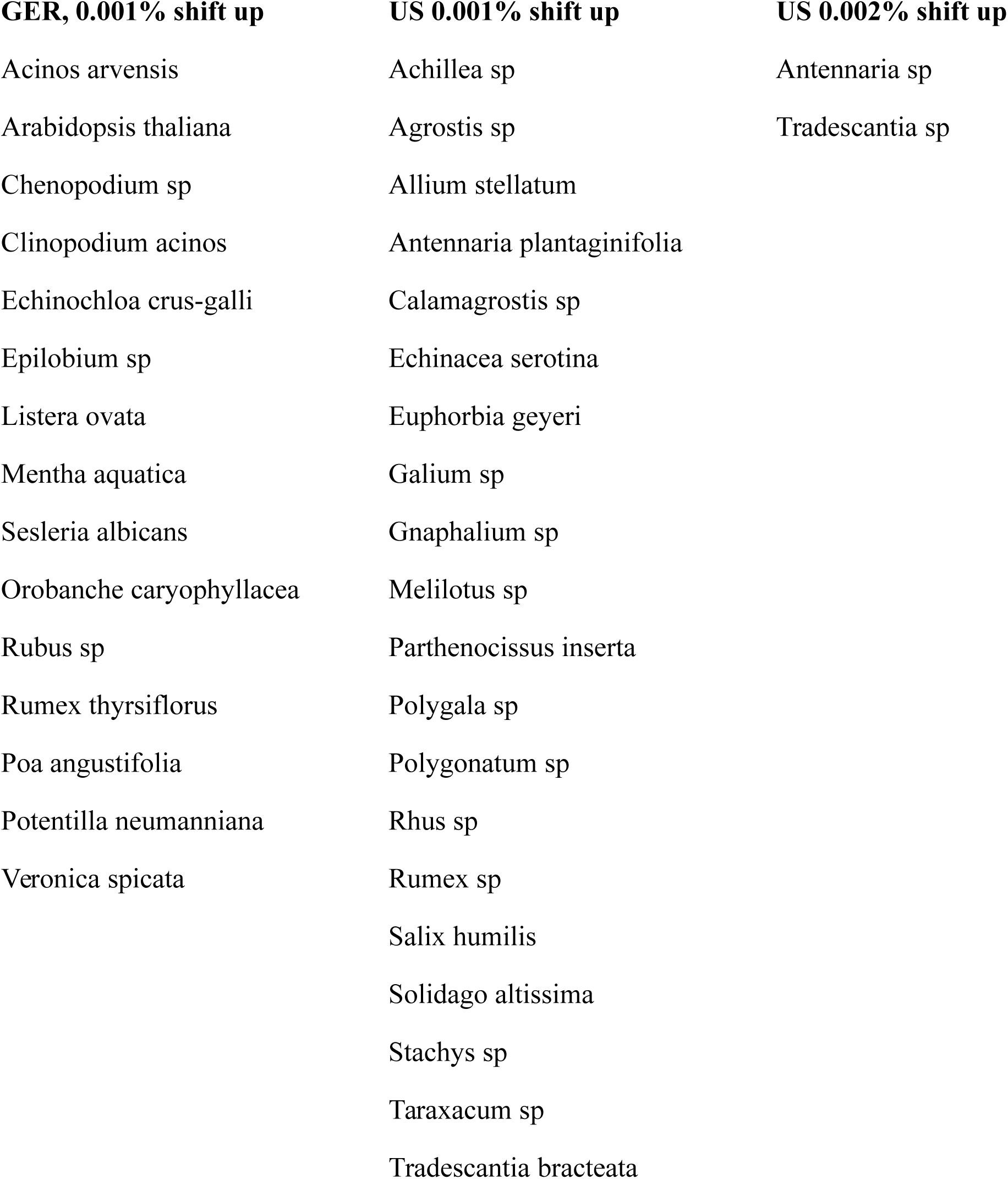
Species with altered trait values to avoid Gower dissimilarity zeros. Species are sorted by region (GER=Germany, US=USA) and by the percentage shift that their trait values were subject to. In two cases in the US dataset, there were three same-genus species with identical trait values and here two of them needed different shifts in order to obtain non-zero Gower dissimilarity values.

| GER, 0.001% shift up | US 0.001% shift up | US 0.002% shift up |
| --- | --- | --- |
| Acinos arvensis | Achillea sp | Antennaria sp |
| Arabidopsis thaliana | Agrostis sp | Tradescantia sp |
| Chenopodium sp | Allium stellatum |  |
| Clinopodium acinos | Antennaria plantaginifolia |  |
| Echinochloa crus-galli | Calamagrostis sp |  |
| Epilobium sp | Echinacea serotina |  |
| Listera ovata | Euphorbia geyeri |  |
| Mentha aquatica | Galium sp |  |
| Sesleria albicans | Gnaphalium sp |  |
| Orobanche caryophyllacea | Melilotus sp |  |
| Rubus sp | Parthenocissus inserta |  |
| Rumex thyrsiflorus | Polygala sp |  |
| Poa angustifolia | Polygonatum sp |  |
| Potentilla neumanniana | Rhus sp |  |
| Veronica spicata | Rumex sp |  |
|  | Salix humilis |  |
|  | Solidago altissima |  |
|  | Stachys sp |  |
|  | Taraxacum sp |  |
|  | Tradescantia bracteata |  |

**Table S17.**
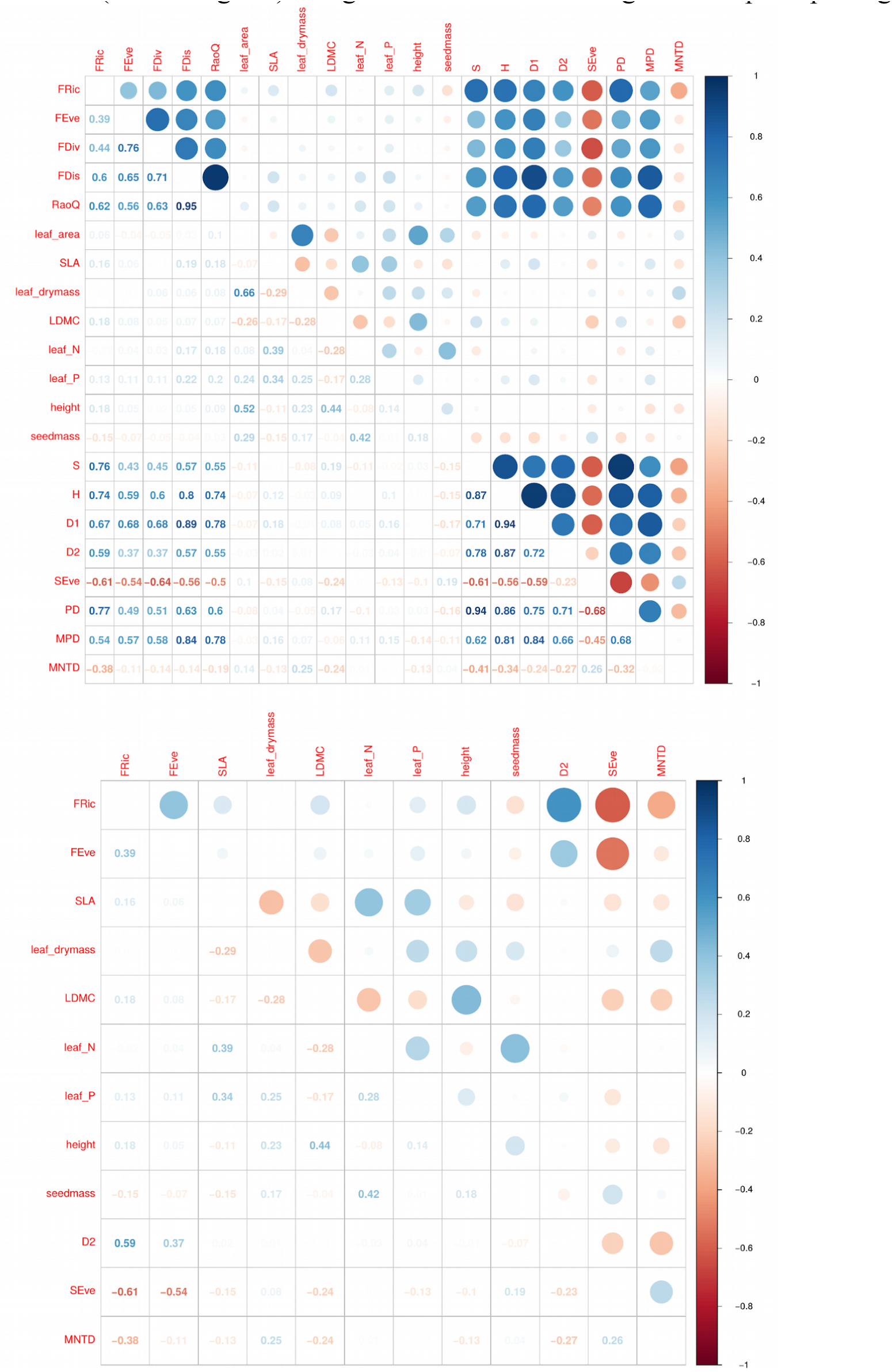
Correlation coefficients for 21 plant community properties for the German dataset. Pearson correlation coefficients and color code (see legend) for all 21 properties (upper diagram) and the subset of 12 community properties retained after stepwise removal due to variance inflation factors above 3 (lower diagram). Diagrams were created using the “corrplot” package^26^ in R.

**Table S18.**
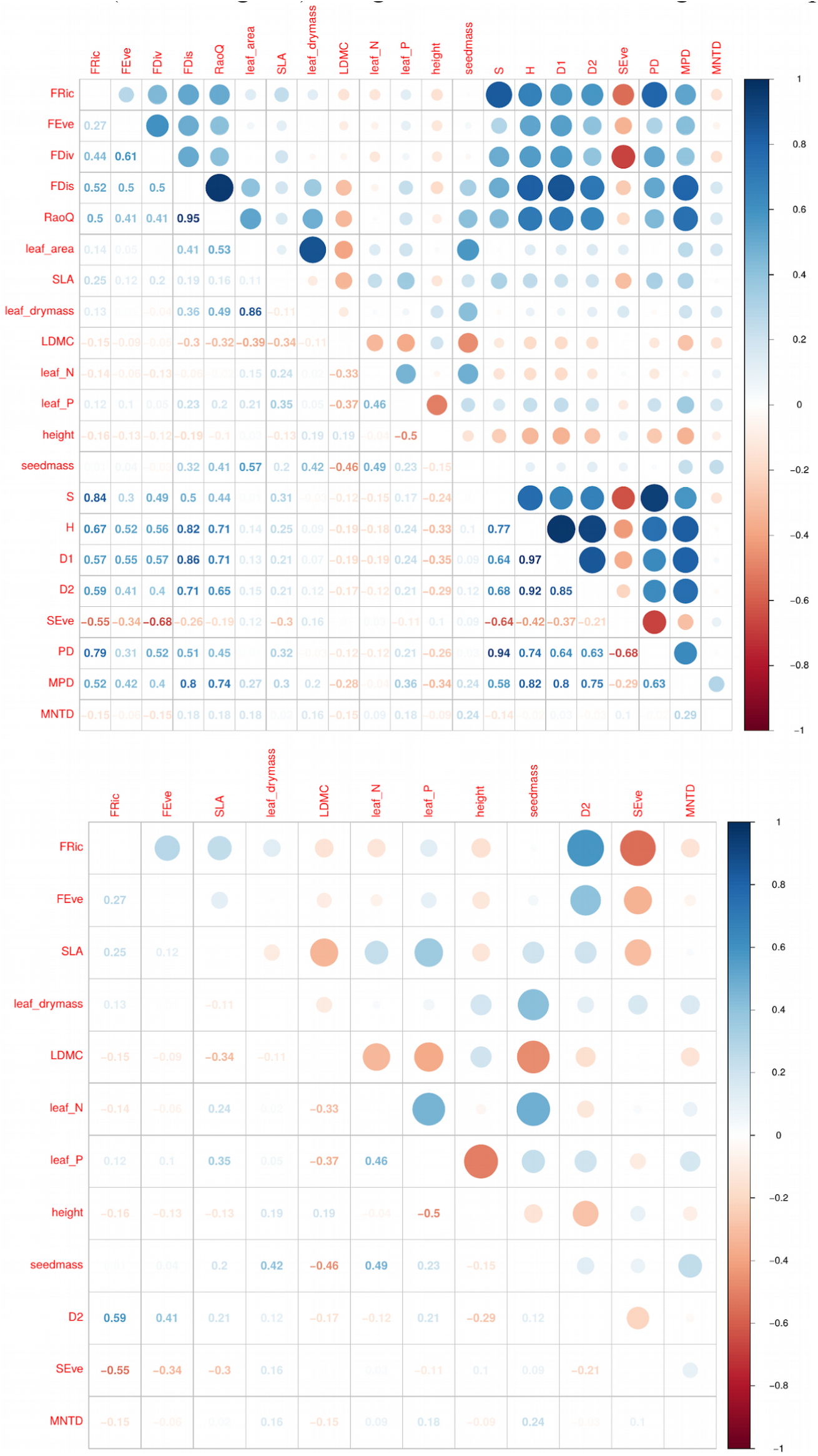
Correlation coefficients for 21 plant community properties for the US dataset. Pearson correlation coefficients and color code (see legend) for all 21 properties (upper diagram) and the subset of 12 community properties retained after stepwise removal due to variance inflation factors above 3 (lower diagram). Diagrams were created using the “corrplot” package^26^ in R.

## References

1. Cardinale, B. J. et al. Biodiversity loss and its impact on humanity. Nature 486, 59–67 (2012).

2. Tilman, D., Isbell, F. & Cowles, J. M. Biodiversity and ecosystem functioning. Annu. Rev. Ecol. Evol. Syst. 45, 471–493 (2014).

3. Isbell, F. et al. Linking the influence and dependence of people on biodiversity across scales. Nature 546, 65–72 (2017).

4. van der Plas, F. Biodiversity and ecosystem functioning in naturally assembled communities. Biol. Rev. online-early, (2019).

5. Tilman, D. et al. Diversity and productivity in a long-term grassland experiment. Science 294, 843–845 (2001).

6. Roscher, C., Schumacher, J. & Baade, J. The role of biodiversity for element cycling and trophic interactions: an experimental approach in a grassland community. Basic Appl. Ecol. 121, 107– 121 (2004).

7. Hooper, D. U. et al. Effects of biodiversity on ecosystem functioning: a consensus of current knowledge. Ecol. Monogr. 75, 3–35 (2005).

8. Cardinale, B. J. et al. The functional role of producer diversity in ecosystems. Am. J. Bot. 98, 572–592 (2011).

9. O’Connor, M. I. et al. A general biodiversity–function relationship is mediated by trophic level. Oikos 126, 18–31 (2017).

10. Huston, M. A. Hidden treatments in ecological experiments: re-evaluating the ecosystem function of biodiversity. Oecologia 110, 449–460 (1997).

11. Grime, J. P. Benefits of plant diversity to ecosystems: Immediate, filter and founder effects. J. Ecol. 86, 902–910 (1998).

12. Wardle, D. A. et al. Biodiversity and ecosystem function: an issue in ecology. Bull. Ecol. Soc. Am. 81, 235–239 (2000).

13. Leps, J. What do the biodiversity experiments tell us about consequences of plant species loss in the real world? Basic Appl. Ecol. 5, 529–534 (2004).

14. Srivastava, D. S. & Vellend, M. Biodiversity-ecosystem function research: is it relevant to conservation? Annu. Rev. Ecol. Evol. Syst. 36, 267–294 (2005).

15. Duffy, J. E. Why biodiversity is important to the functioning of real-world ecosystems. Front. Ecol. Environ. 7, 437–444 (2008).

16. Wardle, D. A. & Jonsson, M. Biodiversity effects in real ecosystems – a response to Duffy. Front. Ecol. Environ. 8, 10–11 (2010).

17. Duffy, J. E. Biodiversity effects: trends and exceptions – a reply to Wardle and Jonsson. Front. Ecol. Environ. 8, 11–12 (2010).

18. Wardle, D. A. Do experiments exploring plant diversity-ecosystem functioning relationships inform how biodiversity loss impacts natural ecosystems? J. Veg. Sci. 27, 646–653 (2016).

19. Schulze, E.-D. & Mooney, H. Biodiversity and Ecosystem Functioning. (Springer, 1993).

20. Naeem, S., Thompson, L. J., Lawler, S. P., Lawton, J. H. & Woodfin, R. M. Declining biodiversity can alter the performance of ecosystems. Nature 368, 734–737 (1994).

21. Balvanera, P. et al. Quantifying the evidence for biodiversity effects on ecosystem functioning and services. Ecol. Lett. 9, 1146–56 (2006).

22. Tilman, D., Wedin, D. & Knops, J. Productivity and sustainability influenced by biodiversity in grassland ecosystems. Nature 379, 718–720 (1996).

23. Loreau, M. et al. Biodiversity and Ecosystem Functioning: Current Knowledge and Future Challenges. Science 294, 804–808 (2001).

24. Reich, P. B. et al. Impacts of biodiversity loss escalate through time as redundancy fades. Science 336, 589–592 (2012).

25. Wilsey, B. J. & Potvin, C. Biodiversity and ecosystem functioning: Importance of species evenness in an old field. Ecology 81, 887–892 (2000).

26. Wilsey, B. J. & Polley, H. W. Realistically Low Species Evenness Does Not Alter Grassland Species-Richness–Productivity Relationships. Ecology 85, 2693–2700 (2004).

27. Hillebrand, H., Bennett, D. & Cadotte, M. Consequences of dominance: A Review of evenness effects on local and regional ecosystem processes. Ecology 89, 1510–1520 (2008).

28. Schmitz, M. et al. Consistent Effects of Biodiversity on Ecosystem Functioning Under Varying Density and Evenness. Folia Geobot. 48, 335–353 (2013).

29. Finn, J. A. et al. Ecosystem function enhanced by combining four functional types of plant species in intensively managed grassland mixtures: A 3-year continental-scale field experiment. J. Appl. Ecol. 50, 365–375 (2013).

30. Weisser, W. W. et al. Biodiversity effects on ecosystem functioning in a 15-year grassland experiment: patterns, mechanisms, and open questions. Basic Appl. Ecol. 23, 1–73 (2017).

31. Schmid, B. & Hector, A. The value of biodiversity experiments. Basic Appl. Ecol. 5, 535–542 (2004).

32. Eisenhauer, N. et al. Biodiversity-ecosystem function experiments reveal the mechanisms underlying the consequences of biodiversity change in real world ecosystems. J. Veg. Sci. 27, 1061–1070 (2016).

33. Isbell, F. et al. Nutrient enrichment, biodiversity loss, and consequent declines in ecosystem productivity. Proc. Natl. Acad. Sci. 110, 11911–11916 (2013).

34. Duffy, J. E., Godwin, C. M. & Cardinale, B. J. Biodiversity effects in the wild are common and as strong as key drivers of productivity. Nature 549, 261–264 (2017).

35. Buchmann, T. et al. Connecting experimental biodiversity research to real-world grasslands. Perspect. Plant Ecol. Evol. Syst. 33, 78–88 (2018).

36. Tilman, D. et al. The influence of functional diversity and composition on ecosystem processes. Science 277, 1300–1302 (1997).

37. Tilman, D., Reich, P. B. & Isbell, F. Biodiversity impacts ecosystem productivity as much as resources, disturbance, or herbivory. Proc. Natl. Acad. Sci. 109, 10394–10397 (2012).

38. Isbell, F. et al. Biodiversity increases the resistance of ecosystem productivity to climate extremes. Nature 526, 574–577 (2015).

39. Fischer, M. et al. Implementing large-scale and long-term functional biodiversity research: The Biodiversity Exploratories. Basic Appl. Ecol. 11, 473–485 (2010).

40. Soliveres, S. et al. Biodiversity at multiple trophic levels is needed for ecosystem multifunctionality. Nature 536, 456–459 (2016).

41. Tilman, D. Secondary succession and the pattern of plant dominance along experimental nitrogen gradients. Ecol. Monogr. 57, 189–214 (1987).

42. Clark, C. M. & Tilman, D. Loss of plant species after chronic low-level nitrogen deposition to prairie grasslands. Nature 451, 712–715 (2008).

43. Inouye, R. et al. Old-field succession on a Minnesota sand plain. Ecology 68, 12–26 (1987).

44. Díaz, S. et al. The global spectrum of plant form and function. Nature 529, 1–17 (2015).

45. Craven, D. et al. Multiple facets of biodiversity drive the diversity–stability relationship. *Nat*. Ecol. Evol. 2, 1579–1587 (2018).

46. Manning, P. et al. Grassland management intensification weakens the associations among the diversities of multiple plant and animal taxa. Ecology 96, 1492–1501 (2015).

47. Loreau, M. & Hector, A. Partitioning selection and complementarity in biodiversity experiments. Nature 412, 72–76 (2001).

48. Hillebrand, H. & Matthiessen, B. Biodiversity in a complex world: consolidation and progress in functional biodiversity research. Ecol. Lett. 12, 1405–1419 (2009).

49. Liang, J. et al. Positive biodiversity-productivity relationship predominant in global forests. Science 354, aaf8957 (2016).

50. Grace, J. B. et al. Integrative modelling reveals mechanisms linking productivity and plant species richness. Nature 529, 390–393 (2016).

51. Oehri, J., Schmid, B., Schaepman-Strub, G. & Niklaus, P. A. Biodiversity promotes primary productivity and growing season lengthening at the landscape scale. Proc. Natl. Acad. Sci. 114, 10160–10165 (2017).

52. Díaz, S. et al. Incorporating plant functional diversity effects in ecosystem service assessments. Proc. Natl. Acad. Sci. U. S. A. 104, 20684–20689 (2007).

53. Lavorel, S. et al. Using plant functional traits to understand the landscape distribution of multiple ecosystem services. J. Ecol. 99, 135–147 (2011).

54. Schmid, B. The species richness-productivity controversy. Trends Ecol. Evol. 17, 113–114 (2002).

55. Maestre, F. T. et al. Plant species richness and ecosystem multifunctionality in global drylands. Science 335, 214–218 (2012).

56. Allan, E. et al. Land use intensification alters ecosystem multifunctionality via loss of biodiversity and changes to functional composition. Ecol. Lett. 18, 834–843 (2015).

57. van der Plas, F. et al. Jack-of-all-trades effects drive biodiversity-ecosystem multifunctionality relationships in European forests. Nat. Commun. 7, 11109 (2016).

58. Hobbs, R. J., Higgs, E. & Harris, J. a. Novel ecosystems: implications for conservation and restoration. Trends Ecol. Evol. 24, 599–605 (2009).

59. Roscher, C. et al. Convergent high diversity in naturally colonized experimental grasslands is not related to increased productivity. Perspect. Plant Ecol. Evol. Syst. 20, 32–45 (2016).

60. Ellenberg, H. & Leuschner, C. Vegetation Mitteleuropas mit den Alpen: in ökologischer, dynamischer und historischer Sicht. (UTB, 2010).

61. Blüthgen, N. et al. A quantitative index of land-use intensity in grasslands: Integrating mowing, grazing and fertilization. Basic Appl. Ecol. 13, 207–220 (2012).

62. Tilman, D., Reich, P. B. & Knops, J. M. H. Biodiversity and ecosystem stability in a decade-long grassland experiment. Nature 441, 629–632 (2006).

63. Tilman, D. Community Invasibility, Recruitment Limitation, and Grassland Biodiversity. Ecology 78, 81–92 (1997).

64. Catford, J. A. et al. Traits linked with species invasiveness and community invasibility vary with time, stage and indicator of invasion in a long-term grassland experiment. Ecol. Lett. 22, 593– 604 (2019).

65. Fargione, J. et al. From selection to complementarity: shifts in the causes of biodiversity-productivity relationships in a long-term biodiversity experiment. Proc. R. Soc. B Biol. Sci. 274, 871–876 (2007).

66. Londo, G. The decimal scale for releves of permanent quadrats. Vegetatio 33, 61–64 (1976).

67. Roscher, C. et al. What happens to the sown species if a biodiversity experiment is not weeded? Basic Appl. Ecol. 14, 187–198 (2013).

68. Kattge, J. et al. TRY - a global database of plant traits. Glob. Chang. Biol. 17, 2905–2935 (2011).

69. Cayuela, L., Stein, A. & Oksanen, J. Taxonstand: Taxonomic Standardization of Plant Species Names. R Packag. version 2.*1* (2017).

70. The Plant List, Version 1.1. (2013). Available at: http://www.theplantlist.org/.

71. Qian, H. & Jin, Y. An updated megaphylogeny of plants, a tool for generating plant phylogenies and an analysis of phylogenetic community structure. *J*. Plant Ecol. 9, 233–239 (2016).

72. Martins, W. S., Carmo, W. C., Longo, H. J., Rosa, T. C. & Rangel, T. F. SUNPLIN: Simulation with Uncertainty for Phylogenetic Investigations. BMC Bioinformatics 14, (2013).

73. Rangel, T. F. et al. Phylogenetic uncertainty revisited: Implications for ecological analyses. Evolution (N. Y). 69, 1301–1312 (2015).

74. Goolsby, E. W., Bruggeman, J. & Ane, C. Rphylopars: Phylogenetic Comparative Tools for Missing Data and Within-Species Variation. (2016).

75. Penone, C. et al. Imputation of missing data in life-history trait datasets: Which approach performs the best? Methods Ecol. Evol. 5, 1–10 (2014).

76. Oksanen, J. et al. Vegan: community ecology package. R Packag. version 2.3–4 (2016).

77. Magurran, A. Measuring Biological Diversity. (Blackwell, 2004).

78. Morris, E. K. et al. Choosing and using diversity indices: Insights for ecological applications from the German Biodiversity Exploratories. Ecol. Evol. 4, 3514–3524 (2014).

79. Smith, B. & Wilson, J. B. A Consumer’s Guide to Evenness Indices. Oikos 76, 70–82 (1996).

80. Hill, M. Diversity and Evenness: A Unifying Notation and Its Consequences. Ecology 54, 427– 432 (1973).

81. Tucker, C. M. et al. A guide to phylogenetic metrics for conservation, community ecology and macroecology. Biol. Rev. 92, 698–715 (2017).

82. Kembel, S. W. et al. Picante: R tools for integrating phylogenies and ecology. Bioinformatics 26, 1463–1464 (2010).

83. Villéger, S., Mason, N. W. H. & Mouillot, D. New multidimensional functional diversity indices for a multifaceted framework in functional ecology. Ecology 89, 2290–2301 (2008).

84. Laliberte, E. & Legendre, P. A distance-based framework for measuring functional diversity from multiple traits A distance-based framework for measuring from multiple traits functional diversity. Ecology 91, 299–305 (2010).

85. Mouchet, M. A., Villéger, S., Mason, N. W. H. & Mouillot, D. Functional diversity measures: an overview of their redundancy and their ability to discriminate community assembly rules. Funct. Ecol. 24, 867–876 (2010).

86. Laliberté, E., Legendre, P. & Shipley., B. FD: measuring functional diversity from multiple traits, and other tools for functional ecology. (2014).

87. R Core Team. R: A Language and Environment for Statistical Computing. (2017). doi:10.1007/978-3-540-74686-7

88. Zuur, A. F., Ieno, E. N. & Elphick, C. S. A protocol for data exploration to avoid common statistical problems. Methods Ecol. Evol. 1, 3–14 (2010).

89. Fox, J. & Weisberg, S. An R Companion to Applied Regression. (Thousand Oaks, 2011).

90. Pebesma, E. & Bivand, R. Classes and methods for spatial data. R News 5, 9–13 (2005).

91. Bivand, R. S., Pebesma, E. & Gomez-Rubio, V. Applied spatial data analysis with R. (Springer, 2013).

92. Habel, K., Grasman, R., Gramacy, R. B., Stahel, A. & Sterratt, D. C. geometry: Mesh Generation and Surface Tesselation. R package version 0.4.1 (2019).

93. Blonder, B. & Harris, D. hypervolume: High Dimensional Geometry and Set Operations Using Kernel Density Estimation, Support Vector Machines, and Convex Hulls. R Packag. ver. 2.0.11 (2018).

94. Meyer, S. T. et al. Effects of biodiversity strengthen over time as ecosystem functioning declines at low and increases at high biodiversity. Ecosphere 7, e01619 (2016).

## Supporting Information References

1. Fornara, D. A. & Tilman, D. Plant functional composition influences rates of soil carbon and nitrogen accumulation. J. Ecol. 96, 314–322 (2008).

2. Lange, M. et al. Plant diversity increases soil microbial activity and soil carbon storage. Nat. Commun. 6, 6707 (2015).

3. Ravenek, J. M. et al. Long-term study of root biomass in a biodiversity experiment reveals shifts in diversity effects over time. Oikos 123, 1528–1536 (2014).

4. Meyer, S. T. et al. Consistent increase in herbivory along two experimental plant diversity gradients over multiple years. Ecosphere 8, e01876 (2017).

5. Scheu, S. Automated measurement of the respiratory response of soil microcompartments: Active microbial biomass in earthworm faeces. Soil Biol. Biochem. 24, 1113–1118 (1992).

6. Eisenhauer, N. et al. Plant diversity effects on soil microorganisms support the singular hypothesis. Ecology 91, 485–496 (2010).

7. Strecker, T., Mace, O. G., Scheu, S. & Eisenhauer, N. Functional composition of plant communitites determines the spatial and temporal stability of soil microbial properties in a long-term plant diversity experiment. Oikos 125, 1743–1754 (2016).

8. Beck, T. et al. An inter-laboratory comparison of ten different ways of measuring soil microbial biomass C. Soil Biol. Biochem. 29, 1023–1032 (1997).

9. Hacker, N. et al. Plant diversity shapes microbe-rhizosphere effects on P mobilisation from organic matter in soil. Ecol. Lett. 18, 1356–1365 (2015).

10. Eivazi, F. & Tabatabai, M. A. Phosphatases in soils. Soil Biol. Biochem. 9, 167–172 (1977).

11. Catford, J. A. et al. Traits linked with species invasiveness and community invasibility vary with time, stage and indicator of invasion in a long-term grassland experiment. Ecol. Lett. 22, 593– 604 (2019).

12. Inouye, R. et al. Old-field succession on a Minnesota sand plain. Ecology 68, 12–26 (1987).

13. Tilman, D. Community Invasibility, Recruitment Limitation, and Grassland Biodiversity. Ecology 78, 81–92 (1997).

14. Roscher, C., Schumacher, J. & Baade, J. The role of biodiversity for element cycling and trophic interactions: an experimental approach in a grassland community. Basic Appl. Ecol. 121, 107– 121 (2004).

15. Fischer, M. et al. Implementing large-scale and long-term functional biodiversity research: The Biodiversity Exploratories. Basic Appl. Ecol. 11, 473–485 (2010).

16. Roscher, C. et al. Convergent high diversity in naturally colonized experimental grasslands is not related to increased productivity. Perspect. Plant Ecol. Evol. Syst. 20, 32–45 (2016).

17. Weisser, W. W. et al. Biodiversity effects on ecosystem functioning in a 15-year grassland experiment: patterns, mechanisms, and open questions. Basic Appl. Ecol. 23, 1–73 (2017).

18. Tilman, D. et al. The influence of functional diversity and composition on ecosystem processes. Science (80-.). 277, 1300–1302 (1997).

19. Tilman, D. Secondary succession and the pattern of plant dominance along experimental nitrogen gradients. Ecol. Monogr. 57, 189–214 (1987).

20. Adler, D. & Kelly, T. vioplot: violin plot. R. R Packag. version 0.3.0 (2018).

21. Chen, H. VennDiagram: Generate High-Resolution Venn and Euler Plots. R Packag. version 1.6.20 (2018).

22. Oksanen, J. et al. Vegan: community ecology package. R Packag. version 2.3-4 (2016).

23. Bocci, G. TR8: An R package for easily retrieving plant species traits. Methods Ecol. Evol. 6, 347–350 (2015).

24. Fitter, A. & Peat, H. The Ecological Flora Database. J. Ecol. 82, 415–425 (1994).

25. Kattge, J. et al. TRY - a global database of plant traits. Glob. Chang. Biol. 17, 2905–2935 (2011).

26. Wei, T. & Simko, V. R package ‘corrplot’: Visualization of a Correlation Matrix. R Packag. version 0.84 (2017).

